# A Brain Reward Circuit Inhibited By Next-Generation Weight Loss Drugs

**DOI:** 10.1101/2024.12.12.628169

**Authors:** Elizabeth N. Godschall, Taha Bugra Gungul, Isabelle R. Sajonia, Aleyna K. Buyukaksakal, Orien Dong-Ang Li, Sophia Ogilvie, Austin B. Keeler, Guilian Tian, Omar Koita, Yu Shi, Tyler C. J. Deutsch, Maisie Crook, YuChen Zhang, Nicholas J. Conley, Addison N. Webster, O. Yipkin Calhan, Weile Liu, Amani Akkoub, Karan Malik, Kaleigh I. West, Sara Michel-Le, Arun Karthikeyan, Grace van Gerven, Kevin T. Beier, Larry S. Zweifel, Manoj K. Patel, John N. Campbell, Christopher D. Deppmann, Ali D. Güler

**Affiliations:** Department of Biology, University of Virginia; Charlottesville, VA 22904, USA; Department of Neuroscience, School of Medicine, University of Virginia; Charlottesville, VA 22903, USA; Department of Anesthesiology, University of Virginia Health System, Charlottesville, VA 22904, USA; Program in Fundamental Neuroscience; Charlottesville, VA 22904, USA; Department of Cell Biology, University of Virginia; Charlottesville, VA 22904, USA; Department of Biomedical Engineering, University of Virginia; Charlottesville, VA 22904, USA; Department of Physiology and Biophysics, University of California, Irvine, Irvine, CA, 92617, USA; Department of Pharmacology, University of Washington, Seattle, WA, 98195, USA; Department of Psychiatry, University of Washington, Seattle, WA, 98195, USA

## Abstract

Glucagon-like peptide-1 receptor agonists (GLP1RAs) effectively reduce body weight and improve metabolic outcomes, yet established peptide-based therapies require injections and complex manufacturing. Small-molecule GLP1RAs promise oral bioavailability and scalable manufacturing, but their selective binding to human versus rodent receptors has limited mechanistic studies. Here, we developed humanized GLP1R mouse models to investigate how small-molecule GLP1RAs influence feeding behavior. This approach revealed that these compounds regulate both homeostatic and hedonic feeding through parallel neural circuits. Beyond engaging canonical hypothalamic and hindbrain networks that control metabolic homeostasis, GLP1RAs recruit a discrete population of Glp1r-expressing neurons in the central amygdala, which selectively suppress the consumption of palatable foods by reducing dopamine release in the nucleus accumbens. Stimulating these central amygdalar neurons curtail hedonic feeding, whereas targeted deletion of the receptor in this cell population specifically diminishes the anorectic efficacy of GLP1RAs for reward-driven intake. These findings reveal a dedicated neural circuit through which small molecule GLP1RAs modulate reward processing, suggesting broad therapeutic potential in conditions of dysregulated dopamine signaling including substance use disorder and binge eating.

## Main

Glucagon-like peptide-1 receptor agonists (GLP1RAs) have emerged as highly effective treatments for obesity and diabetes. Beyond their metabolic benefits, these drugs show promise for treating conditions such as substance use disorders, suggesting they act on central circuits governing reward and motivation^1–6,7–14,15–19^. While GLP1RAs are known to engage hypothalamic and hindbrain regions to suppress homeostatic feeding^20–27^, their impact on the neural circuits that drive hedonic food consumption and reward-related behaviors remains largely unknown. Understanding the underlying neural mechanisms is especially timely, as small-molecule GLP1RAs are now poised to expand access to this transformative class of medications^28–31^.

Next-generation GLP1RAs like danuglipron (PF06882961) and orforglipron (LY3502970) offer advantages through oral bioavailability and scalable manufacturing (**Fig. 1a**)^32,33–39^. Yet these compounds show species-specific binding properties and many small-molecule GLP1RAs that activate human GLP1R show minimal activity at rodent receptors, precluding preclinical investigation of their mechanisms of action^37,40,41,42^. Notably, orforglipron has advanced into over ten registered Phase 3 clinical trials^33–35,43^ while danuglipron’s development was discontinued due to side effects that broader preclinical profiling might have uncovered^44^.

**Figure 1:**
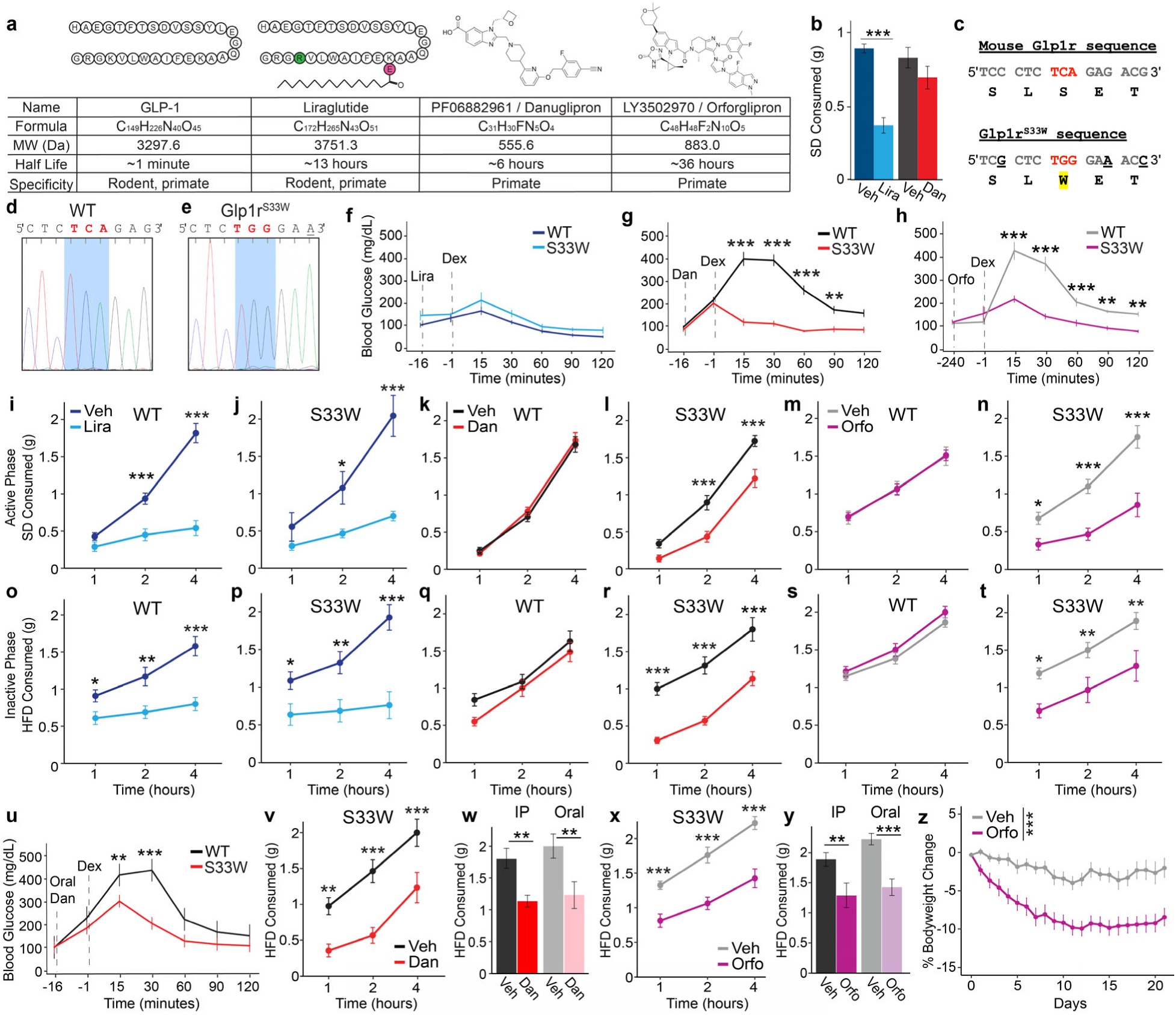
Generation and validation of a novel mouse model responsive to small molecule GLP1R agonists: **a**, Structure, naming convention, molecular formula, molecular weight (MW), half-life in humans, and species binding specificity of glucagon-like peptide-1 (GLP-1), liraglutide, danuglipron, and orforglipron. Amino acid sequences of liraglutide shown with its substitution and additions to GLP-1 highlighted in green and purple, respectively. **b**, Standard diet (SD) consumption over 2 hours post-administration of liraglutide (Lira), vehicle (Veh), or danuglipron (Dan) (*n* = 11 per injection, one-way ANOVA with Bonferroni correction, \*\*\**P*<0.001). **c**, Schematic of the serine (TCA or S) to tryptophan (TGG or W) CRISPR-mediated substitution in Glp1r^S33W^ (bottom) or in mouse Glp1r (top). **d,e**, Sanger sequencing chromatograms of (**d**) WT mouse Glp1r and (**e**) Glp1r^S33W^ sequence, confirming the Glp1r S33W substitution in mice. **f-h**, Glucose tolerance test (GTT) on GLP1RAs. Comparison of blood glucose levels on (**f**) liraglutide (Lira), (**g**) danuglipron (Dan), (**h**) orforglipron (Orfo) followed by dextrose (Dex) in WT and Glp1r^S33W^ mice (*n* = 5-9 per injection, two-way ANOVA with Bonferroni correction, \*\**P*<0.01; \*\*\**P*<0.001). **i-n**, SD intake 1, 2, and 4 hours post-treatment of (**i, j**) Lira (*n* = 9-11), (**k, l**) Dan (*n* = 15-16), and (**m, n**) Orfo (*n* = 8-9) and vehicle controls in WT and Glp1r^S33W^ mice. **o-t**, High fat diet (HFD) intake 1, 2, and 4 hours post-treatment of (**o, p**) Lira (*n* = 8-10), (**q, r**) Dan (*n* = 14-16), and (**s, t**) Orfo (*n* = 8-9) and their vehicle controls in WT and Glp1r^S33W^ mice (two-way ANOVA with Bonferroni correction, \**P*<0.05; \*\**P*<0.01; \*\*\**P*<0.001). **u**, GTT measuring blood glucose levels following oral administration of danuglipron (oDan) and dextrose in Glp1r^S33W^ and WT mice (*n* = 6 per genotype, two-way ANOVA with Bonferroni correction, \*\**P*<0.01; \*\*\**P*<0.001). **v-y**, HFD intake 1, 2, and 4 hours after oral gavage of (**v**) danuglipron or vehicle and (**x**) orforglipron or vehicle (*n* = 8-9, two-way ANOVA with Bonferroni correction, \*\**P*<0.01; \*\*\**P*<0.001). Comparison of the 4th hour between intraperitoneal (IP) and oral routes of administration with (**w**) danuglipron or (**y**) orforglipron (two-way ANOVA with Bonferroni correction, \*\**P*<0.01; \*\*\**P*<0.001). **z**, 21-day weight changes of male mice on chronic HFD for >8 weeks treated with orforglipron or vehicle daily at ZT6 (*n* = 10 per injection, paired t-test, \*\*\**P*<0.001). Data are represented as means ± SEM. \**P*<0.05; \*\**P*<0.01; \*\*\**P*<0.001. See Supplementary Table 1 for statistical details for all figures and Extended Data figures.

To overcome the species specificity of small-molecule GLP1RAs and enable preclinical investigation, we engineered humanized GLP1R mouse models. Through integrated behavioral, neuroanatomical, and functional analyses, we uncovered a novel multi-synaptic hindbrain- amygdala-midbrain circuit that modulates rewarding food consumption through striatal dopaminergic signaling. This discovery advances our understanding of how GLP1R signaling influences not only feeding behavior but also reward processes, highlighting both the therapeutic potential and the need for caution as these treatments see broader use^7–14,45–47,48^.

### Generation and Validation of Humanized Glp1r^S33W^ Mice for Investigating Small-Molecule GLP1RAs

While peptide GLP1RAs, like liraglutide, reduce food consumption in C57BL/6J mice, many small-molecule GLP1RAs do not effectively activate rodent Glp1r (**Fig. 1b**) due to a single amino acid difference from tryptophan to serine at position 33 (**Fig. 1c**)^40,41,42^. Using CRISPR-Cas9-mediated genome editing, we inserted the S33W mutation into the mouse *Glp1r* locus, effectively humanizing it (**Fig. 1d,e**). Homozygous humanized Glp1r S33W mice (Glp1r^S33W^) and wild type (WT) littermates did not exhibit differences in respiratory exchange ratio (RER) (**Extended Data** Fig. 1a-d), energy expenditure (EE) (**Extended Data** Fig. 1e-h) or body weight (**Extended Data** Fig. 1i-j), demonstrating that Glp1r^S33W^ mice maintain normal metabolic functions and energy homeostasis comparable to WT littermates. To evaluate the functionality of the S33W mutation *in vivo*, we performed glucose tolerance tests (GTT) and found that liraglutide improved glucose tolerance in both Glp1r^S33W^ mice and WT mice (**Fig. 1f**), whereas danuglipron and orforglipron were effective only in Glp1r^S33W^ mice (**Fig. 1g,h**). Similar to a recently developed Eli Lilly Glp1r^S33W^ rat model^49^, these results demonstrate that Glp1r^S33W^ mice retain responsiveness to peptide-based GLP1RAs while gaining sensitivity to human-specific small molecule GLP1RAs, establishing a valuable model system for investigating this next-generation class of drugs.

### Small-Molecule GLP1RAs Mirror Liraglutide in Suppressing Feeding in Glp1r^S33W^ Mice

Beyond their effects on treating type II diabetes through enhancing insulin secretion and improving glucose control, GLP1RAs induce significant weight loss^1,5,34,50^. To systematically characterize their impact on distinct feeding modalities, we employed parallel behavioral paradigms examining both homeostatic and hedonic feeding patterns. Homeostatic feeding was quantified through standard diet (SD) consumption during the active phase (zeitgeber time (ZT) 12–16), while hedonic feeding was assessed via high-fat-diet (HFD) intake during the inactive phase (ZT 2–6) when baseline SD consumption is negligible^51,52,53^. After determining the minimal danuglipron dose that robustly suppressed intake (**Extended Data** Fig. 2a), we selected orforglipron and liraglutide doses to match its anorectic effect. At these doses, all three agonists markedly reduced active-phase SD consumption (**Fig. 1j,l,n**) and inactive-phase HFD feeding (**Fig. 1p,r,t**) in Glp1r^S33W^ mice. SD consumption following an overnight fast was likewise suppressed by each treatment (**Extended Data** Fig. 2b-d). As predicted by the species specific receptor activation profile, WT mice exhibited reduced SD (**Fig. 1i,k,m**) and HFD (**Fig. 1o,q,s**) intake exclusively following liraglutide administration. Notably, both liraglutide and orforglipron demonstrated sustained 24-hour inhibition of food intake (**Extended Data** Fig. 2e-g), consistent with their extended pharmacokinetic profiles relative to danuglipron (**Fig. 1a**). To validate the clinical relevance of these orally bioavailable small-molecule GLP1RAs, we confirmed that oral danuglipron significantly reduced blood glucose levels (**Fig. 1u**) and acute HFD intake comparable to intraperitoneal injection effects (**Fig. 1v, w**). Similarly, oral orforglipron inhibited acute HFD intake (**Fig. 1x,y**), and its chronic daily administration in overweight Glp1r^S33W^ mice significantly reduced body weight compared to vehicle controls (**Fig. 1z**), establishing its efficacy in weight management^34^.

### Divergent Behavioral Signatures Reveal Nausea-Like Effects of Liraglutide and Danuglipron, but Not Orforglipron

Because nausea often limits the clinical tolerance of GLP1RAs, we set out to distinguish nausea-like from satiety behaviors induced by peptide and small-molecule drugs. While conditioned taste avoidance (CTA) was evident following LiCl and liraglutide treatment as previously reported^23^, this assay failed to detect subtle potential malaise states in danuglipron or orforglipron treated mice (**Extended Data** Fig. 3a-b). As danuglipron and orforglipron also did not alter anxiety-like behavior in the open field or elevated plus maze (**Extended Data** Fig. 3c-l), we were able to leverage high-resolution home-cage tracking to capture nuanced behavioral differences (**Fig. 2a, Extended Data** Fig. 4). We first generated a “nausea” reference by treating mice with lithium chloride (LiCl) and a “satiety” reference by pre-feeding a HFD for one hour before recording. Using these benchmarks, we then compared home-cage behavior after liraglutide, danuglipron, orforglipron, and their respective vehicle controls to determine which profile each agonist most closely mimics. Mice were video recorded for two hours (ZT 12–14) in their home cages following each treatment (**Extended Data** Figs. 4a-c). Behavioral states were quantified using pose estimation (SLEAP) and a probabilistic classification model (Keypoint-MoSeq), which identified 91 behavioral syllables (**Fig. 2b, Extended Data** Fig. 5-6)^54,55^. These syllables were then grouped into five broader behavioral categories using human annotation and transition-network analysis: food-motivated, drinking, grooming, movement/exploration, and resting/grooming in shelter (**Fig. 2c**).

**Figure 2:**
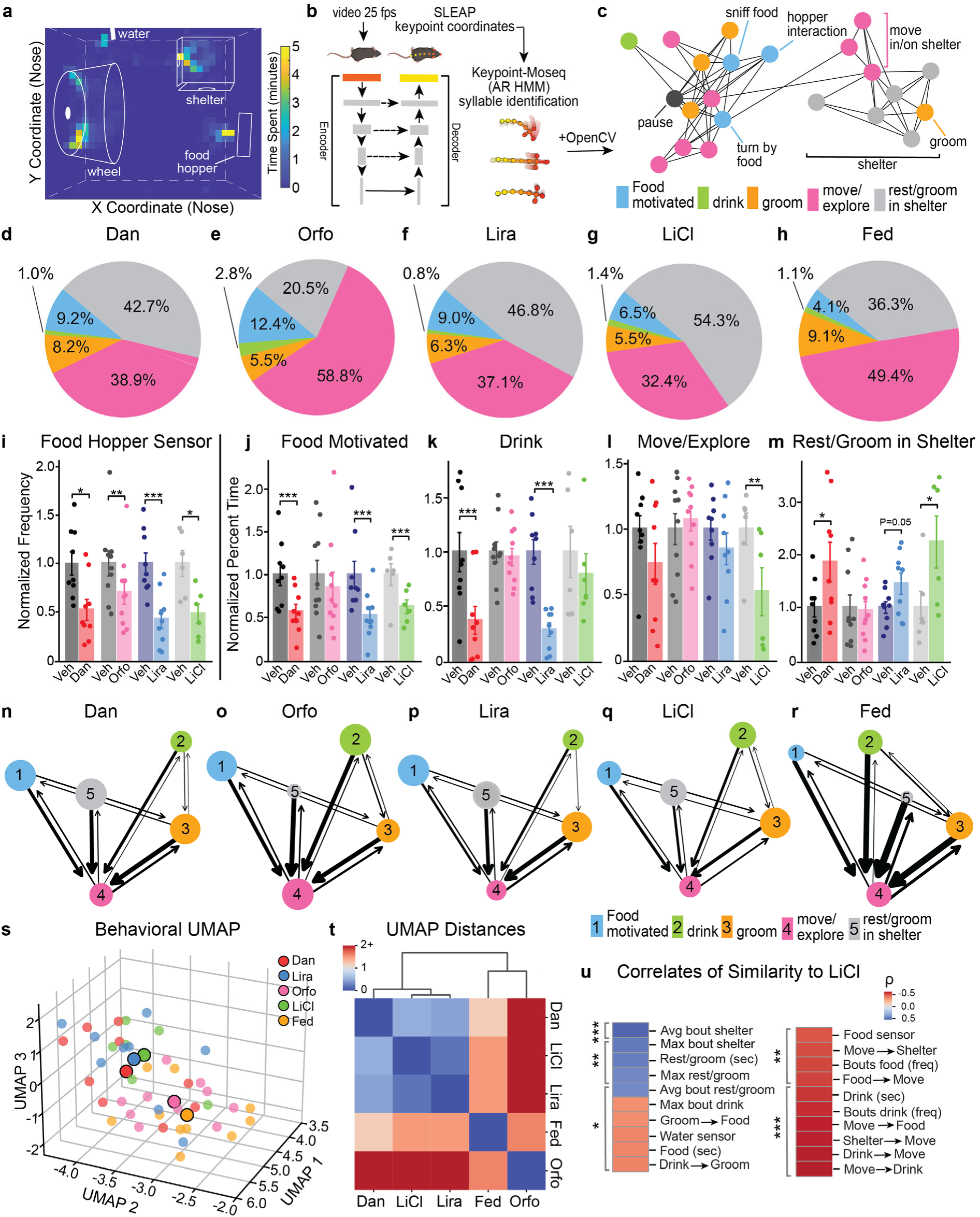
Machine-learning assisted behavior profiling reveals distinct phenotypes associated with GLP1RA, nausea/aversion, and satiety. **a**, Representative heatmap of Glp1r^S33W^ mouse nose location and home cage setup. Color indicates time spent in minutes over 2 hours. **b,** Simplified pipeline for machine learning analysis of behavior^54,55^. Analysis parameters: SLEAP tracking with 9 keypoints at 25 fps, Keypoint-MoSeq fit an autoregressive hidden Markov model (AR-HMM) to PCA-reduced keypoints. A full model was then trained, producing 91 behavioral syllables with a minimum frequency threshold of ≥0.01%. Trained raters manually named behaviors and grouped them by similar behavioral state, incorporating locational context via OpenCV. **c,** Network analysis of transitions between 22 behaviors, grouped into 5 behavioral categories. **d-h,** Proportion of time spent performing each of the 5 behavioral categories and averaged across all mice per condition. (*n* = 9 Dan, *n* = 10 Orfo, *n* = 9 Lira, *n* = 6 LiCl, *n* = 10 Fed). **i,** Total number (frequency) of food hopper head entries normalized to vehicle controls (*n* = 9 Veh/Dan, *n* = 10 Veh/Orfo, *n* = 9 Veh/Lira, *n* = 6 Veh/LiCl); statistics based on paired t-tests on raw data (\**P*<0.05, \*\**P*<0.01, \*\*\**P*<0.001). **j-m**, Percentage of time spent (**j**) food seeking, (**k**) drinking, (**l**) moving/exploring, and (**m**) resting/grooming in the shelter. Generalized linear mixed- effects model (GLMM) with a beta distribution fitted to raw proportion data, including a random intercept for mouse ID to account for paired design (\**P*<0.05, \*\**P*<0.01, \*\*\**P*<0.001). **n-r,** Network analysis of normalized behavioral category transitions of mice, averaged across each condition. Node color and number indicate behaviors, and arrows represent probability of transition. Arrows are weighted by transition probabilities and nodes are scaled by average bout length of behavior. Rare transitions (probability < 0.1, with the exception of *move/explore → drink*, which was retained due to behavioral relevance) were excluded from the plot. *Move/explore in shelter* was excluded from transition calculations to improve interpretability, as it often represents a transient state between shelter and other behaviors. (*n* = 9 Dan, *n* = 10 Orfo, *n* = 9 Lira, *n* = 6 LiCl, *n* = 10 Fed). **s**, UMAP of the full behavioral dataset (normalized transitions, average and max length of bouts of behaviors and total number of behavioral bouts, time spent performing each behavioral category, distance travelled, and food and water sensor data). PCA was used as a preprocessing step to decorrelate variables, with 16 PCs used, explaining 95% of variance. Opaque centroid circles represent the mean behavioral summaries per condition, small circles indicate the specific embedding locations of individual mice within each condition. **t**, Distance between average UMAP location of each condition with dendrogram. **u**, Behavioral features correlated with similarity to the behavioral signature of the LiCl condition. Only features with significant Spearman correlations (*FDR*-corrected, \**P*<0.05, \*\**P*<0.01, \*\*\**P*<0.001) are shown. Correlations reflect the relationship between each behavioral feature and Euclidean distance to the LiCl group centroid in UMAP space. Signs were flipped so that positive values indicate greater similarity to the LiCl behavioral profile, and negative values indicate greater dissimilarity.

Consistent with reduced intake, all GLP1RAs and LiCl suppressed food-hopper head entries. However, only LiCl, liraglutide, and danuglipron reduced time in food-motivated behaviors whereas liraglutide and danuglipron also decreased drinking (**Fig. 2d–k**). LiCl and liraglutide reduced distance traveled and increased grooming, while LiCl and danuglipron increased rest in shelter (**Fig. 2l,m; Extended Data** Fig. 7a–n). In contrast, orforglipron-treated mice maintained a more active, exploratory profile despite reduced food-hopper head entries. This distinct profile led to its significant separation from all other treatment groups in principal component analysis (PCA) space (**Extended Data** Fig. 7o,p), while liraglutide and danuglipron clustered together with a behavior repertoire characterized by LiCl-induced malaise. Remarkably, none of the treatments fully mimicked true satiety: fed mice spent significantly less time in food-motivated behaviors than any LiCl- or GLP1RA-treated group (**Extended Data** Fig. 7g). Their transition dynamics also stood apart, with fed mice making significantly more rest-to-exploration transitions than other cohorts (**Fig. 2n–r; Extended Data** Fig. 8a-x). PCA of these transition patterns and bout lengths placed the fed condition in its own cluster, clearly separated from all other treatments (**Extended Data** Fig. 8y–z).

To further characterize these distinct behavioral signatures, we performed a multivariate analysis incorporating time spent, transition probabilities, bout metrics, and sensor data, generating a Uniform Manifold Approximation and Projection (UMAP) embedding of the full dataset. The UMAP analysis revealed that LiCl, liraglutide, and danuglipron clustered closely, while orforglipron- treated and fed mice formed distinct, separate clusters (**Fig. 2s–t**). Greater UMAP distance from the LiCl cluster correlated with increased activity, exploration, and behavioral switching, with frequent transitioning out of the shelter being the strongest predictor of dissimilarity from LiCl (ρ=0.865, FDR *P*<0.0001) and longer shelter bouts reflecting similarity (ρ=–0.692, FDR *P*<0.0001) (**Fig. 2u**). These results demonstrate that liraglutide and danuglipron elicit behavioral states resembling LiCl-induced nausea, whereas orforglipron does not. Although all compounds reduce food intake, orforglipron-treated mice maintain a behavioral profile distinct from both nausea-like and satiety states, indicating a dissociation between appetite suppression and malaise-related side effects.

### GLP1RAs Exhibit Distinct Neural Activation Patterns in Feeding Circuits

Given the unique behavioral signatures induced by peptide and small-molecule GLP1RAs, and their distinct pharmacokinetic and likely brain-penetration profiles, we investigated the neural substrates mediating their anorectic effects. We began by examining nuclei known to express *Glp1r* and previously shown to be activated by peptide GLP1RAs (**Fig. 3a-d**), specifically the dorsomedial hypothalamus (DMH), nucleus tractus solitarius (NTS), area postrema (AP), and central amygdala (CeA)^21,24–26,56–59^. We quantified Fos expression as a proxy for GLP1RA-induced neuronal activation in WT and Glp1r^S33W^ mice following treatment with vehicle, danuglipron, orforglipron, or liraglutide (**Fig. 3a-d**). Danuglipron and orforglipron induced significant Fos expression in the NTS (**Fig. 3f**), AP (**Fig. 3g**), and CeA (**Fig. 3h**), but not in the DMH (**Fig. 3e**) of Glp1r^S33W^ mice compared to controls, mirroring previous studies using peptide-based GLP1RAs^21,56,60,61^. As expected, liraglutide induced comparable Fos expression in both groups, matching the small-molecule responses in Glp1r^S33W^ mice across all regions, confirming its effective binding to and activation of the Glp1r^S33W^ variant (**Fig. 3e-i**).

**Figure 3:**
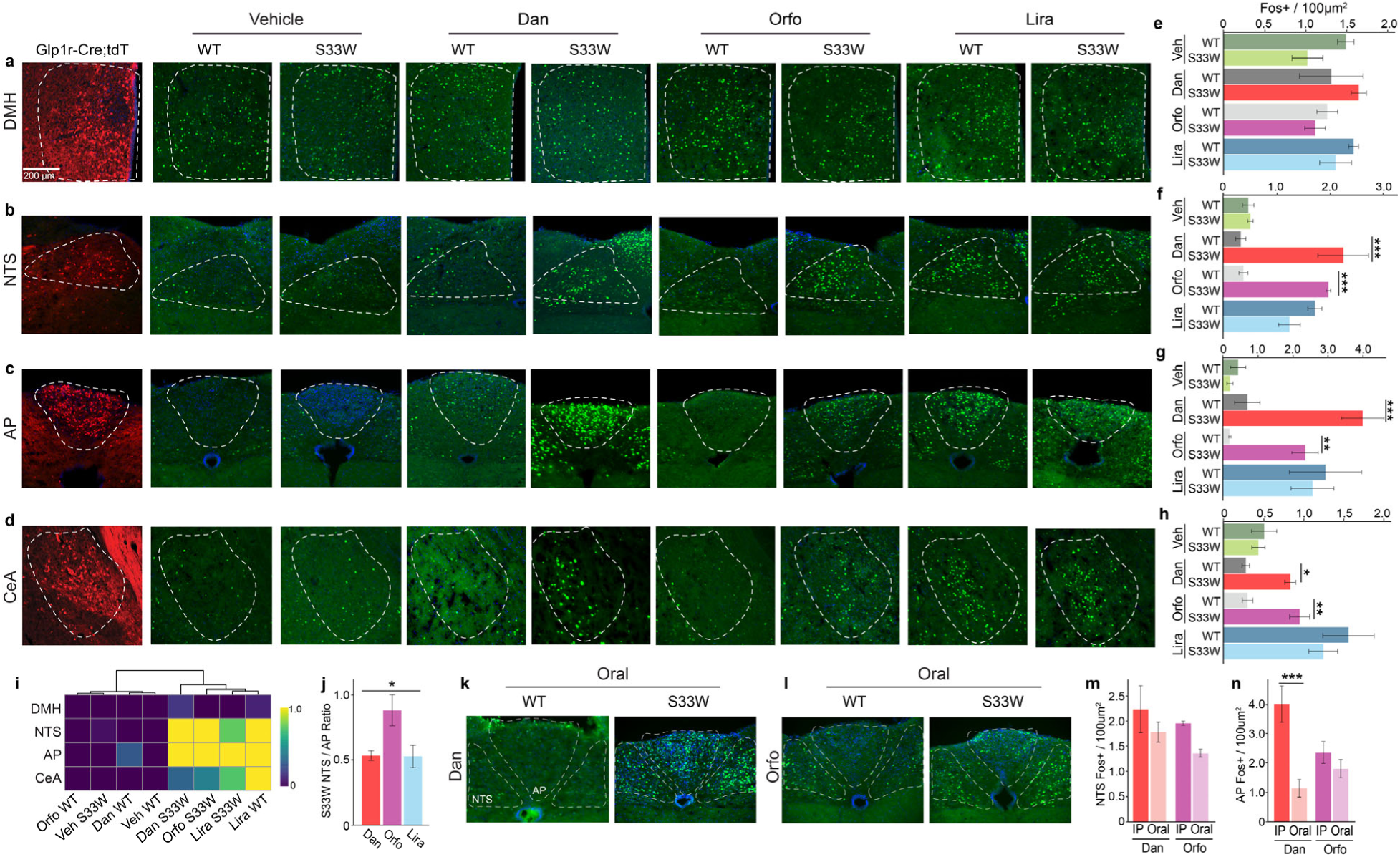
**GLP1RA activation across targeted GLP1R-expressing brain regions**. **a-d**, GLP1R protein expression validated by a Glp1r-Cre;tdTomato mouse line and neuronal Fos activation 2 hours after vehicle, danuglipron, or liraglutide or 6 hours after orforglipron injection in WT and Glp1r^S33W^ mice in the (**a**) DMH, (**b**) NTS, (**c**) AP, or (**d**) CeA. Scale bars = 200 µm. **e-h**, Quantification of Fos in the (**e**) DMH, (**f**) NTS, (**g**) AP and (**h**) CeA (*n* = 3-4 per genotype, two-way ANOVA with Bonferroni correction, \**P*<0.05; \*\**P*<0.01; \*\*\**P*<0.001). **i**, Heatmap of Fos activity in brain regions of interest across drug and vehicle conditions, normalized to WT vehicle. **j,k**, Neuronal Fos activation in the NTS and AP 2 hours after (**j**) danuglipron or (**k**) orforglipron was administered to WT and Glp1r^S33W^ mice via oral gavage. **l**, Ratio of NTS/AP Fos activation in Glp1r^S33W^ mice after danuglipron, orforglipron, and liraglutide (*n* = 3-4 per injection, Kruskal-Wallis test, \**P*<0.05). **m,n**, Quantification of Fos expression in the (m) NTS or (n) AP following danuglipron or orforglipron IP or oral delivery in Glp1r^S33W^ mice (*n* = 3-6 per delivery route, two- way ANOVA with Bonferroni correction, \*\*\**P*<0.001). Data are represented as means ± SEM. \**P*<0.05; \*\**P*<0.01; \*\*\**P*<0.001.

Given that AP activation drives nausea and malaise and NTS activation signals satiety^25,62^, we hypothesized that the balance of Fos induction between these regions might underlie the distinct behavioral profiles elicited by different GLP1RAs^5,34,63,64^. To test this, we calculated the NTS/AP Fos ratio for each treatment. Orforglipron produced a significantly higher NTS/AP ratio reflecting a satiety-dominant signal, whereas danuglipron and liraglutide both yielded lower ratios, consistent with a stronger nausea-like activation profile (**Fig. 3j**). Because these compounds are intended for oral use, we compared intraperitoneal versus oral delivery of danuglipron and orforglipron in Glp1r^S33W^ and WT mice to assess how administration route influences hindbrain Fos activation pattern (**Fig. 3k,l**). Glp1r^S33W^ NTS Fos activation was comparable between both routes and drugs (**Fig. 3m**), but oral delivery significantly reduced AP Fos activation after oral danuglipron (**Fig. 3n**). Preferential activation of the NTS over the AP may minimize nausea and malaise, side effects linked to AP engagement, while retaining NTS-mediated therapeutic benefits ^25^. By leveraging this profile, orally delivered small-molecule GLP-1R agonists offer a promising step forward in weight-loss treatment, combining strong efficacy with improved tolerability.

### Danuglipron Activates Hypothalamic and Hindbrain GLP1R Circuits

To distinguish direct from indirect effects of small-molecule GLP1RAs in specific brain regions, we developed a Cre-dependent adeno-associated virus (AAV) vector expressing full-length human GLP1R (AAV-hSyn-DIO-hGLP1R) (**Fig. 4a**). This approach enables selective expression of hGLP1R in Glp1r-positive cells of Glp1r-Cre mice (**Extended Data** Fig. 9a,b), allowing direct assessment of brain region specific GLP1R activation. In Glp1r-Cre mice conditionally expressing hGLP1R in their basomedial hypothalamus (BMH-hGLP1R) (**Extended Data** Fig. 9c,d), danuglipron significantly decreased active-phase SD intake without affecting inactive-phase HFD consumption (**Fig. 4c,h**) while control mice expressing mCherry in the same region showed no changes in consumption (**Fig. 4b,g**). When targeting the DMH, a BMH subnucleus recently implicated in encoding satiation^26,57,65^, we found similar selective effects on SD intake (**Fig. 4d,i and Extended Data** Fig. 9e,f). Targeting the hindbrain NTS/AP complex resulted in decreased consumption of both SD and HFD (**Fig. 4e,j and Extended Data** Fig. 9g,h), aligning with recent findings on hindbrain GLP1R circuits in aversion and satiety^25^.

**Figure 4:**
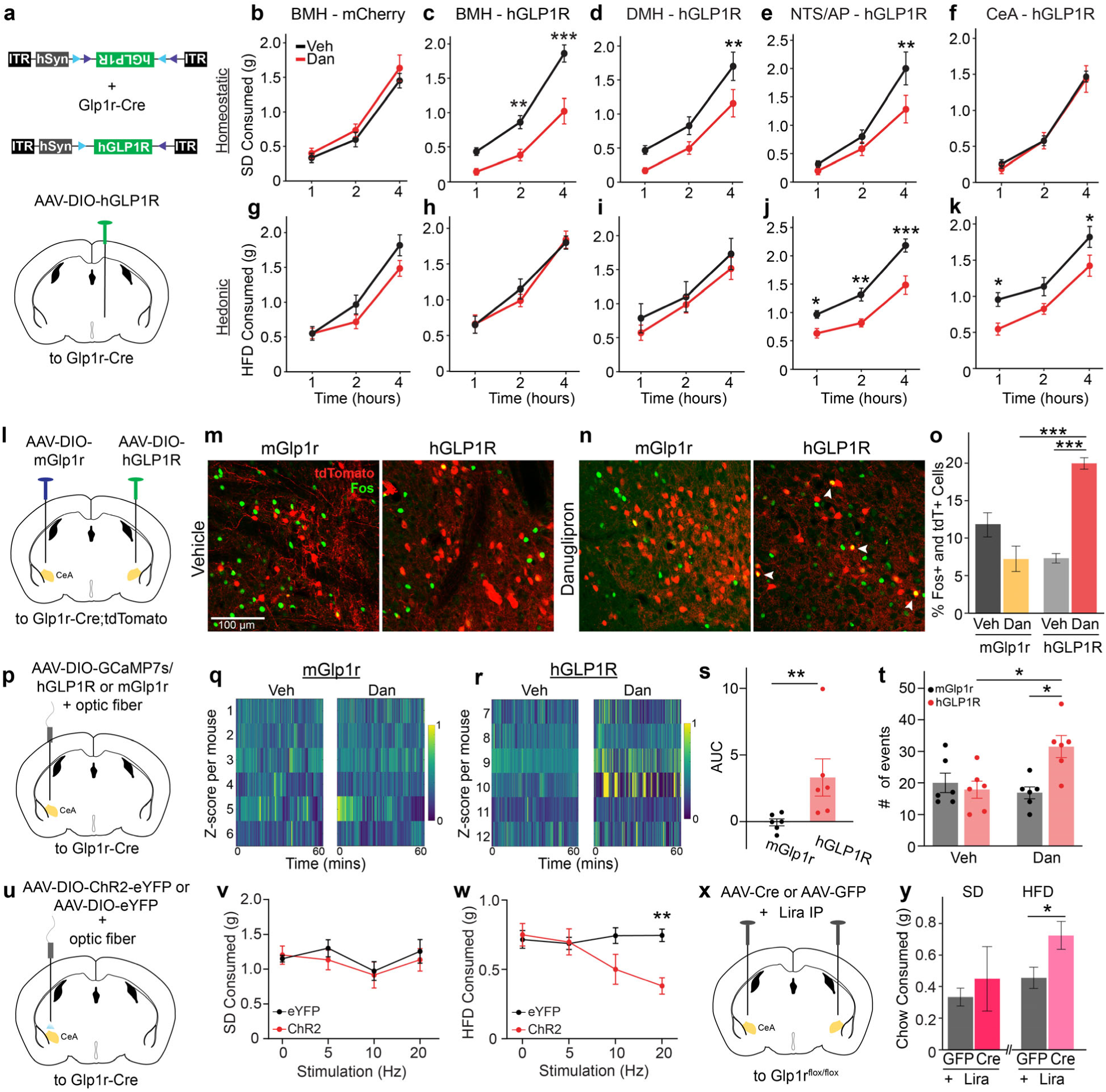
Central amygdala GLP1R activation inhibits hedonic food intake via direct activation by danuglipron. **a**, Representative schematic of AAV-DIO-hGLP1R viral construct and injection into the basomedial hypothalamus (BMH). **b-k**, 1, 2, and 4-hour SD or HFD consumption after vehicle (Veh) or danuglipron (Dan) in Glp1r-Cre mice expressing (**b,g**) AAV- DIO-mCherry (BMH-mCherry, *n* = 6) or (**c,h**) AAV-DIO-hGLP1R (BMH-hGLP1R, *n* = 10) in the BMH, (**d,i**) DMH (*n* = 7), (**e,j**) NTS/AP (*n* = 6), or (**f,k**) CeA (*n* = 9) (two-way ANOVA with Bonferroni correction, \**P*<0.05; \*\**P*<0.01; \*\*\**P*<0.001). **l,** Schematic of AAV-DIO-mGlp1r (left) and AAV-DIO- hGLP1R (right) injections in the CeA of Glp1r-Cre;tdTomato mice. **m,n**, Representative images of the CeA of (**m**) vehicle or (**n**) danuglipron injected mice with unilateral mGlp1r (left) and hGLP1R (right) infection (red = tdTomato; green = Fos; colocalization = white arrow). Scale bars = 100 µm. **o**, Quantification of the percent of Fos+ cells co-express tdTomato in the CeA of mGlp1r and hGLP1R infected mice (*n* = 3-4 per condition, two-way ANOVA with Tukey’s HSD correction, \*\**P*<0.01; \*\*\**P*<0.001). **p**, Schematic of AAV-DIO-GCaMP7s and AAV-DIO-hGLP1R or mGlp1r injection with a fiber optic implant in the CeA of Glp1r-Cre mice. **q,r**, Heatmaps of Z-scored neuronal calcium signals for each mouse during 1-hour of fiber photometry recording after vehicle or danuglipron injection in (**q**) mGlp1r or (**r**) hGLP1R expressing mice (n=6 mice per condition). **s,** Area under the curve (AUC) quantification of calcium recordings for mice expressing mGlp1r or hGLP1R after danuglipron injection (*n* = 6 mice per condition, unpaired t-test, \*\**P*<0.01). **t**, Number of significant calcium events averaged per condition during 1-hour recording session following vehicle or danuglipron in mGlp1r or hGLP1R expressing mice (*n* = 6 mice per injection, two-way ANOVA with Bonferroni correction, \**P*<0.05). **u,** Schematic of AAV-DIO-eYFP or AAV- DIO-ChR2-eYFP injection with a fiber-optic implant in the CeA of Glp1r-Cre mice. **v,w,** 1-hour SD (**v**) and 1-hour HFD (**w**) consumption of eYFP (control) or ChR2-expressing mice with different frequencies of light stimulation on different days (*n* = 5 per group, two-way ANOVA with Bonferroni correction, \**P*<0.05). **x**, Schematic of AAV-Cre to conditionally knockout Glp1r, or AAV-GFP control injection to the CeA of Glp1r^flox/flox^ mice. **y**, SD (left) and HFD (right) consumption 4 hours after injection of liraglutide in AAV-Cre or AAV-GFP expressing mice. (*n* = 6, Welch’s t-test,\**P*<0.05). Data are represented as means ± SEM. \**P*<0.05; \*\**P*<0.01; \*\*\**P*<0.001.

### Danuglipron Directly Activates CeA^Glp1r^ Neurons to Regulate Hedonic Feeding

While the CeA has been implicated in feeding behavior ^59,66–70^, its role in mediating responses to GLP1RAs remains poorly defined ^71^. Our finding that small-molecule GLP1RAs induce Fos expression in the CeA (**Fig. 3h**) raised several fundamental questions: What is the functional significance of CeA^Glp1r^ neurons in feeding behavior? Can danuglipron directly activate these deep brain neurons despite questions about blood-brain barrier penetration? And what is the neurochemical identity of these cells? To address the functional role, we expressed hGLP1R selectively in CeA neurons of Glp1r-Cre mice (CeA-hGLP1R mice). Remarkably, danuglipron selectively suppressed HFD intake without altering SD consumption in these mice (**Fig. 4f,k; Ext. Data Fig. 9i,j**). We confirmed this effect using an orthogonal viral strategy: Cre-dependent expression of the danuglipron-sensitive mGlp1r^S33W^ variant in CeA neurons reproduced HFD suppression, whereas wild-type mGlp1r expression served as an unresponsive control (**Extended Data** Fig. 10a-d). These data identify CeA^Glp1r^ neurons as a previously unrecognized yet vital substrate for GLP1RA-mediated suppression of hedonic feeding.

To characterize the neurochemical properties of these cells we took a bioinformatic and electrophysiological approach. We analyzed published CeA single-nucleus transcriptomic data, revealing that Glp1r-expressing neurons co-express GABAergic markers, lack glutamatergic markers and are in the cluster of CeA neurons that express the vitamin D receptor, *Vdr* (**Extended Data** Fig. 11a-c)^72^. To assess the neurophysiological impact of danuglipron on hGLP1R- expressing CeA neurons, we conducted whole-cell patch-clamp recordings following the co-injection of AAV-DIO-hGLP1R and AAV-DIO-eYFP into the amygdala of Glp1r-Cre mice (**Extended Data** Fig. 12a-f). We measured the resting membrane potential of hGLP1R- expressing neurons before and after infusion of 30 µM danuglipron (**Extended Data** Fig. 12a,b). Consistent with activation of the G_s_/cAMP pathway via hGLP1R^73,74^, danuglipron elicited significant membrane depolarization compared to mGlp1r-expressing control group, confirming functional expression (**Extended Data** Fig. 12c).

Danuglipron’s low molecular weight (555.6 Da) compared to peptide-based GLP1RAs and its ability to suppress hedonic feeding in mice expressing hGLP1R only in CeA^Glp1r^ neurons (**Fig. 4k**) indicates it crosses the blood–brain barrier to act on deep targets beyond the circumventricular organs (CVOs). Although the level of central nervous system penetration by different GLP1RAs remains debated^21,56,75,76,71,77^, defining their direct engagement of deep nuclei is essential. To demonstrate danuglipron’s direct activation of central Glp1r-neurons in freely moving mice, we employed two complementary approaches. First, we injected AAV-DIO-hGLP1R into one CeA and AAV-DIO-mGlp1r into the contralateral CeA of the same Glp1r-Cre mice (**Fig. 4l**). Following danuglipron treatment, Fos colocalization was significantly greater in hGLP1R-expressing neurons versus the mGlp1r side (**Fig. 4m–o**), confirming receptor-specific activation in deep brain tissue. Second, we co-expressed GCaMP7s with either hGLP1R or mGlp1r in the CeA of Glp1r- Cre mice and recorded calcium transients via fiber photometry (**Fig. 4p; Extended Data** Fig. 13a–d). Building on prior work and our own finding that peptide GLP1RAs induce CeA Fos expression in wild-type mice (**Figure 3d,h**)^21,56,60,61^, liraglutide evoked robust calcium responses in CeA-hGLP1R mice, as expected (**Extended Data** Fig. 13e–g). Importantly, danuglipron similarly evoked significant calcium transients only in hGLP1R-expressing mice, with no evoked responses in mGlp1r negative controls (**Fig. 4q–t**). Together, these findings demonstrate that danuglipron crosses the blood–brain barrier to directly activate deep-brain CeA neurons via the human GLP1R.

To define the physiological role of CeA^Glp1r^ neurons in hedonic feeding, we expressed ChR2 in the CeA of Glp1r-Cre mice and delivered light pulses during one-hour feeding sessions. Optogenetic activation significantly reduced HFD consumption without affecting SD intake (**Fig. 4u–w**), recapitulating the anorectic effects of danuglipron in CeA-hGLP1R mice (**Fig. 4k; Ext. Data Fig. 10b,c**). To test whether CeA *Glp1r* is required for GLP1RA-mediated feeding suppression, we bilaterally injected AAV-Cre or AAV-GFP into the CeA of Glp1r^flox/flox^ mice (**Fig. 4x**). Deletion of CeA *Glp1r* had no effect on SD intake but significantly reduced liraglutide’s ability to inhibit HFD consumption (**Fig. 4y**). Together, these results establish CeA^Glp1r^ neurons as a critical node in the circuit through which GLP1RAs suppress reward-driven feeding.

### Small-Molecule GLP1RAs Engage the CeA to Modulate Mesolimbic Dopamine Release in Response to HFD

To further clarify this circuit’s physiological relevance, we investigated whether CeA^Glp1r^ neurons receive endogenous GLP-1 input. We selectively expressed fluorescently-tagged synaptophysin in NTS^Gcg^ neurons, the most likely source of GLP-1 in the CNS ^78^. This allowed us to detect NTS^Gcg^ synaptic terminals in the CeA (**Fig. 5a**), indicating that CeA^Glp1r^ neurons are capable of receiving both GLP-1 and other presynaptic input from NTS^Gcg^ neurons. To functionally confirm this connection, we Cre-dependently expressed ChrimsonR or mCherry in the NTS of Gcg-Cre mice and placed optic fibers bilaterally in their CeA (**Fig. 5b**). Stimulating CeA-projecting NTS^Gcg^ axon terminals significantly inhibited HFD but not SD intake (**Fig. 5c,d**). These findings underscore the CeA as a key integrative hub where GLP-1 signaling, both pharmacologically induced and naturally occurring, converges to curb hedonic feeding^79^.

**Figure 5:**
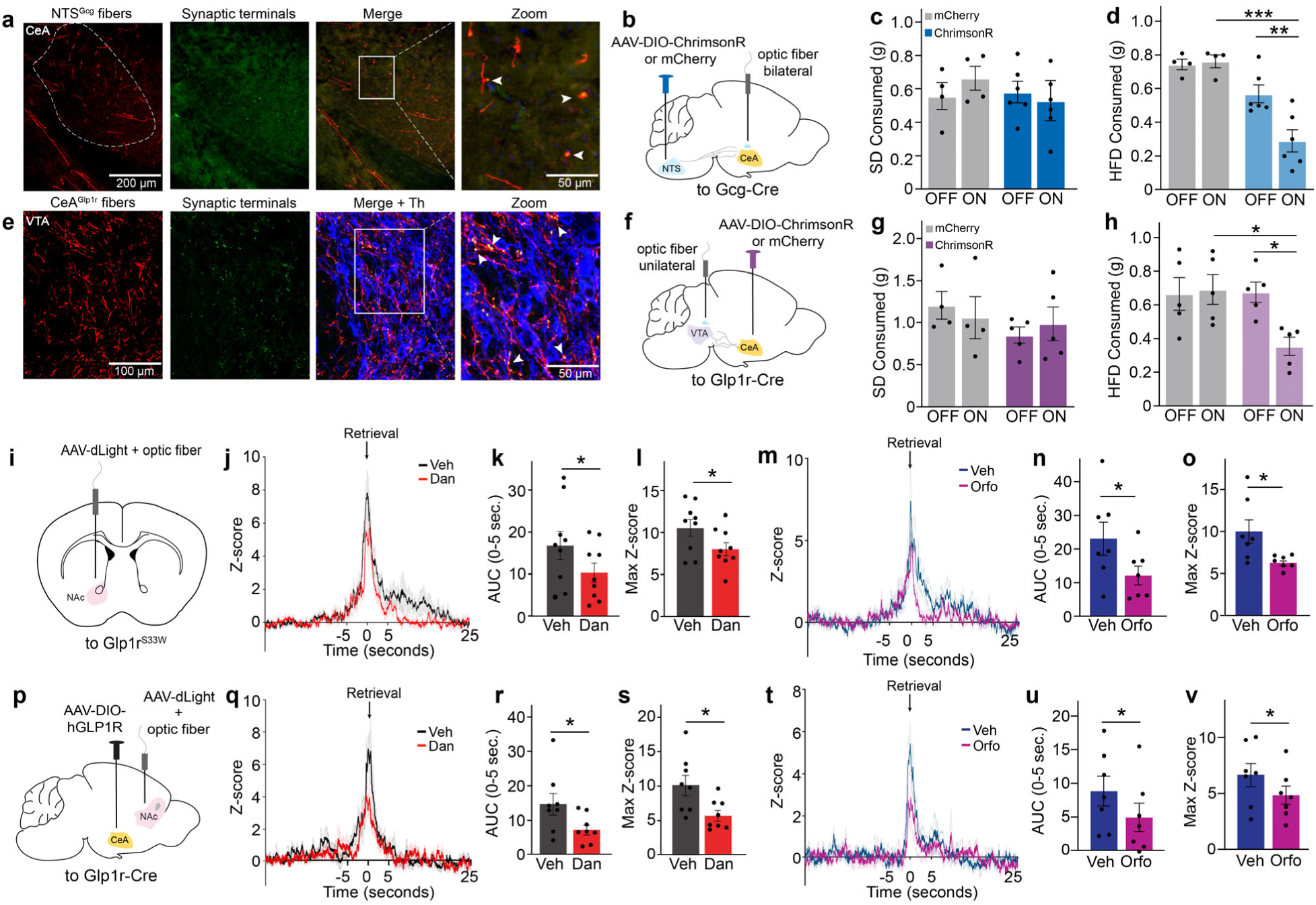
CeA^Glp1r^ neurons project to the VTA to modulate dopamine output to the NAc in response to HFD. a, Representative image of the CeA following injection of AAV-DIO-mGFP- 2A-Synaptophysin-mRuby into the NTS of Gcg-Cre mice. Fibers are shown in red (pseudo- colored mGFP) and synaptic terminals in green (pseudo-colored mRuby). Magnified view (right) highlights synaptic terminals (white arrows). Scale bars: a, 200 µm (main) and 50 µm (magnified). **b**, Schematic of AAV-DIO-ChrimsonR or AAV-DIO-mCherry injection into the NTS of Gcg-Cre mice with bilateral fiber-optic implants in the CeA. **c,d**, 1-hour consumption of (**c**) standard diet (SD) or (**d**) high-fat diet (HFD) in mCherry (control) versus ChrimsonR-expressing mice under light ON versus OFF conditions (*n* = 4–6 per group, two-way ANOVA with Tukey’s HSD, \*\**P*<0.01; \*\*\**P*<0.001). **e,** Representative image of the VTA following injection of AAV-DIO- mGFP-2A-Synaptophysin-mRuby into the CeA of Glp1r-Cre mice. Fibers are shown in red (pseudo-colored mGFP) and synaptic terminals in green (pseudo-colored mRuby). Magnified view (right) highlights synaptic terminals (white arrows). Scale bars: a, 100 µm (main) and 50 µm (magnified). **f**, Schematic of AAV-DIO-ChrimsonR or AAV-DIO-mCherry injection into the CeA of Glp1r-Cre mice with a unilateral fiber optic implant in the VTA. **g,h**, 1-hour consumption of (**g**) SD or (**h**) HFD in mCherry versus ChrimsonR-expressing mice under light ON versus OFF conditions (*n* = 4-5, two-way ANOVA with Tukey’s HSD, *P<0.05). **i**, Schematic of genetically-encoded dopamine sensor, AAV-dLight1.3b, injection and fiber optic implant in the NAc of Glp1r^S33W^ mice. **j,m**, Averaged Z-score traces showing dopamine release in the NAc in response to HFD following administration of (**j**) vehicle or danuglipron and (**m**) vehicle or orforglipron in Glp1r^S33W^ mice. Traces are aligned to food retrieval time (t = 0) and averaged across five food trials per mouse. **k-o**, Quantified (**k,n**) area under the curve (AUC) for Z-scores and (**l,o**) maximum fluorescence Z-scores within the food retrieval window (*n* = 9 for danuglipron, *n* = 7 for orforglipron, paired t- test, \**P*<0.05). **p**, Schematic of AAV-dLight1.3b injection and fiber optic implant into the NAc and AAV-DIO-hGLP1R injection into the CeA of Glp1r-Cre mice. **q,t**, Averaged Z-score traces showing dopamine release in the NAc in response to HFD following administration of (**q**) vehicle or danuglipron and (**t**) vehicle or orforglipron in CeA-hGLP1R mice. Traces are aligned to food retrieval time (t = 0) and averaged across five food trials per mouse. **r-v**, Quantification of (**r,u**) AUC for Z-scores and (**s,v**) maximum fluorescence Z-scores within the food retrieval window (*n* = 8 for danuglipron, *n* = 7 for orforglipron, paired t-test, \**P*<0.05). Data are represented as means ± SEM. \**P*<0.05; \*\**P*<0.01; \*\*\**P*<0.001.

Having established that CeA^Glp1r^ activity selectively suppresses hedonic feeding, we next asked whether this amygdalar node feeds directly into the mesolimbic reward pathway, paralleling other GLP1R-based circuits that impinge on midbrain dopamine neurons^80,81^. Anatomical tracing from the CeA of Glp1r-Cre mice, using AAV-DIO-mGFP-2A-Synaptophysin-mRuby or AAV-DIO-ChR2- eYFP, revealed pronounced projections from CeA^Glp1r^ neurons to the ventral tegmental area (VTA) (**Fig. 5e and Extended Data** Fig. 14a-e). In complementary experiments, fluorescent marker expression from retroAAV-oNLS-oScarlet delivered into the VTA co-localized with Cre- dependent ChR2-eYFP expression in the CeA of Glp1r-Cre mice (**Extended Data** Fig. 15a-c) confirming direct CeA^Glp1r^ to VTA connectivity. Furthermore, monosynaptic rabies tracing of VTA dopamine cells labeled CeA^Glp1r^ neurons (**Extended Data** Fig. 15d-f) demonstrated that CeA^Glp1r^ neurons make synaptic connections with the VTA dopamine neurons. Finally, to test functional relevance, we expressed Cre-dependent ChrimsonR or mCherry in CeA^Glp1r^ neurons and implanted an optic fiber in the VTA. Optogenetic stimulation of CeA axon terminals in the VTA selectively reduced HFD intake without affecting SD consumption (**Fig. 5f–h**), demonstrating that CeA^Glp1r^ to VTA signaling is sufficient to inhibit hedonic feeding.

These findings support a model in which NTS^Gcg^-driven activation of GABAergic CeA^Glp1r^ neurons suppresses VTA dopamine output and thereby blunt accumbal dopamine release during hedonic feeding. To test this, we virally expressed the fluorescent dopamine sensor dLight1.3b in the nucleus accumbens (NAc) of Glp1r^S33W^ mice and recorded dopamine transients in response to HFD consumption following administration of liraglutide, danuglipron, or orforglipron (**Fig. 5i**). All three GLP1RAs significantly attenuated both the peak and the consumption-associated dopamine signals (**Fig. 5j-o and Extended Data** Fig. 16a-d). In contrast, danuglipron had no effect on HFD- evoked dopamine release in wild-type mice (**Extended Data** Fig. 16e-h), confirming that these small molecules require the humanized receptor to dampen reward-related dopamine signaling. To establish a causal link between CeA^Glp1r^ neuron activity and mesolimbic dopamine output, we selectively expressed hGLP1R in the CeA of Glp1r-Cre mice and monitored NAc dopamine dynamics during HFD consumption following danuglipron or orforglipron administration (**Fig. 5p; Extended Data** Fig. 16i-k). Strikingly, both small-molecule treatments markedly blunted the peak and consumption-associated dopamine transients (**Fig. 5q-v**), demonstrating that activation of CeA^Glp1r^ neurons suppresses reward-driven dopamine signaling. By showing that this CeA→VTA→NAc pathway operates alongside previously characterized hindbrain→midbrain GLP1R circuits^81–83^, our results reveal a distributed network of GLP1R-expressing neurons orchestrating the suppression of food consumption^25,26,84,85,25,84,86–88^.

## Discussion

Our data uncover a previously uncharacterized amygdalar pathway through which next- generation GLP1R agonists modulate reward-driven feeding. Specifically, CeA^Glp1r^ neurons integrate exogenous and endogenous GLP-1 input and modulate VTA activity, thereby linking metabolic signals to dopamine-dependent reward circuits^10,81,86–88^. This circuit complements established hindbrain→midbrain GLP1R pathways and helps explain how GLP1R agonists influence not only energy balance but also motivational and reward-related behaviors^12,13,14,48,89,90^. More broadly, our findings suggest potential applications of GLP1R agonists in disorders of dysregulated reward signaling, including substance-use disorders. As small-molecule compounds like orforglipron—hailed as "a product for the masses"—move swiftly toward widespread clinical use^33^, it will be critical to define their long-term effects on brain reward circuits and motivated behaviors.

## Methods

### Mouse lines

All experiments were carried out in compliance with the Association for Assessment of Laboratory Animal Care policies and approved by the University of Virginia Animal Care and Use Committee. Mice were housed on a 12:12-hour light/dark (LD) cycle with food (PicoLab Rodent Diet 5053) and water *ad libitum* unless otherwise indicated. For experiments, we used 8-week or older male and female C57BL/6J mice, Glp1r-IRES-Cre mice (Glp1rtm1.1(cre)Lbrl/RcngJ, strain #029283, RRID:IMSR_JAX:029283), Glp1r-IRES-Cre mice crossed to Ai14 tdTomato reporter line (B6.Cg-Gt(ROSA)26Sortm14(CAG-tdTomato)Hze/J, strain #007914, RRID:IMSR_JAX:007914), Glp1r^flox/flox^ mice (B6(SJL)-Glp1rtm1.1Stof/J, strain #035238, RRID:IMSR_JAX:035238), Dat-Cre mice (B6.SJL-Slc6a3tm1.1(cre)Bkmn/J, strain #006660, RRID:IMSR_JAX:006660) Gcg-Cre mice (C57BL/6J-Tg(Gcg-cre)-1Mmsc/Mmmh, stock #051056-MU, RRID:MMRRC_051056-MU), and Glp1r^S33W^ mice (described below). Gcg-Cre mice were rederived by *in vitro* fertilization from frozen sperm (MMRRC, stock no. 051056-MU).

### Generation of Glp1r^S33W^ mouse

The Glp1r^S33W^ mouse line was created with CRISPR-Cas9 homologous repair at the University of Virginia Genetically Engineered Murine Model Core. Briefly, Cas-9 (Alt-R™ S.p. Cas9 Nuclease V3, 100 µg, catalog no. 1081058), Alt-R™ HDR Donor Oligo repair template (below), tracrRNA (Alt-R® CRISPR-Cas9 tracrRNA, 5 nmol, catalog no. 1072532), and CRISPR-Cas9 crRNA XT (ATTTCTGCACCGTCTCTGAG) were microinjected into a fertilized B6SJL zygote and were implanted into a pseudopregnant female. Founder pups were genotyped as described below and backcrossed to C57BL/6J mice for at least 4 generations before experimentation. This strain will be available at The Jackson Laboratory Repository with the JAX Stock No. 040551 Glp1r^S33W^ mouse line.

*Repair Template* aagagggtgggagtccagtgggaccagaggggctgctggagccacggggcttctgcttttatttctgctttcccttgtagGGTACCA CGGTG**TCGCTC<u>TGG</u>GAAACCGTCCAAAAGTGG**AGAGAATACCGGCGGCAGTGCCAGCGTTTCCTCACGGAAGCGCCACTCCTGGCCACAGgtgcgtccagatgaggcctcacg

### Validating Glp1r^S33W^ mice

Tail snips were obtained from pups at 3 weeks of age. DNA was extracted with an extraction buffer (Sigma, catalog no. E7526) and tissue prep solution (Sigma, catalog no. T3073), heated for 10 and 3 minutes at 55°C and 100°C, respectively, then neutralized with a neutralization solution (Sigma, catalog no. N3910). PrimeSTAR High Fidelity PCR (Takara, catalog no. R050A) was performed with 1 µL of cDNA and 10 µM of 5′-3′ F (GATCCCCAAAGTGGCAGTCA) and 5′- 3′ R (AGCTATGGACTGGGGATCGT) primers. After amplification, the PCR product was run on a 1.2% agarose gel and bands were cut out at 330 bp. DNA was gel extracted and purified (Qiagen, catalog no. 28704), mixed with 5 µM right primer, H_2_O, and subsequently sent to be analyzed via Sanger sequencing (Azenta). Chromatogram results were analyzed to assign wild type, heterozygous, or homozygous genotypes for each mouse.

### Generation of GLP1R viruses

The full length human *Glp1r* gene was obtained by PCR, amplifying the human fragment from GLP1R-tango (plasmid from Addgene, #66295, RRID:Addgene_66295), including the leader sequence present in the GLP1R-tango. The primers used were: 5′-3′ F (AAAGCT- AGCGCCACCATGAAGACGATCATCGCCCTGAGC) and 5′-3′ R (TTTGGCGCGCCCTAA-GAGCAGGACGCCTGACAAGT), ligating the product into pAAV-hSyn-DIO-EGFP (plasmid from Addgene, #50457, RRID:Addgene_50457) in place of the EGFP in NheI and AscI sites to produce the human GLP1R virus construct (AAV-hSyn-DIO-hGLP1R). The full length mouse *Glp1r* wild type gene was synthesized by Twist Biosciences (USA), generating a NheI and AscI fragment. This construct included the same leader sequence present in the human construct, as well as an HA-tag encoded at the C-terminus of the full length mouse protein coding region. The fragment was inserted into pAAV-hSyn-DIO-EGFP (plasmid from Addgene, #50457, RRID:Addgene_50457) in place of EGFP to produce the plasmid construct (AAV-hSyn-DIO- mGLP1R-HA). The full length mouse *Glp1r* gene bearing a Ser to Trp mutation at position 33 (S33W) was made by inserting a synthetic NheI and StuI fragment prepared by Twist Biosciences, containing the single mutation within this fragment. This was cloned into the sites present in the wild type construct to produce the S33W mouse mutant, followed by an HA-tag encoded at the C-terminus (AAV-hSyn-DIO-mGLP1R^S33W^-HA). Viral plasmid constructs were confirmed by Sanger sequencing. Virus plasmid constructs were prepared and sent to the University of North Carolina Viral Core (Chapel Hill, NC) for preparation of the AAV (serotype 8). In experiments using AAV-DIO-hGLP1R/mGlp1r/mGLP1R^S33W^-HA to drive receptor expression, viral titers were carefully calibrated to approximate endogenous receptor levels (Extended Data Fig. 9); nonetheless, this overexpression approach may alter receptor distribution or signaling and constitutes a limitation of the model.

### Stereotactic surgery

Mice were anesthetized with isoflurane (5% induction and 2 to 2.5% maintenance; Isothesia) and placed in a stereotaxic apparatus (AWD). A heating pad was used for the duration of the surgery to maintain body temperature, and ocular lubricant was applied to the eyes to prevent desiccation. A total of 200-400 nL was microinjected per side of [rAAV8/AAV2-hSyn-DIO-hGLP1R, plasmid from Addgene, virus packed at UNC GTC Vector Core, Lot #AV9862 (100 µL at titer ≥ 1.5 ×10^13^ vg/mL); AAV8-hSyn-DIO-mGLP1R^S33W^-HA, synthesized by Twist Biosciences, virus packed at UNC GTC Vector Core, Lot #AV10104 (100 µL at titer ≥ 8.2×10^12^ vg/mL); AAV8-hSyn-DIO- mGLP1R-HA, synthesized by Twist Biosciences, virus packed at UNC GTC Vector Core, Lot #AV10103 (100 µL at titer ≥ 4.5×10^12^ vg/mL); pAAV9-syn-dLight1.3b, plasmid from Addgene, #135762, RRID:Addgene_135762, virus packed at UNC GTC Vector Core (100 µL at titer ≥ 1.5×10^13^ vg/mL); pAAV1-EF1a-DIO- hChR2(H134R)-EYFP-WPRE-HGHpA, plasmid from Addgene, #20298, RRID:Addgene_20298, virus packed at UNC GTC Vector Core (100 µL at titer ≥ 7×10¹² vg/mL); pGP-AAV1-syn-DIO-jGCaMP7s-WPRE, plasmid from Addgene, #104491, RRID: Addgene_104491, virus packed at UNC GTC Vector Core (100 µL at titer ≥ 1×10¹³ vg/mL); pAAV1-Ef1a-DIO-EYFP, plasmid from Addgene, #27056, RRID:Addgene_27056, virus packed at UNC GTC Vector Core (100 µL at titer ≥ 1×10¹³ vg/mL); AAV8-hSyn-DIO-mCherry, plasmid from Addgene #50459, RRID:Addgene_50459, and virus packed at UNC Vector Core (100 µL at titer ≥ 7×10¹² vg/mL); pAAV-hSyn-FLEx-mGFP-2A-Synaptophysin-mRuby, plasmid from Addgene, #71760, RRID:Addgene_71760, and virus packed at UNC Vector Core; pENN.AAV.hSyn.HI.eGFP-Cre.WPRE.SV40, plasmid from Addgene, #105540-AAV8, RRID:Addgene_105540-AAV8, and virus packed at UNC Vector Core (100 µL at titer ≥ 1×10¹³ vg/mL); pAAV-hSyn-DIO-ChrimsonR-mRuby2-ST, plasmid from Addgene, #105448, RRID:Addgene_105448-AAV9, and virus packed at UNC Vector Core (100 µL at titer ≥ 1×10¹³ vg/mL); pAAV-hSyn-EGFP, plasmid from Addgene, #50465-AAV8, RRID:Addgene_50465- AAV8, and virus packed at UNC Vector Core (100 µL at titer ≥ 7×10¹² vg/mL)], retroAAV2-EF1a- oNLS-oScarlet ^91^ (100 µL at titer ≥ 3×10¹³ vg/ml), virus was delivered using a glass pipette at a flow rate of 50 nl/min driven by a microsyringe pump controller (World Precision Instruments, model Micro 4). The syringe needle was left in place for 10 min and was completely withdrawn 17 min after viral delivery. For *in vivo* calcium and dopamine imaging and optogenetics, a unilateral fiber optic cannula (RWD, Ceramic Ferrule, Ø400-μm, 0.5 numerical aperture) was implanted 0.2- mm dorsal to the viral injection coordinates following viral delivery and stabilized on the skull with dental cement (C&B METABOND, Parkell). Three weeks minimum were allowed for recovery and transgene expression after surgery. Stereotaxic coordinates relative to Bregma (George Paxinos and Keith B. J. Franklin): basomedial hypothalamus (encompassing the dorsomedial hypothalamus, arcuate nucleus, median eminence, and ventromedial hypothalamus), mediolateral (ML): ±0.3 mm, anterior posterior (AP): −1.4 mm, dorsoventral (DV): −5.9 mm; DMH, ML: ±0.3 mm, AP: −1.8 mm, DV: −5.4 mm; CeA, ML: ±2.7 mm, AP: −1.3 mm, DV: −4.6 mm; VTA, ML: ±0.5 mm, AP: −3.6 mm, DV: −4.5 mm; NAc, ML: ±1.25 mm, AP: +1.0 mm, DV: −4.7 mm from Bregma; and NTS/AP, ML: ± 0.15 mm, AP: −0.3 mm, DV: −0.1, −0.4 mm from the Zero point of the calamus scriptorius.

For rabies tracing, A total of 500 nL, containing a 1:1 mixture of AAV5-FLEx^loxP^-TC (UNC Vector Core, titer ≥ 2.4×10^12^ gc/mL) and AAV8-FLEx^loxP^-RABV-G (UNC Vector Core, titer ≥ 1.0×10^12^ gc/mL), was injected into the VTA of DAT-Cre mice. Fourteen days later, 500 nL of G-deleted, GFP-expressing, EnvA-pseudotyped rabies virus (RABVΔG-H2B-GFP-EnvA, (generated at UC Irvine, Beier lab, titer ≥ 5×10^8^ colony forming unit (cfu)/mL) was injected into the same site. Five days following the RABV injection, brains were collected for further processing. All surgical procedures were performed under sterile conditions and in accordance with University of Virginia Institutional Animal Care and Use Committee guidelines. Histological analysis was performed to validate the success of intracranial surgeries. Mice with unsuccessful viral/implant targeting were excluded from the analysis.

### GLP1R agonists

Liraglutide powder (Selleck, catalog no. S8256) was dissolved in 0.9% NaCl sterile saline, lightly sonicated, and further diluted in 0.9% NaCl sterile saline to 0.03mg/mL. Danuglipron powder (Selleck, catalog no. S9851) was dissolved to 30 mg/mL in 100% ethanol with gentle sonication, then diluted to 3mg/mL (food intake) or 0.3mg/mL (GTT) in vehicle [1N NaOH, 2% Tween 80, 5% polyethylene glycol (PEG) 400, 5% dextrose]^37^. Orforglipron powder (MedChemExpress, catalog no. HY-112185) was dissolved to 10mg/mL in Dimethyl sulfoxide (DMSO) and further diluted in 0.9% NaCl sterile saline to 0.1mg/mL. Dosage for danuglipron and orforglipron was decided based on previous literature.

### Histological analysis and imaging

For fixed tissue collection, mice were deeply anesthetized (ketamine:xylazine, 280:80 mg/kg, intraperitoneally) and perfused intracardially with ice-cold 0.01M phosphate buffer solution (PBS), followed by fixative solution [4% paraformaldehyde (PFA) in PBS at a pH of 7.4]. For testing brain region Fos activation (Fig. 3), vehicle (control), danuglipron (30 mg/kg), orforglipron (1 mg/kg), or liraglutide (0.3 mg/kg) was delivered via intraperitoneal injection or oral gavage 2 hours (or 6 hours for orforglipron) before perfusion and brain harvesting. After perfusion, brains were harvested and postfixed overnight at 4°C in PFA. Fixed brains were then transferred into 30% sucrose in PBS for 24 hours and then frozen on dry ice. Frozen brains were sectioned immediately or stored in −80°C for future processing. Coronal sections (30 μm) were collected with a cryostat (Microm HM 505 E). Sections were permeabilized with 0.3% Triton X-100 in PBS (PBS-T) and blocked with 3% normal donkey serum (Jackson ImmunoResearch, RRID:AB_2337258) in PBS- T (PBS-T DS) for 30 min at room temperature. Sections were then incubated overnight at room temperature in primary antibodies diluted in PBS-T DS. For visualization, sections were washed with PBS-T and incubated with appropriate secondary antibodies diluted in the PBS-T DS for 2 hours at room temperature. Sections were washed three times with PBS and mounted using DAPI Fluoromount-G (Southern Biotech, catalog no. 0100-20). Images were captured on a Zeiss Axioplan 2 Imaging microscope equipped with an AxioCam MRm camera using AxioVision 4.6 software (Zeiss) or confocal microscope imaging was performed on a Zeiss LSM 800 microscope (Carl Zeiss). The following primary antibodies were used for fluorescent labeling: anti-c-Fos (rabbit, 1:1000; Synaptic Systems, #226003, RRID:AB_2231974), anti-DsRed (rabbit, 1:1000; Takara Bio, catalog no. 632496, RRID:AB_10013483), anti-TdTomato (goat, 1:1000; Arigobio, catalog no. ARG55724), anti-hGLP1R (rabbit, 1:200; Invitrogen, catalog no. PA5-97789, RRID: AB_2812404), anti-HA (rabbit, 1:1000, Cell Signaling, catalog no. 3724), anti-Th (rabbit, 1:500; Chemicon, catalog no. AB152), and anti-GFP (goat, 1:500; Rockland, catalog no. 600-101-215). The secondary antibodies (Jackson ImmunoResearch) used were Cy2-conjugated donkey anti-rabbit (1:250; catalog no. 711-225-152, RRID:AB_2340612), Cy3-conjugated donkey anti-rabbit (1:250; catalog no. 711-165-152, RRID:AB_2307443), Cy5-conjugated donkey anti-rabbit (1:250; catalog no. 711-175-152, RRID:AB_2340607), Cy3-conjugated donkey anti-goat (1:250; catalog no. 705-165-147, RRID:AB_2307351), and Alexa-Fluor® 488 donkey anti-goat (1:250; catalog no. 705-545-003, RRID:AB_2340428).

### Antigen retrieval for hGLP1R staining

Antigen retrieval was performed before immunohistochemistry staining of human GLP1R, by incubating the sections in the following solutions sequentially in room temperature: 1% NaOH + 0.3% H_2_O_2_ in PBS for 20 min, 0.3% glycine in PBS for 10 min, and 0.03% sodium dodecyl sulfate (SDS) in PBS for 10 min. Then, antigen retrieval–treated sections were stained following the immunohistochemistry staining procedures described.

### Fos analysis pipeline

Fos images were uploaded to ImageJ (FIJI) and cropped based on brain regions outlined in the Allen Brain Atlas. The area of the cropped regions were measured and recorded. Image thresholds were set per image and particles were analyzed within the size restriction of 50-500 pixels. Fos particles were analyzed per image, and total particles of each image were divided by total area of the image. At least three Fos images per region for each mouse was quantified and averaged per mouse and per genotype (WT or Glp1r^S33W^). Ratios of NTS/AP Fos activation in Glp1r^S33W^ mice were calculated by dividing Fos/area of NTS over AP for each mouse and averaged per injection. The heatmap was generated by first normalizing each condition to WT control by region and clustered by column.

### snRNA-seq analysis

Using a previously published snRNA-seq atlas^72^, we calculated the number of Glp1r+ cells and other markers of interest and quantified their overlap, defining positive cells as mRNA counts >0.

### Behavioral Assays

#### Metabolic analysis in comprehensive lab animal monitoring system

Indirect calorimetry in the comprehensive lab animal monitoring system (CLAMS, Columbus Instruments) was used to evaluate metabolic parameters of WT and Glp1r^S33W^ mice. All WT and Glp1r^S33W^ mice were single housed and maintained on a 12:12-hour LD cycle with *ad libitum* access to food (PicoLab Rodent Diet 5053) and water. Metabolic measures of respiratory exchange ratio and energy expenditure were averaged over 3 days per mouse and per genotype. (*n* = 10/11 mice per genotype). Averaged LD cycle and total 24 hour respiratory exchange ratio and energy expenditure was analyzed per genotype and per sex.

#### Glucose tolerance tests

WT or Glp1r^S33W^ mice were overnight fasted for 16 hours prior to experiment start (zeitgeber time [ZT] 10 - ZT2). Mice received a tail snip and blood glucose measure using a Glucometer (OneTouch Ultra Test Strips for Diabetes), along with injection or oral gavage of danuglipron (3mg/kg) or liraglutide (0.3mg/kg) 15 minutes prior to dextrose (D-glucose) injection, or orforglipron (1mg/kg) 240 minutes prior to dextrose. Orforglipron was intentionally administered 4 hours prior due to its partial agonist properties, taking longer for it to act. At time point 0, mice received a blood glucose measure and injection of dextrose (1g/kg). At 15, 30, 60, 90, and 120 minutes after injection, blood glucose levels were measured.

#### Homeostatic (SD) food intake

Home cages were changed and food was removed from the home cage 1 hour prior to experiment start. Mice were injected with vehicle or drug (danuglipron 30mg/kg, liraglutide 0.3mg/kg, orforglipron 1mg/kg) at ZT11.5 (ZT8 for orforglipron) and two pellets of standard diet (PicoLab Rodent Diet 5053) were placed on the home cage floor at ZT12. Food intake measurements were taken at 1, 2, and 4 hours after ZT12 using infrared night vision goggles (Nightfox Swift Night Vision Goggles). For the danuglipron dose response experiment, 3mg/kg, 10 mg/kg, or 30 mg/kg danuglipron was injected at ZT 12 and SD intake was measured 2 hours later. For post-fast refeeding experiments (Extended Data Figure 2), mice got new bedding and 16 hours of food deprivation (ZT 10 to ZT 2) followed by a drug/vehicle injection at ZT 2 and refeeding with standard diet (30 mins after injection for danuglipron and liraglutide, 4 hours after injection for orforglipron).

#### Hedonic (HFD) food intake

Mice were habituated to high fat diet (HFD; Open Source, D12451; 4.73 kcal/gram; 45% fat, 20% protein, 35% carbohydrates; 17% sucrose) for 1 hour over two days before testing days. Standard diet was removed from the home cage 1 hour prior to experiment start. Mice were injected with vehicle or drug (danuglipron 30mg/kg, liraglutide 0.3mg/kg, orforglipron 1mg/kg) at ZT1.5 and one pellet of HFD was placed on the home cage floor at ZT2 (ZT5.5 for orforglipron). Food intake measurements were taken at 1, 2, and 4 hours after HFD delivery. The same parameters were used in oral gavage experiments with danuglipron and orforglipron.

#### Optogenetic Food Intake

For CeA^Glp1r^ soma stimulation, mice were single-housed for at least five days and habituated daily for one hour to a patch-cord and one hour to high-fat diet (HFD) pellets over two consecutive days prior to testing for the hedonic feeding paradigm. On subsequent testing days, mice underwent optogenetic stimulation at frequencies of 0 Hz (control), 5 Hz, 10 Hz, and 20 Hz for one hour each, conducted on separate days. For homeostatic feeding assessment using standard diet (SD), mice were overnight fasted, whereas for hedonic feeding assessment (HFD), mice had ad libitum access to standard chow prior to testing. Food intake was quantified manually by weighing pre-measured SD or HFD pellets at the beginning and at the end of each one-hour stimulation session. The laser stimulation protocol consisted of 473-nm blue light delivered at 20 Hz in a 1- second on, 3-second off pulse pattern. Light power exiting the fiber optic cable was measured using an optical power meter (Thorlabs) and maintained at 8-10 mW across all experiments. All experiments were performed during the light phase, between Zeitgeber Time (ZT) 3–4. Mice with missed virus injections or off-target fiber placements were excluded from the analysis.

For axon-stimulation experiments, mice expressing ChrimsonR/mCherry in either NTS^Gcg^ terminals in the CeA or CeA^Glp1r^ terminals in the VTA were single-housed for at least five days and habituated daily for one hour to a patch-cord and one hour to high-fat diet (HFD) pellets over two consecutive days prior to testing for the hedonic feeding paradigm. On different days, mice received one-hour light stimulation at 0 Hz (control), 20 Hz (CeA^Glp1r^ axons), or 40 Hz (NTS^Gcg^ axons) with a 1-second on, 3-second off pulse pattern. Simulations were bilateral for NTS^Gcg^ axons in the CeA and unilateral for CeA^Glp1r^ axons in the VTA. Light power exiting the fiber optic cable was measured using an optical power meter (Thorlabs) and maintained at 10 mW across all experiments.Homeostatic and hedonic feeding paradigms were assessed as explained above. All sessions were conducted during the light phase (ZT 3–4), and any animal with mistargeted viral expression or fiber placement was excluded from analysis.

#### Weight loss experiment

Male Glp1r^S33W^ mice >8 weeks old were placed on a high fat diet (Open Source, D12451; 4.73 kcal/gram; 45% fat, 20% protein, 35% carbohydrates; 17% sucrose) for at least 8 weeks prior to experiment start. Mice that did not gain at least 20% of their baseline body weight were excluded from testing, disqualifying female mice from this study. Male mice were randomly assigned to 21- days of vehicle or orforglipron (1mg/kg) injection and retested with the opposite treatment after 10 days of rest. Glp1r^S33W^ mice were injected daily at ZT6 and food and body weight were measured.

#### Conditioned taste avoidance (CTA)

Mice were water deprived from ZT9-ZT2 the next day and habituated to two water bottles for two days. Measurements of water bottles were taken from ZT2-ZT3 to ensure mice were drinking. On the third day, mice got two 0.15% saccharin (Saccharin, 98+%, Thermo Scientific Chemicals, catalog no. 149001000) bottles for 1 hour and immediately injected or orally gavaged with vehicle or drug (lithium chloride (125 mg/kg), danuglipron 30mg/kg, liraglutide 0.3mg/kg, orforglipron 1mg/kg). Normal water bottles were restored for the next 2 days. Water was again deprived at ZT9, and at ZT2 the next day, one water bottle and one 0.15% saccharin bottle was counterbalanced and placed in each cage. Measurements were taken after 24 hours. Preference ratios were calculated as (0.15% Saccharin consumed)/(Water consumed + 0.15% Saccharin consumed).

#### Anxiety testing

Single-housed mice were put to the behavior room to habituate one hour prior to experiment and injected with vehicle or danuglipron (30 mg/kg) at ZT 11.5 or orforglipron (1 mg/kg) at ZT 8.5. Experiments were conducted during ZT 12.5 to ZT 13.5. Mice were individually placed in the center of the arena (50.8 cm diameter with 24 cm walls) with dim light (∼25 lux) and freely explored for 5 mins. In the open field test (OFT), the arena is a circular box with a diameter of 50 cm. In the elevated plus maze (EPM) test, the arena is an elevated cross (height of 40 cm), with two closed arms (5*30 cm), and two open arms of the same dimension. Mouse movement was captured by a camera, and nose-points and center-points were tracked by Ethovision software. The arena was cleaned with 70% ethanol between trials. In the OPT, the total center area (33.9 cm diameter) was defined as the center zone. Total distance traveled and time percentage in the center zone were measured. In the EPM, head dip zone was defined as within 5 cm outside of the open arm. Total distance traveled, time spent in each arm, and head dip activity were measured in Ethovision.

#### Home cage monitoring of Glp1r^S33W^ mice

Mice were singly housed and acclimated to PhenoTyper home cages (Noldus, Netherlands) for 5 days prior to testing. Cages were maintained on a 12:12-hour light-dark cycle. During acclimation, mice had ad libitum access to standard chow (SD) in a food hopper, water bottles, running wheels, shelters, and bedding. Mice that failed to meet a baseline criterion of ≥50 food hopper head entries between ZT 12–14 were excluded to ensure sufficient engagement with the feeding setup. Of 59 mice tested, 12 did not meet this threshold and were excluded.

For testing, lithium chloride (125 mg/kg) or vehicle was administered 5 minutes prior to recording. Danuglipron (30 mg/kg) or vehicle was administered at ZT 11.5, liraglutide (0.3 mg/kg) or vehicle at ZT 10, and orforglipron (1 mg/kg) or vehicle at ZT 8. Injections were counterbalanced across animals. Behavior was recorded for 2 hours starting at ZT 12. In the “fed” condition, mice had access to a high-fat diet (HFD) placed on the cage floor from ZT 11–12. At ZT 12, HFD was removed and only SD remained in the food hopper. Fed mice were habituated to HFD exposure (1 hour/day) for 3 days prior to cage habituation. All mice were also habituated to handling and saline injections for 3 days prior to recording.

PhenoTyper sensors continuously recorded food hopper entries, water licks, and wheel rotations. Behavioral sessions were video recorded from above using infrared cameras (960×540 px, 25 fps, grayscale). Videos were cropped to 2-hour segments using Adobe Premiere Pro and re-encoded using H.264 compression with FFmpeg for consistent playback and frame indexing.

#### Machine Learning–Assisted Behavior Classification

Animal pose tracking was performed using SLEAP (v1.3.3; ^54,55^). Nine keypoints were tracked: nose, ears (L/R), tail base, and five body center points (Extended Data Fig. 4b). A total of 10,770 frames from 18 representative videos were manually labeled and split into 90% training (9,693 frames) and 10% validation (1,077 frames). The model was trained using a U-Net architecture with max stride 32, 16 filters, and ±180° rotation. Default parameters were used for the single-animal pipeline. Performance metrics included: average error = 1.59 px, OKS = 0.85, mAP = 0.81, and mAR = 0.84. Inference was run on ZT 12–14 video segments using flow tracking.

Representative heatmaps of the nose keypoint were generated by cropping to exclude cage walls and binning into 62 × 27 spatial grids. Keypoint-MoSeq^54,55^ was used to infer behavioral syllables from 80 hours of keypoint data. After alignment and centering, four latent dimensions explained 90% of variance. An autoregressive hidden Markov model (AR-HMM) with kappa = 105 (reflecting behavior timescale) identified 91 syllables. Syllables comprising ≥0.5% of frames were included. Those between 0.01–0.5% were retained; syllables <0.01% were excluded. Syllables were grouped into behavioral categories (Extended Data Fig. 6) by two trained raters. Some syllables captured blended actions (e.g., "groom/sniff"), likely due to overhead view limitations. Low-quality syllables (∼1%)—often under the wheel—were excluded.

#### Behavior Localization and Categorization

The locations of the food hopper, water spout, and shelter were identified using OpenCV (<u>OpenCV, 2024</u>), and corresponding regions of interest (ROIs) were defined. The nose keypoint was used to detect entries into the food and water ROIs; center-point 3 was used for shelter entry. Behaviors were labeled contextually (e.g., “sniff by food” vs “sniff” elsewhere). This produced 22 distinct behaviors, grouped into 5 broader categories based on behavioral similarity and transition frequency: *rest/groom in shelter*, *groom*, *move/explore*, *food-motivated*, and *drink*. In some analyses (Extended Data Fig. 7C, Extended Data Fig. 8K, 8Q, 8W), movement within the shelter was analyzed separately. “Pause” outside the shelter was excluded due to lack of behavioral relevance. These behaviors captured 95.7% of all behavioral time across videos. For each 2-hour recording, time spent per behavioral category was normalized to total behavioral time.

### Behavior Analysis

Behavioral proportions were analyzed using beta-distributed generalized linear mixed-effects models, with mouse as a random effect. Different link functions (e.g., logit, cloglog) were used depending on the distribution of each behavior. Proportions in the control condition were normalized to mean = 1; treatment data for each animal were scaled using this normalization to highlight magnitude of change. For network analysis, transition probabilities between behaviors were computed per mouse and normalized by total transitions from the starting behavior. Transitions were averaged per group. Line thickness in network plots reflects groupwise maximum-normalized probabilities; node size corresponds to average bout length and was scaled manually in Illustrator.

To explore full dataset differences (duration, bouts, transitions, sensors, distance), we used UMAP with PCA preprocessing (16 PCs explaining 95% of variance). UMAP was fit with 3 components. Distances between condition centroids were visualized as heatmaps and dendrograms. To link features to the LiCl signature, Spearman correlations were calculated between each behavioral feature and the Euclidean distance to the LiCl group centroid in UMAP space. FDR correction was applied; only significant correlations were shown. Signs were flipped so positive values reflect greater similarity to LiCl. All features were z-scored prior to PCA to ensure equal weighting across variables. UMAP was implemented using the *scikit-learn* library^92,93^, and behavioral transition networks were generated with *NetworkX*^94^. All analysis code is available on <u>GitHub</u>.

### Electrophysiology Recordings

#### Brain slice preparation

Preparation of acute brain slices for patch-clamp electrophysiology experiments was modified from standard protocols previously described^95,96,97^. Mice were anesthetized with isoflurane and decapitated. The brains were rapidly removed and kept in chilled artificial cerebrospinal fluid (ACSF) (1°C) containing (in mM): 125 NaCl, 2.5 KCl, 1.25 NaH_2_PO_4_, 2 CaCl_2_, 1 MgCl_2_, 0.5 L-ascorbic acid, 10 glucose, 25 NaHCO_3_, and 2 Na-pyruvate (osmolarity 310 mOsm). Slices were continuously oxygenated with 95% O2 and 5% CO2 throughout the preparation. Coronal brain sections, 300 μm, were prepared using a Leica Microsystems VT1200 vibratome. Slices were collected and placed in ACSF warmed to 37°C for 30 minutes and then kept at room temperature for up to 5 hours.

#### Recordings

Brain slices were placed in a chamber superfused (∼3 mL/min) with continuously oxygenated ACSF solution warmed to 32 ± 1°C. hGLP1R-expressing amygdala neurons were identified by video microscopy based on the expression of eYFP and regional marker. Whole-cell electrophysiology recordings were performed using a Multiclamp 700B amplifier with signals digitized by a Digidata 1550B digitizer. Currents were amplified, lowpass-filtered at 2 kHz, and sampled at 35 kHz. Borosilicate electrodes were fabricated using a Brown-Flaming puller (model P1000, Sutter Instruments) to have pipette resistances between 2.5 and 4.5 mΩ. Current-clamp recordings of membrane potentials were collected in an ACSF solution identical to that used for preparation of brain slices. The internal solution contained the following (in mM): 120 K-gluconate, 10 NaCl, 2 MgCl_2_, 0.5 K_2_EGTA, 10 HEPES, 4 Na_2_ATP, and 0.3 NaGTP, pH 7.2 (osmolarity 290 mOsm). Resting membrane potential was recorded as previously described (1,2). After 5 mins of baseline membrane potential recordings, 30 μm of danuglipron was perfused. Danuglipron powder (Selleck, catalog no. S9851) was dissolved to 30 mg/mL in 100% ethanol with gentle sonication, then diluted to 3 mg/mL (5.4 mM) in vehicle [1 N NaOH, 2% Tween 80, 5% polyethylene glycol (PEG) 400, 5% dextrose]. For electrophysiology recordings, 280 µL of this stock was added to 50 mL ACSF (1:180 dilution) to yield a final danuglipron concentration of 30 µM, corresponding to final concentrations of approximately 0.056% ethanol, 0.011% Tween 80 and 0.028% PEG 400.

### Statistics

Electrophysiology recordings were analyzed using ClampFit 11.2. All statistical comparisons were made using the appropriate test in GraphPad Prism 9.5.0. For membrane, AP properties, and amplitude underwent descriptive statistics followed by normality and lognormality test using gaussian distribution. Data were assessed for normality using the D’Agostino-Pearson omnibus normality test, Anderson-Darling test, Shapiro-Wilk test, and Kolmogorov-Smirnov test with Dallal-Wilkinson-Lillie for P-values. Data were tested for outliers using the ROUT method, and statistical outliers were not included in data analysis. Followed by Tukey’s test with no gaussian distribution creating a nonparametric t-test using Wilcoxon matched-pairs signed rank test to compare responses caused by the drug. Data are presented as individual data points and/or mean ± SEM.

### In Vivo Fiber Photometry Recordings Calcium recordings (GCaMP)

Mice underwent 20-minute daily habituation sessions over two consecutive days to acclimate to the fiber-optic cable (Doric Lenses, Ø400-μm core, 0.57 numerical aperture). On the test day, mice were injected with either vehicle, danuglipron (30 mg/kg) or liraglutide (0.3 mg/kg) two hours prior to recording. The order of injections was randomized to avoid order effects. Following the two-hour post-injection period, mice were connected to patch cables which were interfaced with rotary joints to enable free movement. Recordings were conducted for one hour in mice home cage without food and water. Fiber photometry data were recorded using fluorescent signals from both calcium-dependent (465 nm) and calcium-independent isosbestic (405 nm) excitation wavelengths (Doric). The isosbestic (405 nm) signal served to control for artifacts. The light power of the fiber optic cable was measured before each experiment and maintained at approximately 20-30 μW for both the calcium-independent isosbestic (405 nm) and calcium-dependent (465 nm) signals.

### Calcium analysis (GCaMP)

The isosbestic signal (405 nm) was fitted to the calcium-dependent (465 nm) signal using a linear least squares method implemented in a custom MATLAB script and ΔF/F was calculated as (465 nm – fitted 405 nm) / fitted 405 nm. For z-score calculation, we then implemented a paired z-score normalization: for each mouse, we used the full ΔF/F time series from its vehicle session to calculate μ_(vehicle)_ (mean) and σ_(vehicle)_ (SD), and all ΔF/F values—both vehicle and drug—were converted to z-scores via [ΔF/F(t) – μ_(vehicle)_] / σ_(vehicle)_, anchoring comparisons to a common baseline distribution. Significant calcium transients were detected on these z-scored traces using a threshold of median + 2 SD (of the entire recording) with a minimum duration of 1.5 s; events were counted per trial and displayed as heatmaps of z-score to ensure full transparency of raw recording structure. Finally, the area under the curve of each mouse’s complete z-scored trace was calculated using a custom MATLAB script to validate overall activity differences between conditions. Heatmaps were generated in MATLAB using Min-Max normalization, scaled to a range of 0–1. For each mouse, the normalization range was determined based on the Vehicle condition: the average of the lowest 360 data points was set as the minimum, and the average of the highest 360 data points was set as the maximum. This normalization range, derived from the Vehicle condition, was then applied to the corresponding paired Danuglipron or Liraglutide data for the same mouse. A moving average with a window and bin size of 10 smoothed the data, which was then plotted as a heatmap. Mice with missed virus injection or off-target fiber placement were excluded from analysis.

### Dopamine recordings (dLight)

Mice (WT, Glp1r-Cre or Glp1r^S33W^) were single-housed and habituated to the fiber optic cable and high-fat diet (HFD) for one hour over two consecutive days. On the test day, mice received an injection of either a drug (liraglutide [0.3 mg/kg], danuglipron [30 mg/kg], or orforglipron [1mg/kg]) or vehicle on different days, with the order of drug versus vehicle injections randomized. Liraglutide and danuglipron were administered two hours before recording, while orforglipron was given four hours prior. Fiber photometry data were recorded as described above. Fluorescent signals were collected from both dopamine-dependent (465 nm) and dopamine-independent isosbestic (405 nm) excitation wavelengths. During the testing sessions, small pellets of HFD (∼10 mg) were dropped into a cup at two-minute intervals after the mice retrieved the pellet. 5 to 6 trials were conducted per mouse. The recording session was video recorded to timestamp food retrieval time and recordings were conducted during the light phase, between ZT3 and ZT6.

### Dopamine analysis (dLight)

The isosbestic signal (405 nm) was fitted to the dopamine-dependent (465 nm) signal using a linear least squares method implemented in a custom MATLAB script. Then ΔF/F was calculated as (465 nm – fitted 405 nm) / fitted 405 nm. To account for inter-animal differences in signal intensities, Z-scores were calculated for the ΔF/F signals. The baseline period for each food trial was defined as the 30-second interval prior to food retrieval. The mean and standard deviation of the baseline period were used to compute the Z-scores, with the formula: *Z* score = (F- Fμ(baseline))/std(baseline), where F is the 405 nm corrected 465 nm signal (ΔF/F), μ(baseline) is the mean, and std is the standard deviation of the baseline period. Video frames were analyzed to determine the exact timestamp when the mouse retrieved the pellet, which was defined as time 0 for each retrieval. The 30-second window centered around the food retrieval time was extracted. The area under the curve (AUC) and maximum fluorescence Z-scored within the food retrieval window was further extracted and analyzed for quantification of dopaminergic activity. 5 food trials were averaged per mouse. Mice with missed virus injections or off-target fiber placements were excluded from the analysis.

### Statistical analyses

All data are presented as mean ±SEM unless otherwise noted. Statistical tests including paired or unpaired two-tailed t-tests, Kruskal-Wallis tests, Wilcoxon signed-rank tests, Pearson regression, one-way ANOVA, two-way/repeated-measures ANOVA (with Bonferroni correction or Tukey’s HSD post hoc tests), MANOVA (with Bonferroni correction post hoc test), linear mixed effect models with beta regression (with Holm post hoc test) were performed using RStudio (v4.3.0), Python (v3.11.5), JupyterLab (v3.6.3), MATLAB (R2023a), or GraphPad Prism (v10.4.0). Brief descriptions of all experiments in each figure panel, sample sizes, mean ±SEM, statistical test, test statistics, and *P*-values are presented in Supplementary Table 1. \**P* <0.05; \*\**P* <0.01; \*\*\**P* <0.001.

## Data availability

All data will be available on Dryad upon publication and select code will also be available on Github at https://github.com/UVACircMetNeuLab.

## Acknowledgements

We thank the members of the Güler, Deppmann, Campbell, and Provencio laboratories (University of Virginia) for comments and suggestions on the preparation of the manuscript. We are thankful for technical help from Stefani A. Mancuso and technical advice from Amrita Pathak, along with Anthony Spano (University of Virginia) who developed the three viral constructs. We thank Talmo Pereira (Salk Institute for Biological Studies) and Caleb Weinreb (Harvard Medical School) for their technical advice. We thank Songwei He and Ruediger Klein (Max-Planck Institute for Biological Intelligence) for sharing their processed single cell RNA-seq CeA dataset. We thank the Genetically Engineered Murine Model (GEMM) Core (University of Virginia) for aiding in the CRISPR-Cas9 development of our mouse model.

## Author contributions

Conceptualization: ENG, TBG, IRS, ABK, JNC, CDD, ADG Data curation: ENG, TBG, IRS, TCJD, SO, YZ, ODL, GT, OK

Formal analysis: ENG, TBG, IRS, TCJD, ADG

Funding acquisition: ENG, TBG, IRS MKP, JNC, CDD, ADG

Investigation: ENG, TBG, IRS, AKB, ODL, YZ, YS, TCJD, GT, OK, MC, NJC, ANW, OYC, SO, WL, AA, KM, KIW, SML, AK, GG

Methodology: ENG, TBG, IRS, ABK, AKB Project administration: ADG

Resources: KTB, LSZ, MKP, JNC, CDD, ADG

Supervision: ENG, TBG, IRS, KTB, LSZ, MKP, JNC, CDD, ADG Validation: ENG, TBG, IRS, AKB, CDD, ADG

Visualization: ENG, TBG, IRS

Writing – original draft: ENG, TBG, IRS, CDD, ADG

Writing – review & editing: ENG, TBG, IRS, JNC, CDD, ADG

## Extended Data

**Extended Data Fig. 1.**
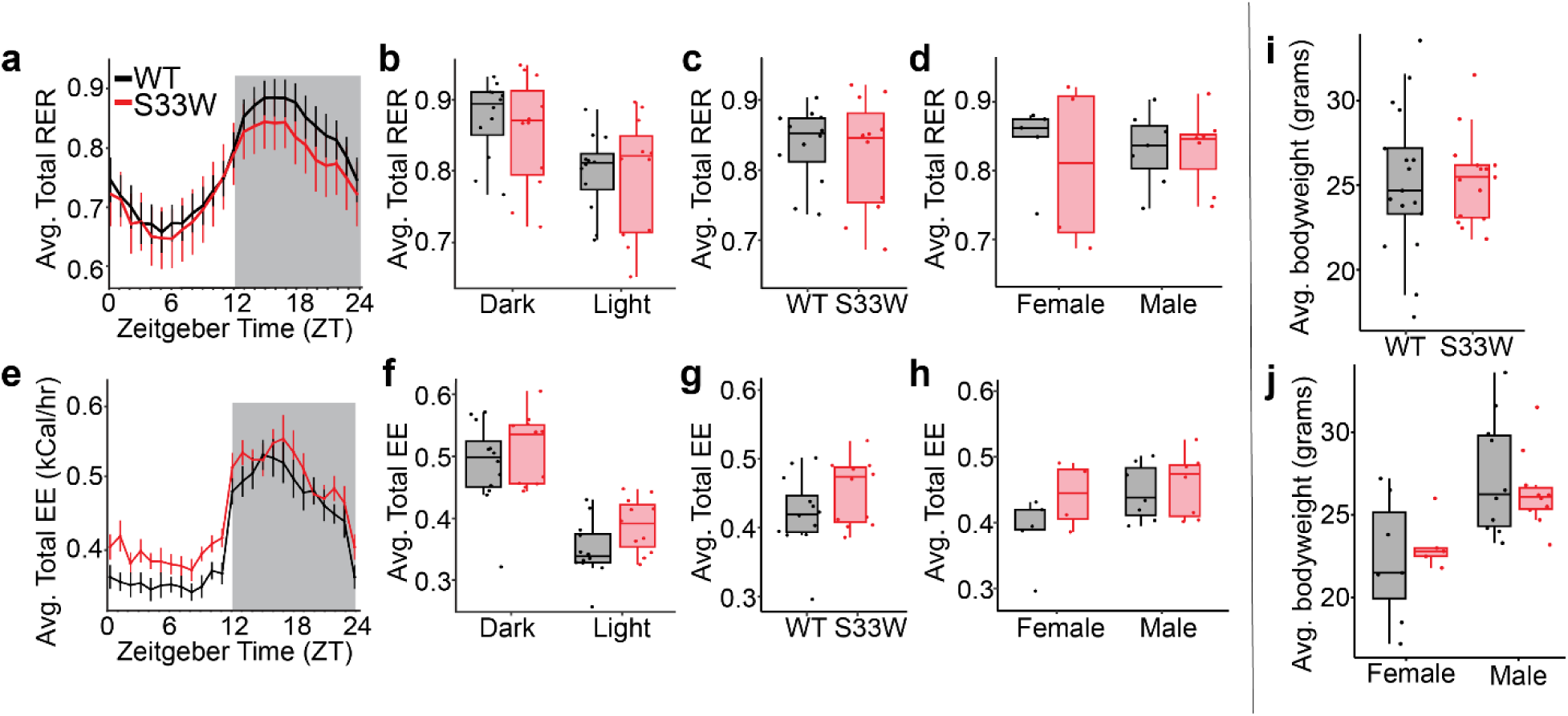
**Metabolic profiling of WT and Glp1r^S33W^ mice. a,e**, Diurnal rhythms averaged over 3 days of (**a**) respiratory exchange ratio (RER) and (**e**) energy expenditure (EE) in WT and Glp1r^S33W^ mice. **b-g**, Average dark/light phase and total 24 hour (**b,c**) RER and (**f,g**) EE between WT and Glp1r^S33W^ mice (*n* = 11-12 per genotype, two-way ANOVA with Bonferroni correction or Welch’s t-test). **d,h**, Total 24 hour (**d**) RER and (**h**) EE averaged by sex (*n* = 4-5 females, *n* = 7 males per genotype, two-way ANOVA with Bonferroni correction). **i**, Average baseline body weight of WT and Glp1r^S33W^ mice 10 to 20 weeks old (*n* = 18-19 per genotype, Welch’s t-test). **j**, Average baseline body weight of WT and Glp1r^S33W^ male and female mice (*n* = 8-9 females, *n* = 10 males per group, two-way ANOVA with Bonferroni correction). Data are represented as medians ± Q1-Q3. \**P*<0.05; \*\**P*<0.01; \*\*\**P*<0.001.

**Extended Data Fig. 2.**
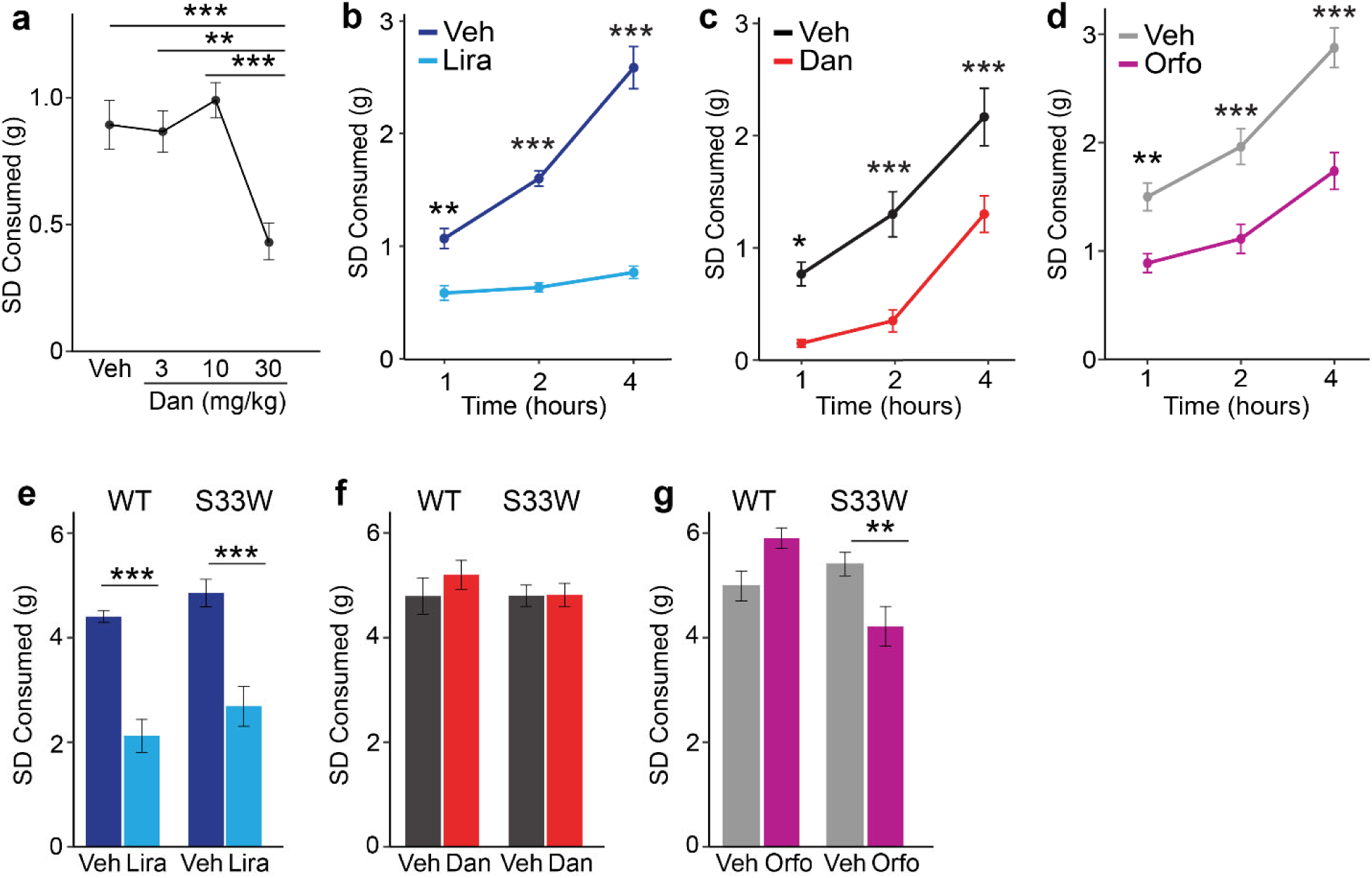
24-hour and post-fast refeed effects of GLP1RAs on standard diet consumption. **a**, 2-hour SD intake (ZT 12 - ZT 14) after vehicle (Veh) or escalating doses of danuglipron (3mg/kg, 10 mg/kg, or 30 mg/kg) (n = 8-15 per group, one-way ANOVA with Tukey’s HSD, \*\**P*<0.01; \*\*\**P*<0.001). **b-d**, Post-fast refeeding of SD intake 1, 2, and 4 hours post-treatment of (**b**) Lira (*n* = 6) or (**c**) Dan (*n* = 6) (**d**) Orfo (*n* = 8) and vehicle controls in Glp1r^S33W^ mice (two-way ANOVA with Bonferroni correction, \**P*<0.05; \*\**P*<0.01; \*\*\**P*<0.001). **e-g**, Standard diet (SD) consumption 24 hours after (**e**) liraglutide, (**f**) danuglipron, and (**g**) orforglipron injection in WT and Glp1r^S33W^ mice. (*n* = 6 per injection, two-way ANOVA with Bonferroni correction, \*\**P*<0.01; \*\*\**P*<0.001). Data are represented as means ± SEM.

**Extended Data Fig 3.**
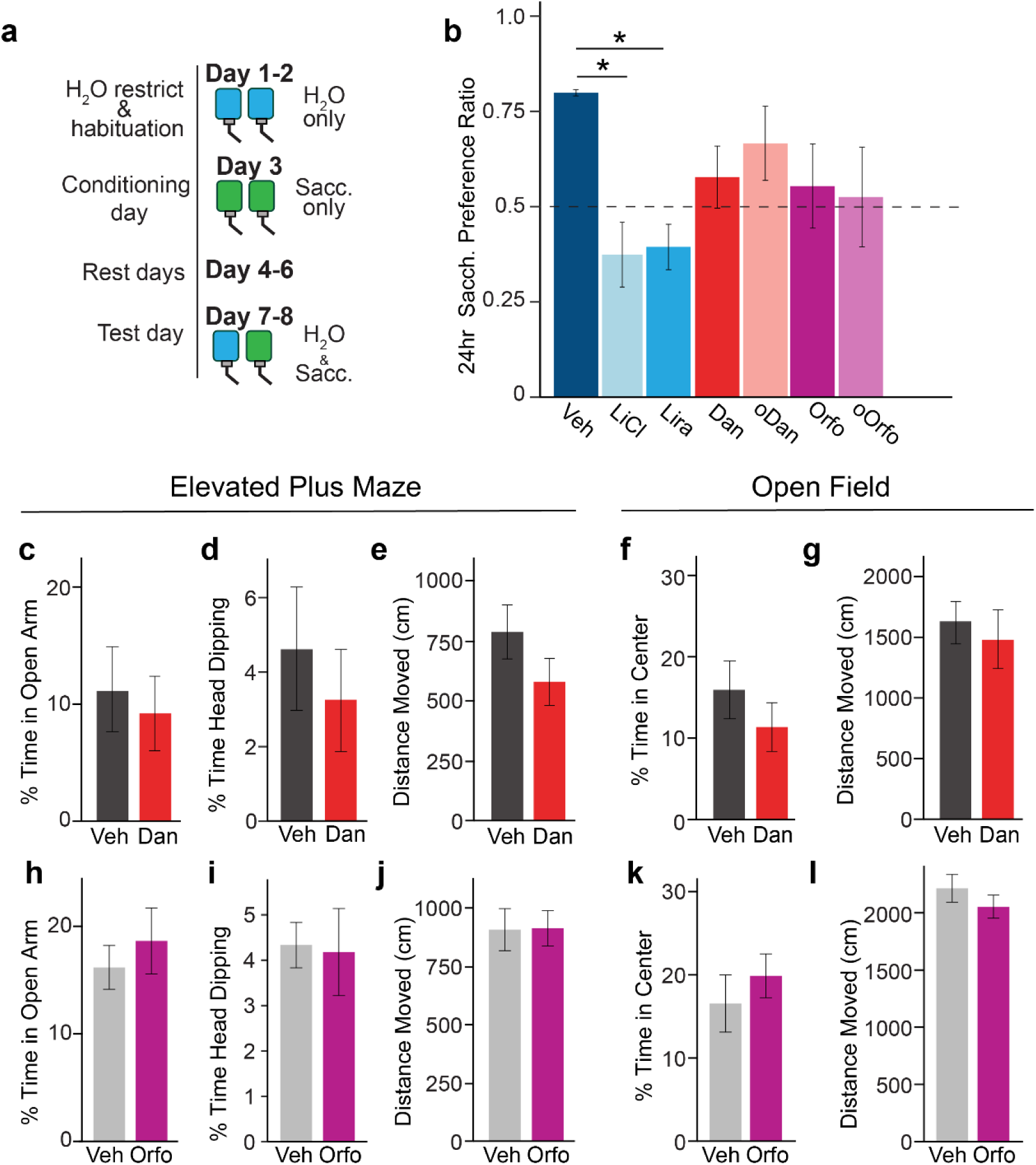
GLP1RAs effect on conditioned taste avoidance (CTA) and anxiety measures. **a**, Pipeline for CTA protocol. **b**, 24-hour saccharin preference after injections of vehicle (Veh), lithium chloride (LiCl), liraglutide (Lira), danuglipron (Dan), oral Dan (oDan), orforglipron (Orfo) or oral Orfo (oOrfo) (*n* = 6-8 per injection, one-way ANOVA with Tukey’s HSD correction, \**P*<0.05).**c-l**, Elevated plus maze and open field anxiety tests after injections of (**c-g**) vehicle (Veh) or danuglipron (Dan) and (**h-l**) vehicle (Veh) or orforglipron (Orfo). Glp1r^S33W^ mice were injected with Veh/Dan at ZT 11.5 or Veh/Orfo at ZT 8.5 and tested at ZT 12.5. For elevated plus maze, the percentage of time spent in open arms, percentage of time spent head dipping, and distance traveled were measured. For the open field test, the percentage of time spent in the center (defined as 4/9 of center area) and distance traveled were measured (*n* = 8 per injection, paired t-test). Data are represented as means ± SEM.

**Extended Data Fig. 4.**
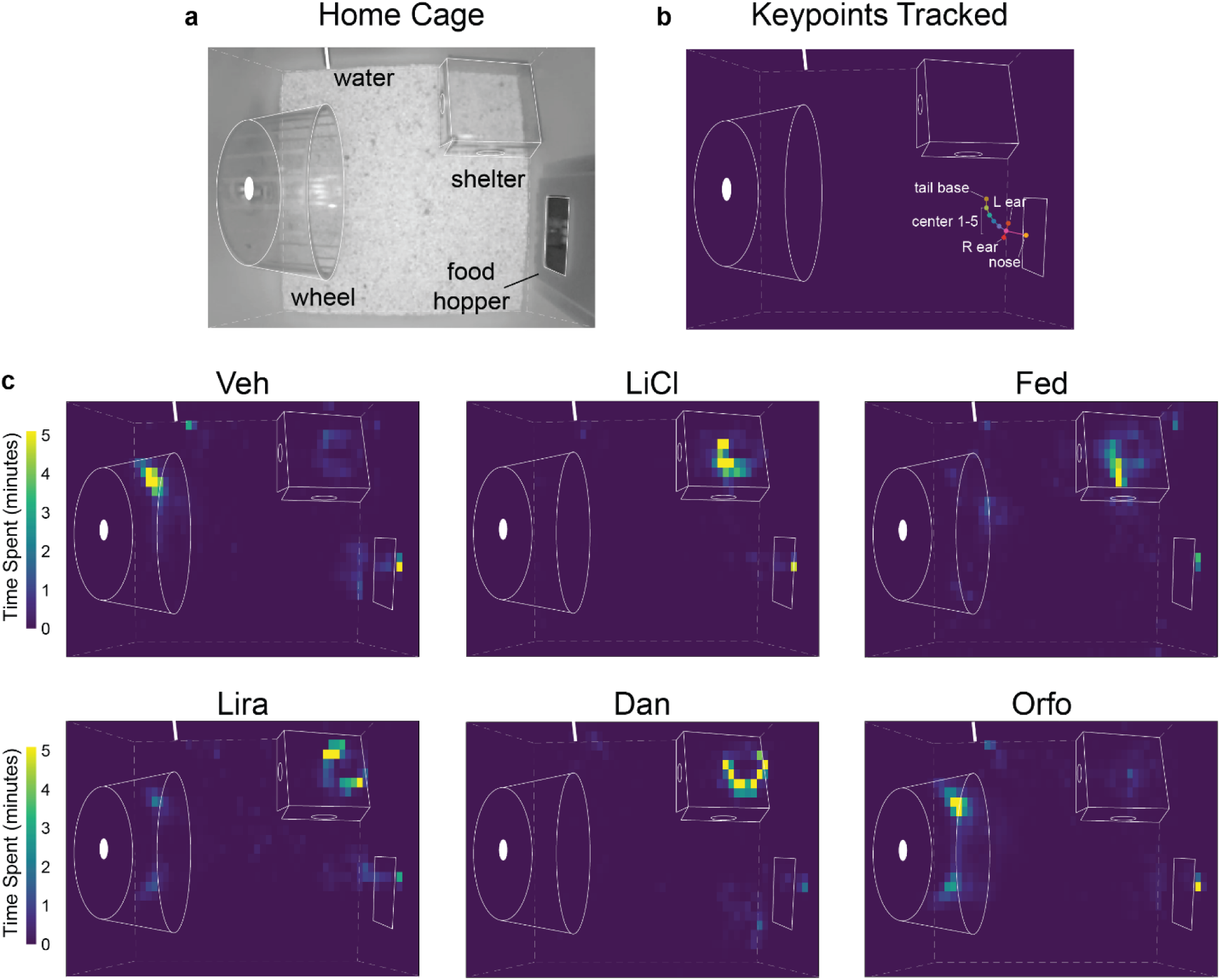
Home cage monitoring heatmaps of Glp1r^S33W^ mice. (**a**), Representative image of home cage set-up (dimensions: 30 x 30 x 43.5 cm) with calibrated positions of wheel, IR translucent shelter (10.5 x 10.5 x 6 cm), food hopper (5.5 cm from floor), and water bottle spout (5 cm from floor) identified in white outline (top). (**b**) 9 Keypoints tracked on mice using SLEAP: nose, ears (L/R), tail base, and five body center points. (**c**), Coordinate-based heatmaps of nose keypoint trajectories recorded at 25 fps over 2-hour sessions for each treatment (ZT 12-14). Color scale represents dwelling time in minutes.

**Extended Data Fig. 5.**
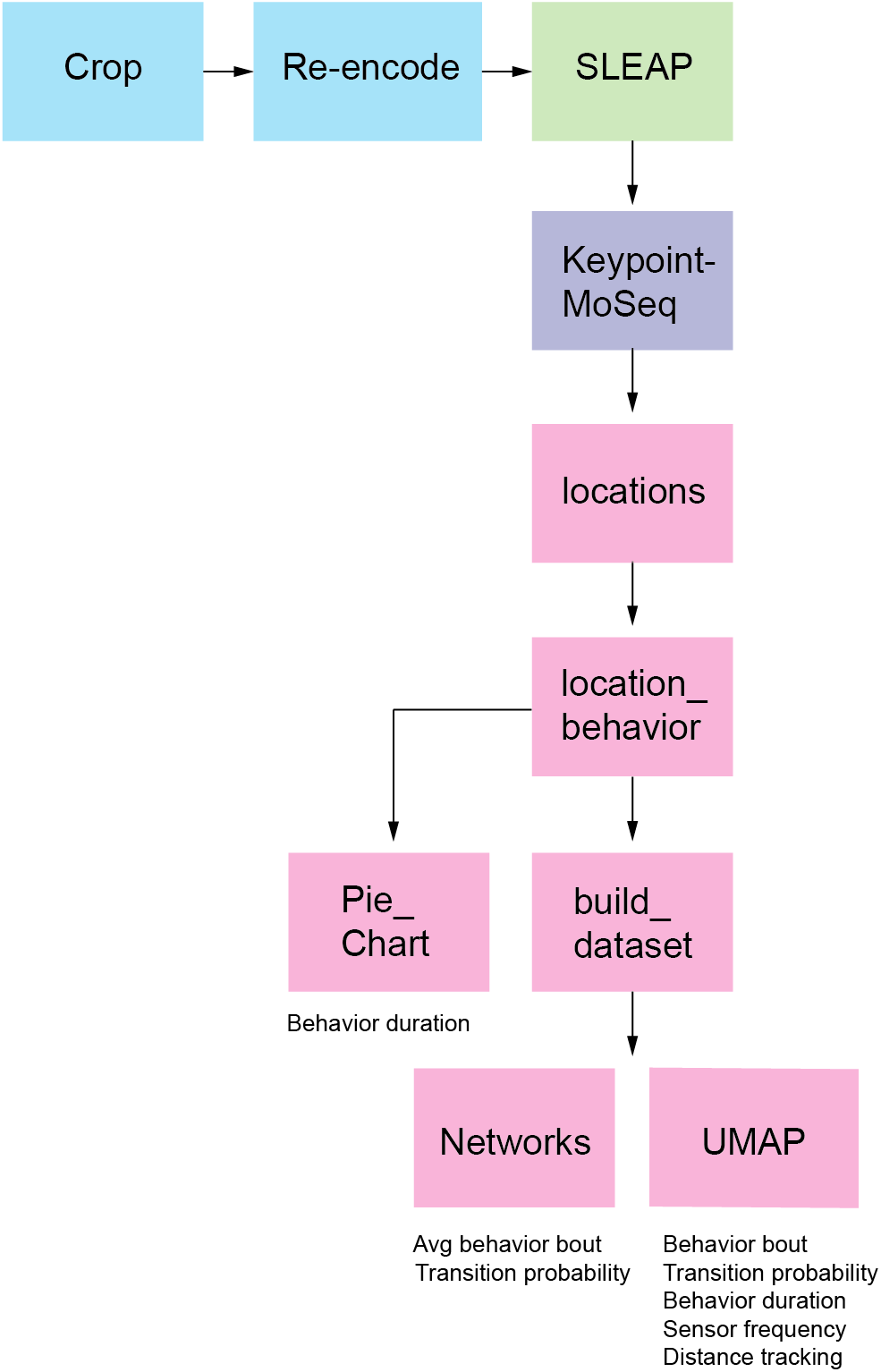
Machine-learning assisted pipeline for home cage behavioral analysis. Blue squares represent video pre-processing steps. Social LEAP Estimates Animal Poses (SLEAP) was used to extract nine keypoint coordinates for each mouse in the videos (Fig. 2a**-b****, Extended Data** Fig. 4b-c). Keypoint Motion Sequencing (Keypoint-MoSeq) identified behavioral syllables from these coordinates (**Extended Data** Fig. 6). Pink squares represent Python scripts used for the analysis of Keypoint-MoSeq syllable results: locations (identify regions of interest), location_behavior (Fig. 2b), build_dataset, Pie_Chart (Fig. 2d**-h**), Networks (Fig. 2n**- r**), and UMAP (Fig. 2s**-u**).

**Extended Data Fig. 6.**
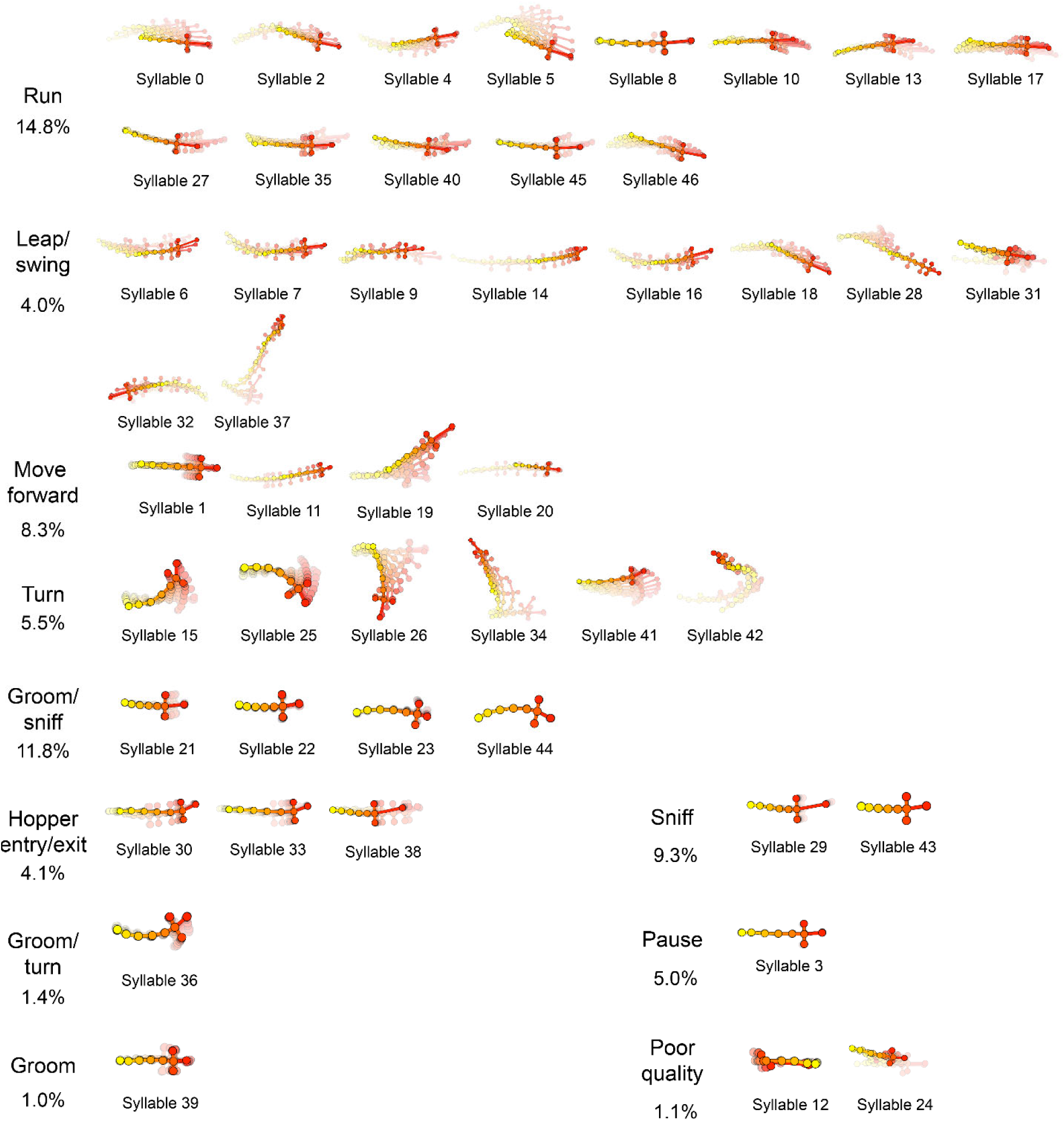
Trajectory plots of main behavioral syllables identified by Keypoint-MoSeq. Syllables identified from keypoint coordinates by Keypoint-MoSeq were sorted into 11 categories by trained raters using grid movies generated for each syllable. The proportion of each syllable category is presented as a percentage. Rare syllables (minimum frequency < 0.5% but ≥ 0.01%) are not visually depicted but were included in the analysis.

**Extended Data Fig. 7.**
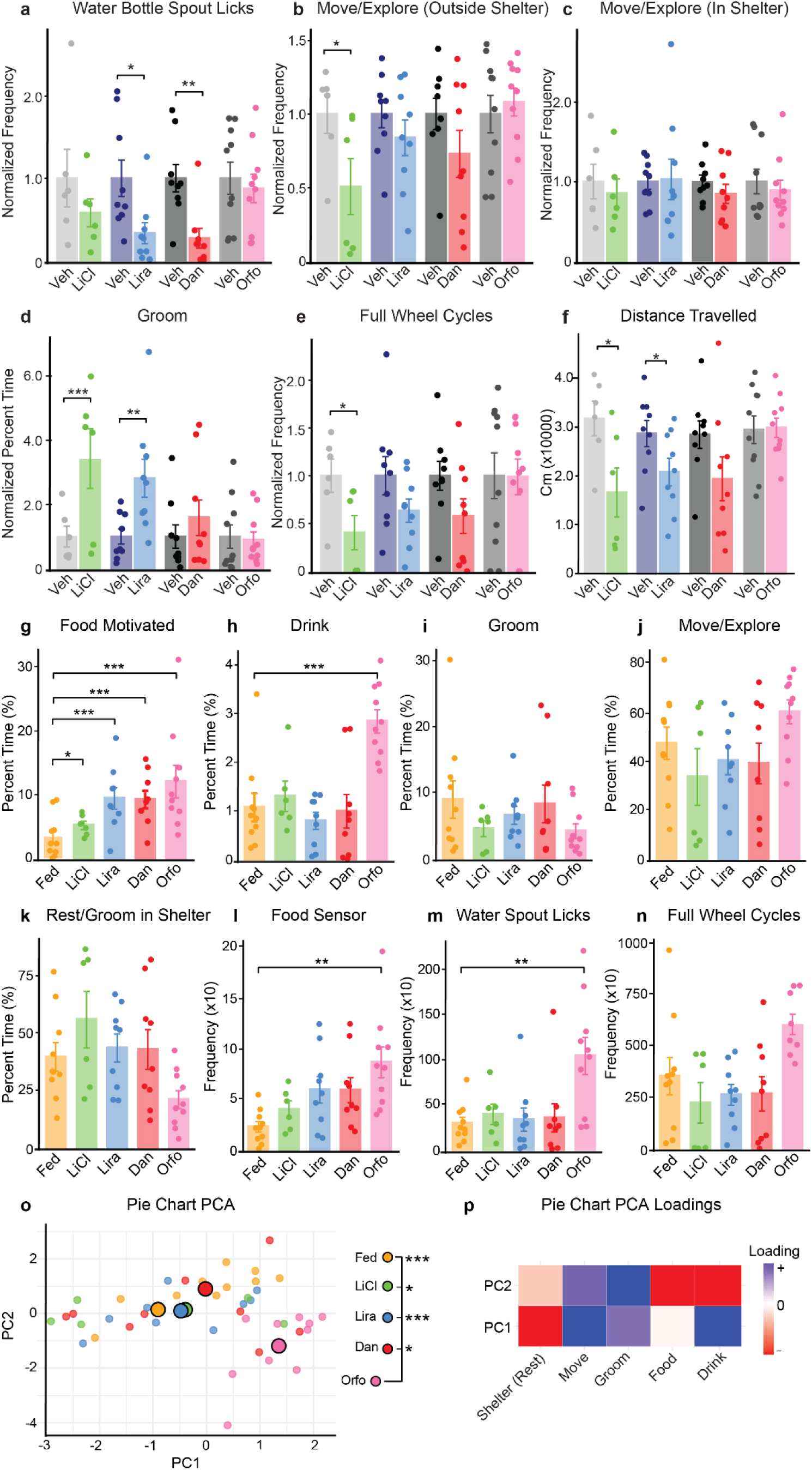
Percent time engaged in behaviors, sensor activity, and PCA of pie chart behavior profiles. **a**, Sensor data normalized to paired controls: Total number of water spout licks (frequency). (paired *t*-test on raw frequency data; \**P* < 0.05; \*\**P* < 0.01). **b–d**, Percent time spent performing each behavior, normalized to paired control groups for liraglutide-, danuglipron-, orforglipron-, and LiCl-treated mice. (*n* = 6 Veh/LiCl, *n* = 9 Veh/Lira, *n* = 9 Veh/Dan, *n* = 10 Veh/Orfo, generalized linear mixed-effects model (GLMM) with beta regression on raw (non-normalized) proportion data, comparing treatment to control with random intercept for mouse ID (paired); \**P* < 0.05; \*\**P* < 0.01; \*\*\**P* < 0.001). **e**, Sensor data normalized to paired controls: Total number of complete wheel rotations. (paired *t*-test on raw frequency data; \**P* < 0.05; \*\**P* < 0.01). **f**, Total distance travelled (centimeters) tracked using the neck/upper back keypoint of mice administered LiCl, danuglipron (Dan), liraglutide (Lira), orforglipron (Orfo), or their paired controls (Veh). (Paired t-test, *n* = 6 Veh/LiCl, *n* = 9 Veh/Dan, *n* = 9 Veh/Lira, *n* = 10 Veh/Orfo, ; \**P* < 0.05). **g–k**, Percent time spent performing behaviors across unpaired treatment groups. (*n* = 10 fed, *n* = 6 LiCl, *n* = 9 Lira, *n* = 9 Dan, *n* = 10 Orfo, GLMM with beta regression on raw proportion data, comparing fed group to each treatment group with Holm-corrected post hoc tests; \**P* < 0.05; \*\*\**P* < 0.001). **l–n**, Sensor-based activity: **l**, Total number of head entries at the food hopper (TTL beam breaks); **m**, Total number of licks from the water spout; **n**, Total number of complete wheel turns. (one-way ANOVA with post hoc comparisons of each group vs fed, Holm correction applied; \*\**P* < 0.01). **o–p**, Principal component analysis (PCA) of behavior profiles (used for pie chart data): **o**, PCA plot where small dots represent individual mice and centroids represent group means. **p**, PCA loadings for PC1 and PC2; color intensity reflects behavioral contribution (blue = positive loading, red = negative loading). Group differences in PCA space were assessed using pairwise MANOVAs on PC1 and PC2 with Bonferroni correction (\**P* < 0.05; \*\*\**P* < 0.001). All data are presented as mean ± SEM.

**Extended Data Fig. 8.**
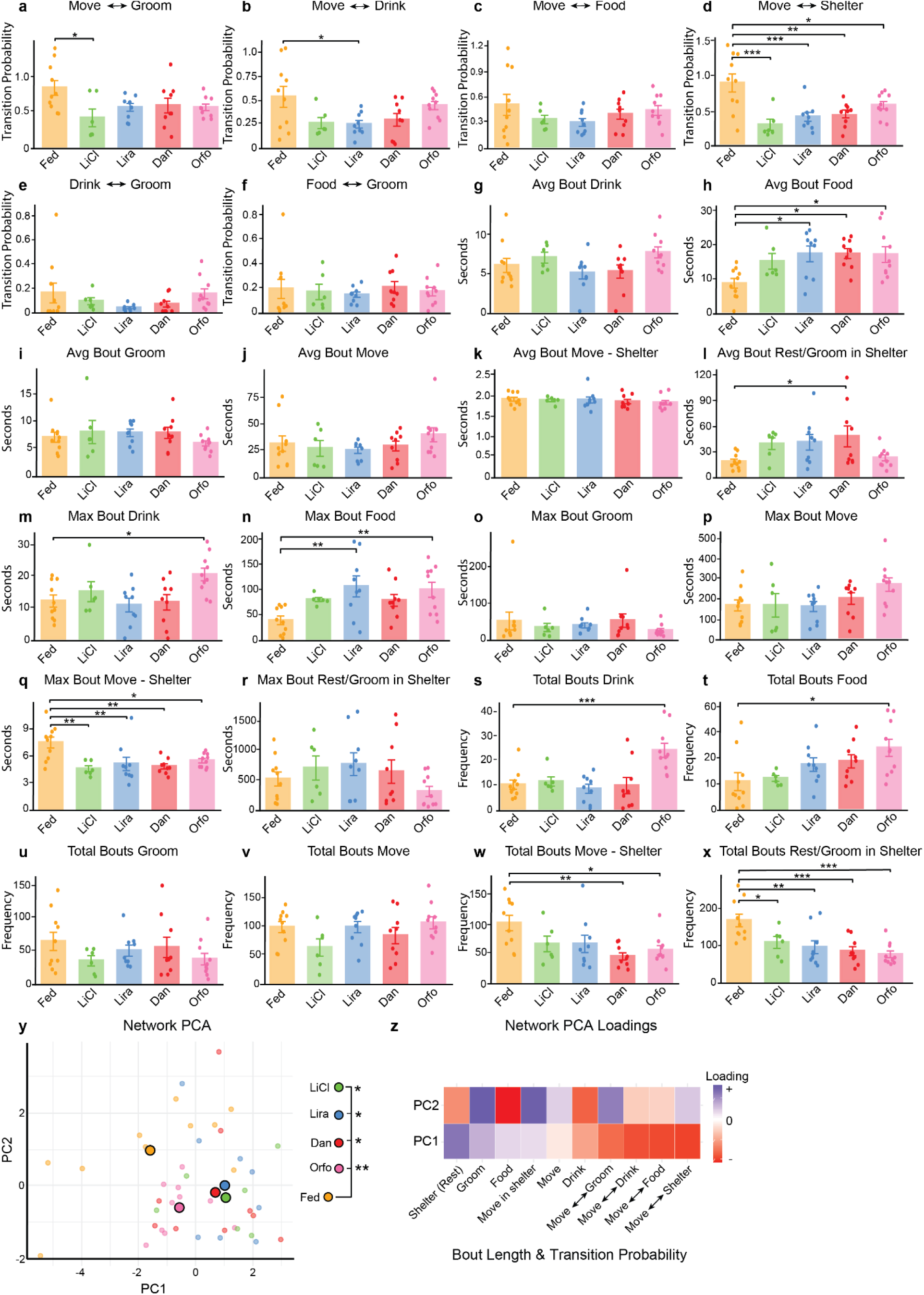
**Behavioral transitions, bout characteristics, and network PCA across treatment groups. a–f**, Transition probabilities between behaviors for fed, LiCl-, liraglutide-, danuglipron-, or orforglipron-treated mice. Directionality of transitions was combined for simplicity. Rare transitions (<0.1 probability) were excluded from analysis, with the exception of *move/explore → drink*, which was retained due to behavioral relevance. Statistical comparisons were performed using one-way ANOVA with Holm-corrected post hoc pairwise tests comparing fed mice to each treatment group (*n* = 10 fed, *n* = 6 LiCl, *n* = 9 Lira, *n* = 9 Dan, *n* = 10 Orfo, \**P* < 0.05; *\*\*P* < 0.01; \*\*\**P* < 0.001). **g–l**, Average bout length (seconds) for each behavior across conditions. **m–r**, Maximum bout length (seconds) observed per mouse during the recording period. **s–x**, Total number of behavior bouts (frequency) across conditions. Statistical comparisons for panels g–x used one-way ANOVA with Holm correction as above. **(y)**, Principal component analysis (PCA) of behavioral networks combining transition probabilities and average bout lengths. Each point represents an individual mouse; group centroids reflect mean behavioral profiles. **(z)**, PCA loadings for PC1 and PC2; color intensity reflects strength of contribution (blue = positive loading, red = negative loading). Behavior abbreviations: *food* = food motivated behavior, *move* = move/explore, *shelter* = resting and grooming in shelter. *Note:* The behavior *move/explore in shelter* was excluded from transition calculations to improve interpretability, as it often represents a transient state between shelter and other behaviors. Group differences in PCA space were assessed by conducting pairwise MANOVAs comparing the fed group to each treatment group on PC1 and PC2, with Bonferroni correction applied (\**P* < 0.05; \*\**P* < 0.01). All data are presented as mean ± SEM.

**Extended Data Fig. 9.**
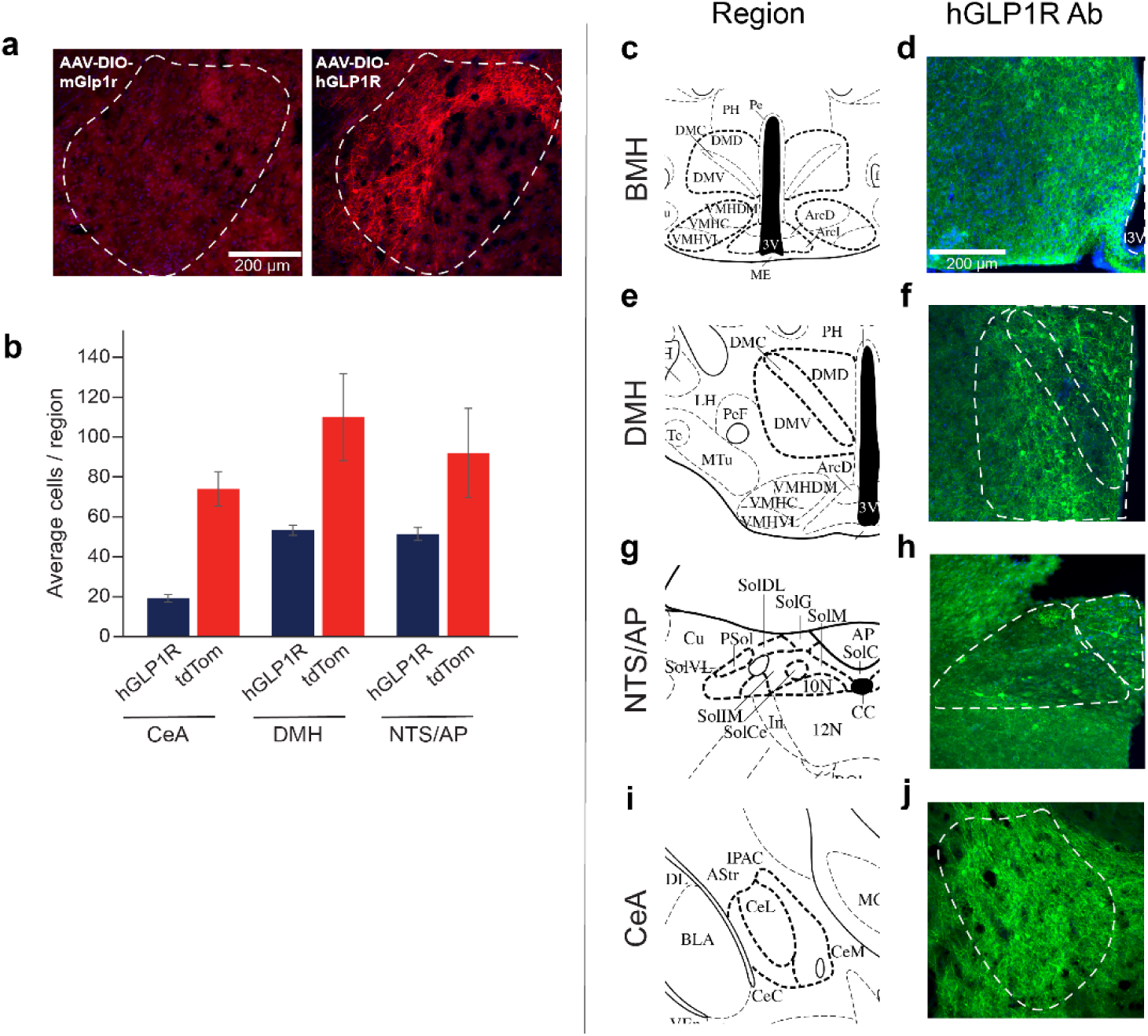
Validation of the AAV-hSyn-DIO-hGLP1R virus. **a**, Validation of human GLP1R (hGLP1R; red) expression in the CeA of Glp1r-Cre mice injected with AAV-DIO-hGLP1R (right). hGLP1R protein expression is absent in Glp1r-Cre mice injected with AAV-DIO-mGlp1r in the CeA (left). Scale bar = 200 µm. **b**, Quantification of hGLP1R+ cells and average number of Glp1r+ cells (measured in Glp1r-Cre;tdTomato mice) in the CeA, DMH, or NTS/AP regions. **c-j,** Fluorescent microscopy images of the brain regions stereotaxically injected with AAV-DIO- hGLP1R. Images from Allen Brain Atlas, with regions of interest in bold and hGLP1R antibody staining (green) in the (**c,d**) basomedial hypothalamus (BMH), (**e,f**) DMH, (**g,h**) NTS/AP, or (**i,j**) CeA. Scale bars = 200 µm.

**Extended Data Fig. 10.**
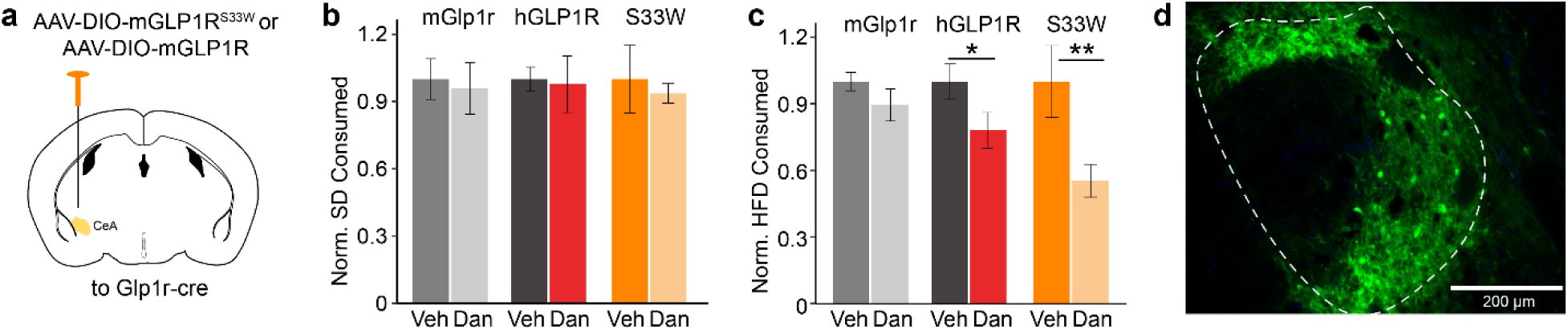
AAV-DIO-mGLP1R^S33W^-HA targeted to the CeA of Glp1r-Cre mice recapitulates effects of AAV-DIO-hGLP1R. **a**, Schematic of Glp1r-Cre mice injected with a Cre-dependent AAV carrying full length mouse Glp1r (mGlp1r) or AAV carrying full length mouse GLP1R with the S33W mutation at position 33 (mGLP1R^S33W^) in the CeA. **b,c**, Normalized 4-hour (**b**) SD and (**c**) HFD consumption post-injection of vehicle or danuglipron in mGlp1r, full length human GLP1R (hGLP1R) or mGlp1r-S33W (S33W) expressing mice (*n* = 4-9 per group, two-way ANOVA with Bonferroni correction, \**P*<0.05; \*\**P*<0.01). **d**, Representative image of AAV-DIO- mGLP1R^S33W^-HA expression in the CeA (green). Scale bars = 200 µm. Data are represented as means ± SEM. \**P*<0.05; \*\**P*<0.01; \*\*\**P*<0.001.

**Extended Data Fig. 11.**
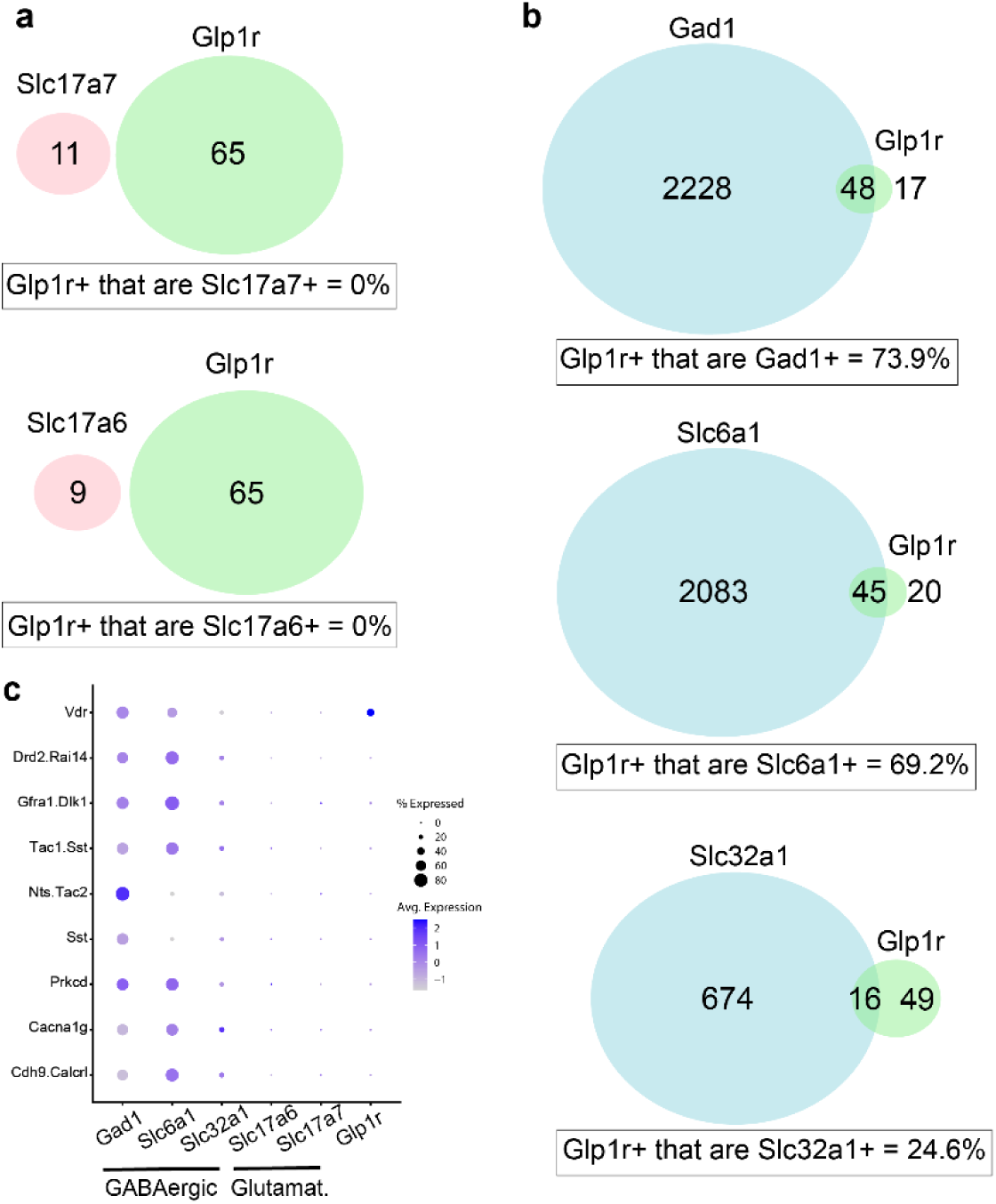
Central amygdala transcriptomic analyses reveal Glp1r neurons are GABAergic ^72^. **a-b**, Coexpression analysis of Glp1r mRNA and glutamatergic markers (Slc17a7 and Slc17a6) and Glp1r and GABAergic markers (Gad1, Slc6a1, and Slc32a1). **c**, Dotplot showing Glp1r is expressed in the Vdr cluster that contains GABAergic markers.

**Extended Data Fig. 12.**
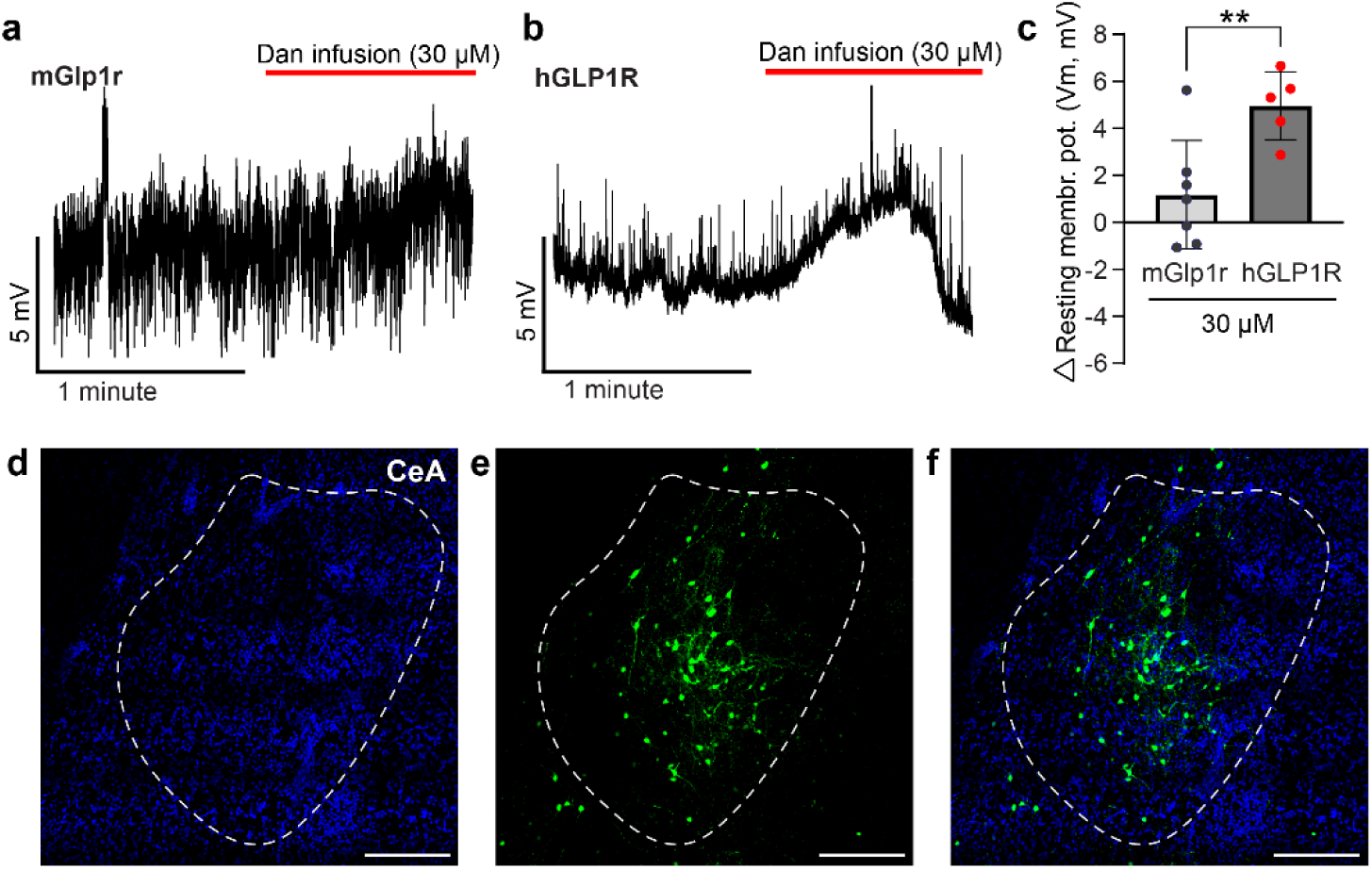
Danuglipron depolarizes the resting membrane potential in hGLP1R-expressing amygdala neurons. Representative traces showing the effect of danuglipron perfusion (30 μM) on the resting membrane potential of **a,** mGlp1r-expressing **b,** hGLP1R-expressing amygdala neurons. **c,** Average depolarization induced by danuglipron in mGlp1r- and hGLP1R-expressing amygdala neurons (*n* = 5-7 cells, Unpaired t-test, \*\**P*<0.01). **d-f,** Localization of eYFP-labeled hGLP1R-expressing neurons in the amygdala. Scale bars = 100 μm.

**Extended Data Fig. 13.**
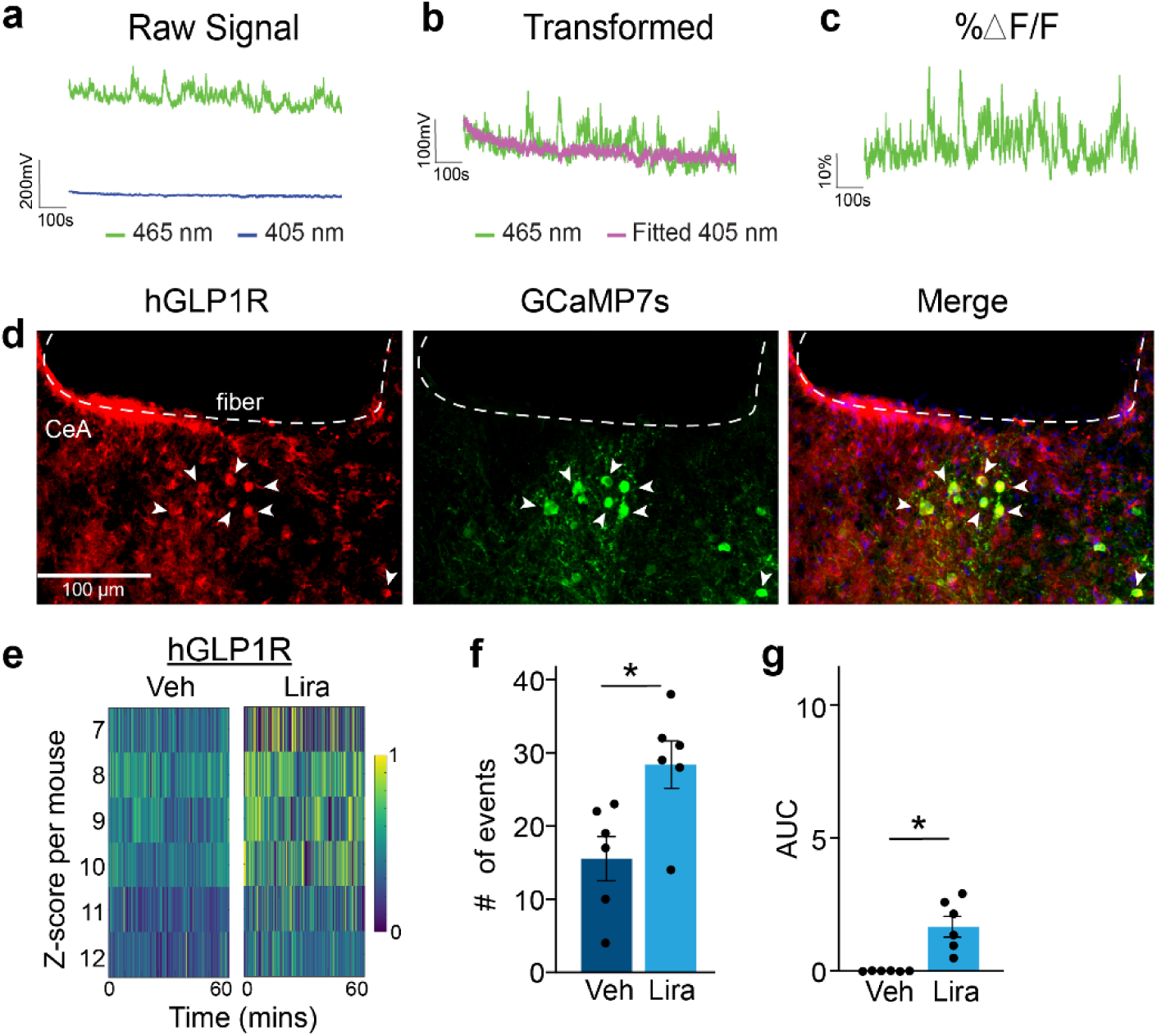
Fiber photometry recordings and validation of CeA^Glp1r^ neurons. **a**, Representative traces of fiber photometry recordings showing calcium-dependent fluorescence (465nm) and calcium-independent, isosbestic fluorescence (405nm) signals. **b**, Fitted 405nm signal (pink) transformed for ΔF/F calculation. **c,** Representative trace of %ΔF/F used for z score analysis. **d**, Representative images of AAV-DIO-GCaMP7s + hGLP1R targeted to the CeA. Validation with hGLP1R antibody (red; left), GCaMP7s (green; middle), and verification of colocalization (right). Scale bars = 100 µm. **e,** Heatmaps of Z-scored neuronal calcium signals for each mouse during 1-hour of fiber photometry recording after vehicle (left) or liraglutide (right) injection in hGLP1R expressing mice. **f**, Number of significant calcium events averaged per condition during 1-hour recording session following vehicle or liraglutide injection (*n* = 6 mice per injection, paired t-test \**P*<0.05). **g**, Area under the curve (AUC) quantification of calcium recordings (*n* = 6 mice per condition, paired t-test, \**P*<0.05). Data are represented as means ± SEM. \**P*<0.05; \*\**P*<0.01; \*\*\**P*<0.001.

**Extended Data Fig. 14.**
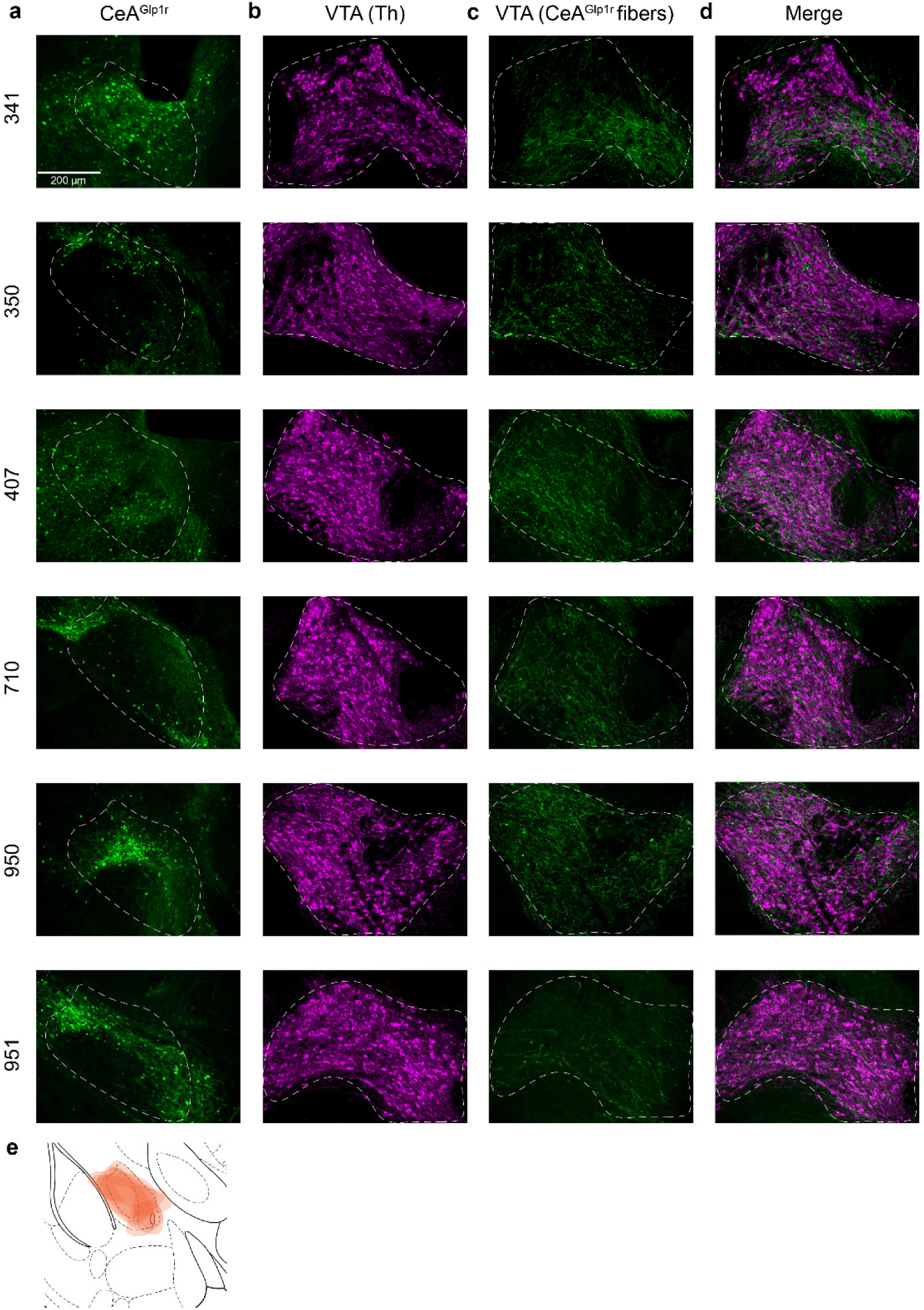
CeA^Glp1r^ neurons innervate the VTA and validation of AAVs targeting CeA and NAc. **a**, Representative images of AAV-DIO-ChR2-eYFP expression targeted to the CeA of Glp1r-Cre mice (green; *n* = 6). **b**, Tyrosine hydroxylase (Th) expression in the VTA (magenta). **c**, Axon fiber projections from CeA^Glp1r^ neurons into the VTA, and **d**, colocalization of Th with CeA^Glp1r^ fibers in the VTA. **e**, Traces of superimposed AAV-DIO-ChR2-eYFP injection targeting per mouse, collapsed on the Allen Brain Atlas figure.

**Extended Figure 15.**
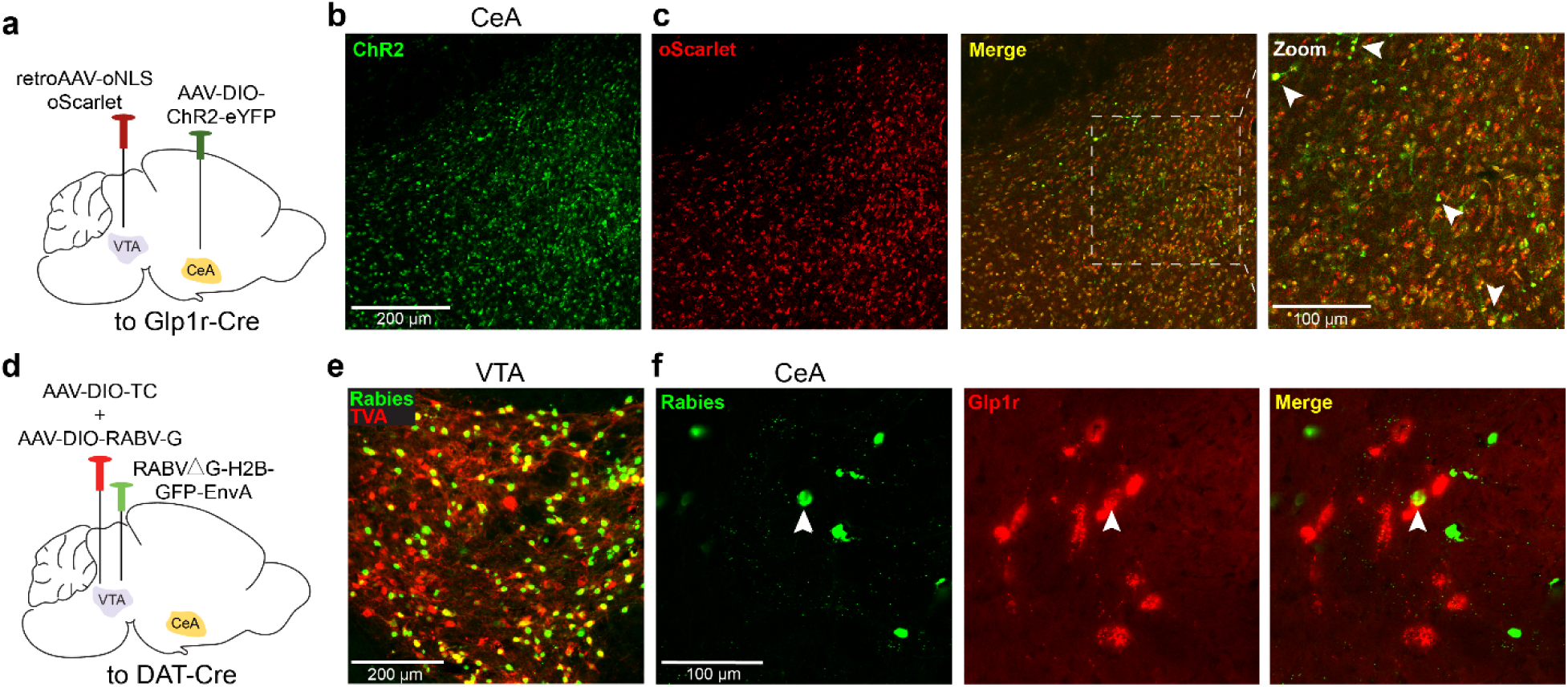
Tracing from CeA to VTA reveals connections between CeA^Glp1r^ and VTA^DAT^ neurons. a, Illustration of AAV-mediated retrograde tracing in Glp1r-Cre mice injected with retroAAV-oNLS-oScarlet in the VTA and Cre-dependent ChR2-eYFP in the CeA. **b,** Representative image showing ChR2-eYFP expression the CeA (scale bar = 200 µm). **c,** Representative images showing Left: oNLS-oScarlet expression in the CeA, Middle: merged image, Right: Magnified view of the merged image; arrowheads mark Glp1r neurons colocalized with oNLS-oScarlet expression (scale bar = 100 µm). **d**, Illustration of cell-type specific, monosynaptic retrograde rabies tracing approach targeting VTA dopamine neurons. **e,** Representative image showing the VTA injection site in Dat-Cre mice (scale bar = 200 µm). **f**, Left: GFP+ input neurons identified in the CeA. Middle: In situ hybridization for Glp1r mRNA in the CeA. Right: merged image demonstrating colocalization (white arrow) of Glp1r mRNA with rabies-derived GFP in the CeA (scale bar = 100 µm).

**Extended Data Fig. 16.**
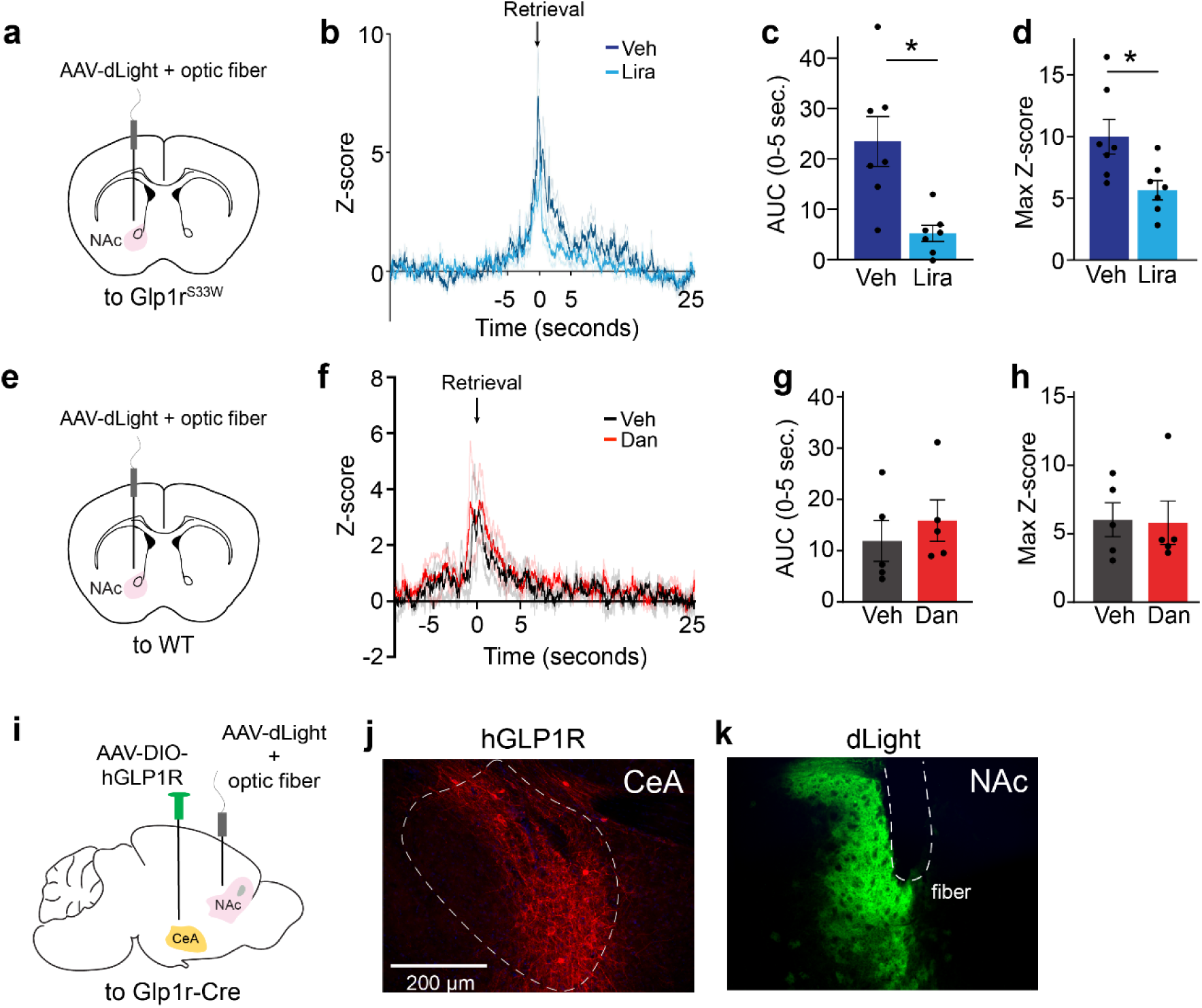
Liraglutide reduces NAc dopamine release in response to HFD while WT mice do not respond to Danuglipron. **a**, Schematic of genetically-encoded dopamine sensor, AAV-dLight1.3b, injection and fiber optic implant in the NAc of Glp1r^S33W^ mice. **b,** Averaged Z-score traces showing dopamine release in the NAc in response to HFD following administration of vehicle or liraglutide in Glp1r^S33W^ mice. Traces are aligned to food retrieval time (t = 0) and averaged across five food trials per mouse. **c,d**, Quantified (**c**) area under the curve (AUC) for Z-scores and (**d**) maximum fluorescence Z-scores within the food retrieval window (*n* = 7, paired t-test, \**P*<0.05). **e**, Schematic of AAV-dLight1.3b injection and fiber optic implant in the NAc of WT mice. **f**, Averaged Z-score traces showing dopamine release in the NAc in response to HFD following administration of vehicle or danuglipron in WT mice. Traces are aligned to food retrieval time (t = 0) and averaged across five food trials per mouse. **g,h**, Quantified (**g**) area under the curve (AUC) for Z-scores and (**h**) maximum fluorescence Z-scores within the food retrieval window (*n* = 5 per injection, paired t-test). **i**, Schematic showing AAV-dLight1.3b injection into the NAc and AAV-DIO-hGLP1R injection into the CeA of Glp1r-Cre mice, with fiber optic implants in the NAc. **j**, Validation of AAV-DIO-hGLP1R targeting the CeA using hGLP1R antibody staining (red). **k**, Representative AAV-dLight expression in the NAc and fiber optic implant placement (green). Scale bars = 200 µm. Data are represented as means ± SEM. \**P*<0.05; \*\**P*<0.01; \*\*\**P*<0.001.

