## Supplemental Table 1 for "A Brain Reward Circuit Inhibited By Next-Generation Weight Loss Drugs"

| Figure | Sample Type (mouse line) | Number of Sample (n/trial) | Behavior Test | Drug or Virus Injection | Description | Statistical Test | Degrees of Freedom (df) | Mean +/- SEM | Statistics | Adjusted p-v |
|---|---|---|---|---|---|---|---|---|---|---|
| Fig. 1b | C57BL6J | 11 / injection | Food Intake | Liraglutide, Danuglipron, Saline, Vehicle | SD intake measured 2 hours after injection | One-way ANOVA w/ Bonferroni Correction | 40 | V1: 0.9 +/- 0.03<br>L: 0.37 +/- 0.05<br>V: 0.84 +/-0.07<br>D: 0.7 +/- 0.08 | F(3, 40) = 15.12<br>p = <0.0001 | D-V: 0.711<br>L-V1: <0.0001 |
| Fig. 1f | Glp1r S33W and WT | 8 WT / 6 S33W | GTT | Liraglutide | Fasted glucose levels up to 2 hours (liraglutide, danuglipron) or 6 hours (orforglipron) after injection | Two-way ANOVA w/ Bonferroni Correction | 84 | | F(13, 84) = 10.62<br>p = <0.0001 | -15: 0.037<br>-1: 0.403<br>15: 0.021<br>30: 0.055<br>60: 0.318<br>90: 0.228<br>120: 0.164 |
| Fig. 1g | | 9 WT / 6 S33W | | Danuglipron | | | 91 | | F(13, 91) = 37.32<br>p = <0.0001 | -15: 0.885<br>-1: 0.415<br>15: <0.0001<br>30: <0.0001<br>60: <0.0001<br>90: 0.008<br>120: 0.026 |
| Fig. 1h | | 5 WT / 5 S33W | | Orforglipron | | | 56 | | F(13, 56) = 35.86 p = <0.0001 | -240: 0.833<br>-1: 0.1401<br>15: <0.0001<br>30: <0.0001<br>60: 0.0004<br>90: 0.0037<br>120: 0.0032 |
| Fig. 1i | | 10-11 / injection | Food Intake | Liraglutide and Vehicle | SD intake measured up to 4 hours after injection for danuglipron and liraglutide or up to 8 hours after orforglipron | | 57 | 1V: 0.43+/-0.05<br>1L: 0.29+/-0.06<br>2V: 0.94+/-0.08<br>2L: 0.45+/-0.08<br>4V: 1.82+/-0.13<br>4L: 0.54+/-0.06 | F(5, 57) = 43.81<br>p = <0.0001 | 1hr = 0.271<br>2hr = 0.0002<br>4hr = <0.0001 |
| Fig. 1j | | 9 / injection | | | | | 48 | 1V: 0.56+/-0.19<br>1L: 0.3+/-0.06<br>2V: 1.08+/-0.22<br>2L: 0.48+/-0.06<br>4V: 2.04+/-0.27<br>4L: 0.7+/-0.06 | F(5, 48) = 14.33<br>p = <0.0001 | 1hr = 0.289<br>2hr = 0.014<br>4hr = <0.0001 |
| Fig. 1k | | 16 / injection | | Danuglipron and Vehicle | | | 90 | 1V: 0.25+/-0.05<br>1D: 0.21+/-0.03<br>2V: 0.71+/-0.06<br>2D: 0.78+/-0.05<br>4V: 1.68+/-0.11<br>4D: 1.74+/-0.10 | F(5, 90) = 84.77<br>p = <0.0001 | 1hr = 0.717<br>2hr = 0.469<br>4hr = 0.587 |
| Fig. 1l | | 15 / injection | | | | | 84 | 1V: 0.34+/-0.06<br>1D: 0.14+/-0.05<br>2V: 0.89+/-0.10<br>2D: 0.43+/-0.07<br>4V: 1.73+/-0.07<br>4D: 1.22+/-0.12 | F(5, 84) = 54.7 p = <0.0001 | 1hr = 0.087<br>2hr = 0.0001<br>4hr = <0.0001 |
| Fig. 1m | | 8 / injection | | Orforglipron and Vehicle | | | 42 | 1V: 0.67+/-0.09<br>1O: 0.7+/-0.05<br>2V: 1.08+/-0.09<br>2O: 1.06+/-0.07<br>4V: 1.5+/-0.12<br>4O: 1.51+/-0.07 | F(5, 42) = 18.14<br>p = <0.0001 | 1hr = 0.9181<br>2hr = 0.9181<br>4hr = 0.9181 |
| Fig. 1n | | 9 / injection | | | | | 48 | 1V: 0.68+/-0.08<br>1O: 0.33+/-0.08<br>2V: 1.1+/-0.09<br>2O: 0.47+/-0.08<br>4V: 1.76+/-0.15<br>4O: 0.86+/-0.16 | F(5, 48) = 21.43<br>p = < 0.0001 | 1hr = 0.0334<br>2hr = 0.0002<br>4hr = <0.0001 |

| | | | | | | | | | | |
|---|---|---|---|---|---|---|---|---|---|---|
| Fig. 1o | Glp1r S33W and WT | 10 / injection | Food Intake | Liraglutide and Vehicle | HFD intake measured up to 4 hours after injection for danuglipron and liraglutide or up to 8 hours after orforglipron | Two-way ANOVA w/ Bonferroni Correction | 54 | 1V: 0.91+/-0.08<br>1L: 0.61+/-0.09<br>2V: 1.17+/-0.13<br>2L: 0.69+/-0.08<br>4V: 1.58+/-0.13<br>4L: 0.8+/-0.09 | F(5, 54) = 12.7<br>p = <0.0001 | 1hr = 0.041<br>2hr = 0.002<br>4hr = <0.0001 |
| Fig. 1p | | 8 / injection | | | | | 42 | 1V: 1.09+/-0.12<br>1L: 0.64+/-0.14<br>2V: 1.32+/-0.15<br>2L: 0.69+/-0.15<br>4V: 1.92+/-0.17<br>4L: 0.76+/-0.19 | F(5, 42) = 10.65<br>p = <0.0001 | 1hr = 0.042<br>2hr = 0.005<br>4hr = <0.0001 |
| Fig. 1q | | 16 / injection | | Danuglipron and Vehicle | | | 90 | 1V: 0.84+/-0.08<br>1D: 0.55+/-0.06<br>2V: 1.09+/-0.10<br>2D: 1.0+/-0.11<br>4V: 1.63+/-0.14<br>4D: 1.49+/-0.13 | F(5, 90) = 14.3<br>p = <0.0001 | 1hr = 0.054<br>2hr = 0.563<br>4hr = 0.342 |
| Fig. 1r | | 14 / injection | | | | | 78 | 1V: 1.0+/-0.08<br>1D: 0.31+/-0.04<br>2V: 1.31+/-0.12<br>2D: 0.57+/-0.06<br>4V: 1.8+/-0.16<br>4D: 1.14+/-0.09 | F(5, 78) = 28.87<br>p = <0.0001 | 1hr = <0.0001 2hr = <0.0001<br>4hr = <0.0001 |
| Fig. 1s | | 8 / injection | | Orforglipron and Vehicle | | | 42 | 1V: 1.15+/-0.06<br>1O: 1.21+/-0.7<br>2V: 1.39+/-0.07<br>2O: 1.5+/-0.08<br>4V: 1.86+/-0.06<br>4O: 2.0+/-0.08 | F(5, 42) = 23.94<br>p = <0.0001 | 1hr = 0.535<br>2hr = 0.267<br>4hr = 0.176 |
| Fig. 1t | | 9 / injection | | | | | 48 | 1V: 1.19+/-0.07<br>1O: 0.69+/-0.09<br>2V:1.5+/-0.10<br>2O: 0.97+/-0.17<br>4V: 1.89+/-0.11<br>4O: 1.29+/-0.20 | F(5, 48) = 9.728<br>p = <0.0001 | 1hr = 0.0111<br>2hr = 0.007<br>4hr = 0.0027 |
| Fig. 1u | | 6 WT / 6 S33W | Oral GTT | Danuglipron | Fasted glucose levels up to 2 hours after danuglipron oral gavage | | 70 | | F(13, 70) = 15.95<br>p = <0.0001 | -15: 0.971<br>-1: 0.265<br>15: 0.005<br>30: <0.0001<br>60: 0.019<br>90: 0.177<br>120: 0.278 |
| Fig. 1v | Glp1r S33W | 8-9 / injection | Food Intake | Danuglipron and Vehicle | Oral administration of Dan or Orfo + HFD Food Intake | Two-way ANOVA w/ Bonferroni Correction | 45 | 1V: 0.98+/-0.12<br>1D: 0.36+/-0.09<br>2V: 1.46+/-0.16<br>2D: 0.57+/-0.11<br>4V: 2.0+/-0.19<br>4D: 1.23+/-0.21 | F(4, 45) = 15.55<br>p <0.0001 | 1hr = 0.0062<br>2hr = 0.0001<br>4hr = 0.0009 |
| Fig. 1w | | | | | | | 41 | IP-V: 1.8+/-0.16<br>IP-D: 1.14+/-0.1<br>O-V: 2.0+/-0.19<br>O-D: 1.2+/-0.63 | F(3, 41) = 6.97<br>p = 0.0007 | IP = 0.0017<br>Oral = 0.005 |
| Fig. 1x | | 8 / injection | | Orforglipron and Vehicle | | | 42 | 1V: 1.33+/-0.06<br>1O: 0.81+/-0.10<br>2V: 1.76+/-0.12<br>2O: 1.06+/-0.09<br>4V: 2.22+/-0.09<br>4O: 1.42+/-0.14 | F(5, 42) = 24.72 p <0.0001 | 1hr = 0.0009<br>2hr = <0.0001<br>4hr = <0.0001 |
| Fig. 1y | | | | | | | 30 | IP-V: 1.89+/-0.11<br>IP-O: 1.29+/-0.2<br>O-V: 2.22+/-0.09<br>O-O: 1.42+/-0.14 | F(3, 30) = 8.508 p = 0.00031 | IP = 0.0056<br>Oral = 0.0008 |
| Fig. 1z | | 10-11 / injection | Weight Loss | Orforglipron and Vehicle | Chronic orfo or saline injections to overweight mice on HFD for > 8 week | Paired t-test | 399 | Day 21:<br>O: -8.45+/-1.1<br>V: -2.0+/-1.2 | t(399) = - 16.1<br>p <0.0001 | |

| | | | | | | | | | |
|---|---|---|---|---|---|---|---|---|---|
| Fig. 2i | Glp1r S33W | veh/dan: 9, veh/orfo:10, veh/lira: 9, veh/licl: 6 | Home cage behavior | Danuglipron, Orforglipron, Liraglutide, LiCl, Vehicle | Frequency of head entries using TTL beam break in 2 hour home cage recording (ZT 12-14) | Paired t-test (on raw data) | veh/dan: 8, veh/orfo:9, veh/lira: 8,veh/licl: 5 | V (lira): 139.22222 +/- 15.180193 lira: 59.22222 +/- 13.099807 V: 113.44444 +/- 13.631164 D: 58.88889 +/- 12.393372 V(licl): 82.66667 +/- 11.101551 licl: 39.83333 +/- 8.392126 V(orfo): 123.00000 +/- 15.783254 O: 86.30000 +/- 15.182629 | Lira-V: t(8) = -6.3762 p <0.0001 D-V: t(8) = -3.0464 p = 0.0159 Licl-V: t(5) = -3.6663 p=0.0145 O-V: t(9) = -3.9432 p=0.00339 |
| Fig. 2j | | | | | Proportion of time spent performing behavior in 2 hour home cage recording (ZT 12-14) | GLMM with beta regression, cloglog link due to right-skewed distribution, and a random intercept for mouse ID (paired design) | | V (lira): 0.179 +/- 0.0277 lira: 0.0937 +/- 0.0167 V: 0.162 +/- 0.0206 D: 0.0919 +/- 0.0138 V(licl): 0.0825 +/- 0.0105 licl: 0.0515 +/- 0.00665 V(orfo): 0.142 +/- 0.0237 O: 0.120 +/- 0.0255 | Lira-V: β = -0.717 p <0.001 D-V: β = -0.612 p <0.001 Licl-V: β = -0.479 p<0.001 O-V: β = -0.183 p=0.3903 |
| Fig. 2k | | | | | Proportion of time spent performing behavior in 2 hour home cage recording (ZT 12-14) | GLMM with beta regression, cloglog link due to right-skewed distribution, and a random intercept for mouse ID (paired design) | | V (lira): 0.0279 +/- 0.00321 lira: 0.00786 +/- 0.00173 V: 0.0268 +/- 0.00486 D: 0.00985 +/- 0.00347 V(licl): 0.0163 +/- 0.00386 licl: 0.0130 +/- 0.00310 V(orfo): 0.0298 +/- 0.00280 O: 0.0284 +/- 0.00241 | Lira-V: β = -1.38 p <0.001 D-V: β = -1.28 p <0.001 Licl-V: β = -0.225 p=0.1348 O-V: β = -0.045 p=0.6291 |
| Fig. 2l | | | | | Proportion of time spent performing behavior in 2 hour home cage recording (ZT 12-14) | GLMM with beta regression, logit link due to approximately symmetric distribution, and a random intercept for mouse ID (paired design) | | V(lira): 0.472 +/- 0.0407 lira: 0.401 +/- 0.0578 V: 0.530 +/- 0.0546 D: 0.389 +/- 0.0829 V(licl): 0.640 +/- 0.0770 licl: 0.334 +/- 0.116 V(orfo): 0.561 +/- 0.0661 O: 0.602 +/- 0.0492 | Lira-V: β = -0.317 p =0.1735 D-V: β = -0.611 p =0.1092 Licl-V: β = -1.36 p=0.0064 O-V: β = 0.18 p=0.2886 |
| Fig. 2m | | | | | Proportion of time spent performing behavior in 2 hour home cage recording (ZT 12-14) | GLMM with beta regression, logit link due to approximately symmetric distribution, and a random intercept for mouse ID (paired design) | | V(lira): 0.299 +/- 0.0332 lira: 0.433 +/- 0.0607 V: 0.229 +/- 0.0411 D: 0.427 +/- 0.0863 V(licl): 0.248 +/- 0.0753 licl: 0.557 +/- 0.123 V(orfo): 0.222 +/- 0.0526 O: 0.209 +/- 0.0389 | Lira-V: β =0.533 p =0.0543 D-V: β = 0.78 p = 0.0432 Licl-V: β = 1.29 p=0.0162 O-V: β = -0.0773 p=0.7003 |

| | | | | | | | | | | |
|---|---|---|---|---|---|---|---|---|---|---|
| Fig. 2u | Glp1r S33W | licl: 6 | Home cage behavior | LiCl | Spearman correlations between behavioral features and Euclidean distance to the LiCl group centroid in UMAP space. Positive Spearman ρ values indicate that higher values of a behavioral feature are associated with greater dissimilarity from LiCl-treated mice, whereas negative ρ values indicate that higher values are associated with greater similarity to the LiCl group. | Spearman's ρ. P-values adjusted for multiple comparisons using the Benjamini–Hochberg false discovery rate (FDR) method | | | move to drink: ρ = 0.628, p <0.001<br>drink to move: ρ = 0.612, p <0.001<br>shelter to move: ρ = 0.602, p <0.001<br>move to food motivated: ρ = 0.593, p <0.001<br># bouts drink: ρ = 0.575, p = 0.0000, p <0.001<br>drink sec: ρ = 0.559, p <0.001<br>avg bout length shelter: ρ = -0.555, p <0.001<br>food motivated to move: ρ = 0.534, p <0.001<br># bouts food motivated: ρ = 0.530, p <0.001<br>move to shelter: ρ = 0.515, p <0.001<br>Food hooper head entries: ρ | move to drink: p <0.001<br>drink to move: p <0.001<br>shelter to move: p <0.001<br>move to food motivated: p <0.001<br># bouts drink: p <0.001<br>drink sec: p <0.001<br>avg bout length shelter: p <0.001<br>food motivated to move: p = 0.0010<br># bouts food motivated: p = 0.0010<br>move to shelter: p = 0.0015<br>Food hooper head entries: p = 0.0026<br>max bout length shelter: p = 0.0031<br>rest/groom in shelter p = 0.0051 |
| Fig. 3e | Glp1r S33W and WT | 3-4 / injection | Image Analysis | Vehicle, Danuglipron, Orforglipron, Liraglutide | Quantified Fos+ neurons divided by area in brain regions of interest: (e) DMH, (f) NTS, (g) AP, and (h) CEA | Two-way ANOVA w/ Bonferroni Correction | 23 | V-W: 1.5+/-0.1<br>V-S: 1.0+/-0.19<br>D-W: 1.32+/-0.39<br>D-S: 1.65+/-0.09<br>O-W: 1.3+/-0.13<br>O-S: 1.12+/-0.12<br>L-W: 1.59+/-0.06<br>L-S: 1.37+/-0.19 | F(7,23) = 1.30<br>p = 0.2943 | Veh = 0.110<br>Dan = 0.211<br>Orfo = 0.572<br>Lira = 0.409 |
| Fig. 3f | | | | | | | 21 | V-W: 0.48+/-0.11 V-S: 0.52+/-0.05 D-W: 0.34+/-0.10<br>D-S: 2.23+/-0.47<br>O-W: 0.39+/-0.1<br>O-S: 1.96+/-0.04<br>L-W: 1.71+/-0.13<br>L-S: 1.24+/-0.2 | F(7, 21) = 19.49<br>p <0.0001 | Veh = 0.884<br>Dan = <0.0001<br>Orfo = <0.0001<br>Lira = 0.0568 |
| Fig. 3g | | | | | | | 21 | V-W: 0.43+/-0.2<br>V-S: 0.2+/-0.09<br>D-W: 0.71+/-0.4<br>D-S: 4.02+/-0.61<br>O-W: 0.17+/-0.03<br>O-S: 2.4+/-0.38<br>L-W: 2.95+/-1.0<br>L-S: 2.58+/-0.62 | F(7, 21) = 6.724<br>p = 0.0003 | Veh = 0.787<br>Dan = 0.0005<br>Orfo = 0.0077<br>Lira = 0.6223 |
| Fig. 3h | | | | | | | 23 | V-W: 0.51+/-0.16 V-S: 0.44+/-0.08 D-W: 0.28+/-0.05<br>D-S: 0.84+/-0.07<br>O-W: 0.3+/-0.07<br>O-S: 0.95+/-0.12<br>L-W: 1.56+/-0.32<br>L-S: 1.25+/-0.18 | F(7,23) = 8.662<br>p < 0.0001 | Veh = 0.7633<br>Dan = 0.0203<br>Orfo = 0.0076<br>Lira = 0.1698 |
| Fig. 3j | Glp1r S33W | | | | Ratio of NTS / AP Fos+ cells in Glp1r S33W mice compared among injections | Kruskal-Wallis test | 2 | D: 0.54+/-0.04<br>O: 0.88+/-0.12<br>L: 0.53+/-0.09 | Chi-squared = 6.167<br>p = 0.045 | |
| Fig. 3m | | 3-6 / injection | | Danuglipron and Orforglipron | Comparison between Fos activation via IP and oral delivery of danuglipron and orforglipron in the NTS | Two-way ANOVA w/ Bonferroni Correction | 13 | Dan<br>IP: 2.2+/-0.47<br>O: 1.8+/-0.2<br>Orfo<br>IP: 1.96+/-0.04<br>O: 1.4+/-0.08 | F(3,13) = 2.384<br>p = 0.1164 | Dan = 0.1784<br>Orfo = 0.0817 |
| Fig. 3n | | | | | Comparison between Fos activation via IP and oral delivery of danuglipron and orforglipron in the AP | | 13 | Dan<br>IP: 4.02+/-0.61<br>O: 1.14+/-0.3<br>Orfo<br>IP: 2.4+/-0.38<br>O: 1.8+/-0.31 | F(3,13) = 9.647<br>p = 0.00128 | Dan = 0.0001<br>Orfo = 0.3304 |

| | | | | | | | | | | |
|---|---|---|---|---|---|---|---|---|---|---|
| Fig. 4b | Glp1r-cre | 6 / injection | Food Intake | Danuglipron and Vehicle AAV-DIO-mCherry or AAV-DIO-hGLP1R | mCherry (b) or hGLP1R (c) to basomedial hypothalamus + SD | Two-way ANOVA w/ Bonferroni Correction | 30 | 1V: 0.33+/-0.07<br>1D: 0.4+/-0.07<br>2V: 0.6+/-0.11<br>2D: 0.73+/-0.09<br>4V: 1.45+/-0.11<br>4D: 1.63+/-0.19 | $F(5, 30)$ = 23.82 p = <0.0001 | 1hr = 0.679<br>2hr = 0.410<br>4hr = 0.26 |
| Fig. 4c | | 10 / injeciton | | | | | 54 | 1V: 0.43+/-0.06<br>1D: 0.14+/-0.05<br>2V: 0.86+/-0.10<br>2D: 0.38+/-0.08<br>4V: 1.86+/-0.13<br>4D: 1.02+/-0.19 | $F(5, 54)$ = 32.12<br>p = <0.0001 | 1hr = 0.066<br>2hr = 0.0031<br>4hr = <0.0001 |
| Fig. 4g | | 6 / injection | | | mCherry (g) or hGLP1R (h) to basomedial hypothalamus + HFD | | 30 | 1V: 0.55+/-0.10<br>1D: 0.55+/-0.08<br>2V: 0.97+/-0.14<br>2D: 0.72+/-0.10<br>4V: 1.82+/-0.15<br>4D: 1.48+/-0.11 | $F(5, 30)$ = 20.89<br>p = <0.0001 | 1hr = 1.00<br>2hr = 0.135<br>4hr = 0.050 |
| Fig. 4h | | 10 / injeciton | | | | | 54 | 1V: 0.65+/-0.12<br>1D: 0.66+/-0.19<br>2V: 1.15+/-0.14<br>2D: 0.99+/-0.09<br>4V: 1.8+/-0.09<br>4D: 1.84+/-0.12 | $F(5, 54)$ = 20.93<br>p = <0.0001 | 1hr = 0.952<br>2hr = 0.334<br>4hr = 0.808 |
| Fig. 4d | | 7 / injection | | | hGLP1R to DMH + SD (d) or + HFD (i) | | 36 | 1V: 0.47+/-0.06<br>1D: 0.17+/-0.04<br>2V: 0.83+/-0.13<br>2D: 0.5+/-0.09.<br>4V: 1.7+/-0.21<br>4D: 1.16+/-0.20 | $F(5, 36)$ = 16.01<br>p = <0.0001 | 1hr = 0.134<br>2hr = 0.102<br>4hr = 0.009 |
| Fig. 4i | | | | | | | 36 | 1V: 0.79+/-0.22<br>1D: 0.57+/-0.11<br>2V: 1.1+/-0.22<br>2D: 0.99+/-0.12<br>4V: 1.73+/-0.23<br>4D: 1.51+/-0.16 | $F(5,36)$ = 5.65<br>p = 0.0006 | 1hr = 0.416<br>2hr = 0.663<br>4hr = 0.416 |
| Fig. 4e | Glp1r-cre | 6 / injection | Food Intake | Danuglipron and Vehicle AAV-DIO-mCherry or AAV-DIO-hGLP1R | hGLP1R to NTS/AP + SD (e) or + HFD (j) | Two-way ANOVA w/ Bonferroni Correction | 30 | 1V: 0.32+/-0.05<br>1D: 0.20+/-0.07<br>2V: 0.80+/-0.12<br>2D: 0.58+/-0.12<br>4V: 2.0+/-0.29<br>4D: 1.28+/-0.24 | $F(5, 30)$ = 15.36<br>p = <0.0001 | 1hr = 0.636<br>2hr = 0.382<br>4hr = 0.006 |
| Fig. 4j | | | | | | | 30 | 1V: 0.97+/-0.07<br>1D: 0.63+/-0.08<br>2V: 1.32+/-0.11<br>2D: 0.82+/-0.07<br>4V: 2.18+/-0.12<br>4D: 1.48+/-0.16 | $F(5, 30)$ = 27.82<br>p = <0.0001 | 1hr = 0.035<br>2hr = 0.002<br>4hr = 0.0001 |
| Fig. 4f | | 9 / injection | | | hGLP1R to CeA + SD (f) or + HFD (k) | | 48 | 1V: 0.26+/-0.06<br>1D: 0.19+/-0.07<br>2V: 0.58+/-0.07<br>2D: 0.58+/-0.11<br>4V: 1.47+/-0.08<br>4D: 1.43+/-0.19 | $F(5, 48)$ = 29.26<br>p = <0.0001 | 1hr = 0.654<br>2hr = 1.00<br>4hr = 0.823 |
| Fig. 4k | | | | | | | 47 | 1V: 0.96+/-0.10<br>1D: 0.54+/-0.08<br>2V: 1.13+/-0.13<br>2D: 0.83+/-0.08<br>4V: 1.82+/-0.15<br>4D: 1.42+/-0.15 | $F(5, 47)$ = 15.22<br>p = <0.0001 | 1hr = 0.016<br>2hr = 0.074<br>4hr = 0.018 |
| Fig. 4o | Glp1r-cre | 3-4 / injection | Image Analysis | Danuglipron and Vehicle AAV-DIO-hGLP1R | hGLP1R and mGlp1r to CeA | Two-way ANOVA with Tukey's HSD Correction | 10 | mGlp1r<br>V: 11.9+/-1.6<br>D: 7.2+/-1.7<br>hGLP1R<br>V: 7.3+/-0.6<br>D: 20+/-0.8 | $F(3,10)$ = 22.97<br>p <0.0001 | hGLP1R<br>D-V = 0.0002<br>Dan<br>mGlp1r-hGLP1r = 0.0001 |

| | | | | | | | | | | |
|---|---|---|---|---|---|---|---|---|---|---|
| Fig. 4s | | | | | AUC quantification of calcium recordings after DAN injection in mGlp1r vs hGLP1R-expressing mice | Unpaired t-test | n/a | mGlp1r-Dan: -0.050 +/- 0.261 hGLP1R-Dan: 3.304 +/- 1.400 | p = 0.0043 | |
| Fig. 4t | Glp1r-Cre | 6 / injection | in vivo calcium imaging | AAV-DIO-GCaMP7s+hGLP1R or mGlp1r to CeA | Number of calcium events for each group and injection conditions | Two-way ANOVA w/ Bonferroni Correction | 20 | mGlp1r-Veh: -20 +/- 3.109 mGlp1r-Dan: 16.833 +/- 1.905 hGLP1R - Veh: 17.833 +/- 2.725 hGLP1R - Dan: 31.5 +/- 3.481 | F (1, 20) = 8.629 p = 0.0081 | mGlp1r Veh vs hGLP1R Veh : >0.9999 mGlp1r Veh vs mGlp1r Dan : >0.9999 mGlp1r Veh vs hGLP1R Dan : 0.0610 mGlp1r Dan vs hGLP1R Veh : >0.9999 hGLP1R Veh vs hGLP1R Dan : 0.0182 hGLP1R Dan vs mGlp1r Dan: 0.0103 |
| Fig 4v | Glp1r-Cre | 5 eYFP / 5 ChR2 | Opto food intake | AAV-DIO-eYFP or AAV-DIO-ChR2-eYFP to CeA & fiber optic implant to CeA | 1-hour SD intake measured with stim ON or OFF | Two-way ANOVA w/ Bonferroni Correction | 32 | eYFP - 0 Hz: 1.150 +/- 0.047 eYFP - 5 Hz: 1.300 +/- 0.124 eYFP - 10 Hz: 0.972 +/- 0.135 eYFP - 20 Hz: 1.256 +/- 0.170 ChR2 - 0 Hz: 1.202 +/- 0.132 ChR2 - 5 Hz: 1.132 +/- 0.142 ChR2 - 10 Hz: 0.914 +/- 0.188 ChR2 - 20 Hz: 1.134 +/- 0.161 | F (3, 32) = 0.2222 p = 0.8803 | eYFP vs ChR2 (0 Hz): >0.999 eYFP vs ChR2 (5 Hz): >0.999 eYFP vs ChR2 (10 Hz): 0.0906 eYFP vs ChR2 (20 Hz): 0.0043 |
| Fig. 4w | | | | | 1-hour HFD intake measured with stim ON or OFF | Two-way ANOVA w/ Bonferroni Correction | 32 | eYFP - 0 Hz: 0.716 +/- 0.064 eYFP - 5 Hz: 0.688 +/- 0.045 eYFP - 10 Hz: 0.744 +/- 0.056 eYFP - 20 Hz: 0.746 +/- 0.044 ChR2 - 0 Hz: 0.750 +/- 0.080 ChR2 - 5 Hz: 0.698 +/- 0.093 ChR2 - 10 Hz: 0.500 +/- 0.108 ChR2 - 20 Hz: 0.380 +/- 0.059 | F (3,32) = 3.682 p = 0.220 | eYFP vs ChR2 (0 Hz): 07988 eYFP vs ChR2 (5 Hz): 0.4124 eYFP vs ChR2 (10 Hz): 0.7762 eYFP vs ChR2 (20 Hz): 0.5507 |
| Fig. 4y | Glp1r flox/flox | 6 GFP / 6 Cre | Food Intake | AAV-GFP or AAV-Cre to CeA | SD or HFD intake measurements 4 hours after injection of liraglutide | Welch's t-test | SD: 6 HFD: 9 | SD GFP: 0.33+/-0.1 Cre: 0.45+/-0.2 HFD GFP: 0.45+/-0.1 Cre: 0.72+/-0.1 | SD: t(5.7) = 0.55 p = 0.6028 HFD: t(9.4) = 2.4 p = 0.03738 | |
| Fig. 5c | Gcg-Cre | 4 mCherry / 6 ChrimsonR | Opto Food Intake | AAV-DIO-mcherry or AAV-DIO-ChrimsonR to NTS & Bilateral fiber optic implants to CeA | 1-hour SD intake measured with stim ON or OFF | Two-way ANOVA w/ Tukey's HSD | 16 | mCherry-OFF: 0.543 +/- 0.078 mCherry-ON: 0.658 =/- 0.071 ChrimsonR-OFF: 0.568 +/- 0.064 ChrimsonR-ON: 0.523 =/- 0.075 | F (1, 16) = 1.156 p = 0.2982 | OFF:mCherry vs. OFF:ChrimsonR: 0.9946 OFF:mCherry vs. ON:mCherry : 0.7529 OFF:mCherry vs. ON:ChrimsonR: 0.9978 OFF:ChrimsonR vs. ON:mCherry : 0.8312 OFF:ChrimsonR vs. ON:ChrimsonR: 0.9628 ON:mCherry vs. ON:ChrimsonR: 0.5908 |

| | | | | | | | | | | |
|---|---|---|---|---|---|---|---|---|---|---|
| Fig. 5d | Gcg-Cre | 4 mCherry / 6 ChrimsonR | Opto Food Intake | AAV-DIO-mcherry or AAV-DIO-ChrimsonR to NTS & Bilateral fiber optic implants to CeA | 1-hour HFD intake measured with stim ON or OFF | Two-way ANOVA w/ Tukey's HSD | 16 | mCherry-OFF: 0.745 +/- 0.031 mCherry-ON: 0.764 +/- 0.038 ChrimsonR-OFF: 0.570 +/- 0.052 ChrimsonR-ON: 0.290 +/- 0.066 | F (1, 16) = 7.062 p=0.0172 | OFF:mCherry vs. OFF:ChrimsonR: 0.1651 OFF:mCherry vs. ON:mCherry : 0.9963 OFF:mCherry vs. ON:ChrimsonR: 0.0002 OFF:ChrimsonR vs. ON:mCherry : 0.1097 OFF:ChrimsonR vs. ON:ChrimsonR: 0.0058 ON:mCherry vs. ON:ChrimsonR: 0.0001 |
| Fig. 5g | Glp1r-Cre | 4 mCherry / 5 ChrimsonR | Opto Food Intake | AAV-DIO-mcherry or AAV-DIO-ChrimsonR to CeA & Unilateral fiber optic implant to VTA | 1-hour SD intake measured with stim ON or OFF | Two-way ANOVA w/ Tukey's HSD | 14 | mCherry-OFF: 1.205 +/- 0.167 mCherry-ON: 1.060 +/- 0.250 ChrimsonR-OFF: 0.850 +/- 0.094 ChrimsonR-ON: 0.984 +/- 0.200 | F (1, 14) = 0.5884 p=0.4558 | OFF:mCherry vs. OFF:ChrimsonR: 0.5309 OFF:mCherry vs. ON:mCherry : 0.9491 OFF:mCherry vs. ON:ChrimsonR: 0.8253 OFF:ChrimsonR vs. ON:mCherry : 0.8457 OFF:ChrimsonR vs. ON:ChrimsonR: 0.9443 ON:mCherry vs. ON:ChrimsonR: 0.9906 |
| Fig. 5h | Glp1r-Cre | 5 mCherry / 5 ChrimsonR | Opto Food Intake | AAV-DIO-mcherry or AAV-DIO-ChrimsonR to CeA & Unilateral fiber optic implant to VTA | 1-hour HFD intake measured with stim ON or OFF | Two-way ANOVA w/ Tukey's HSD | 16 | mCherry-OFF: 0.666 +/- 0.098 mCherry-ON: 0.692 +/- 0.089 ChrimsonR-OFF: 0.676 +/- 0.061 ChrimsonR-ON: 0.354 +/- 0.055 | F (1, 16) = 4.990 p=0.0401 | OFF:mCherry vs. OFF:ChrimsonR: 0.9997 OFF:mCherry vs. ON:mCherry : 0.9952 OFF:mCherry vs. ON:ChrimsonR: 0.0528 OFF:ChrimsonR vs. ON:mCherry : 0.9989 OFF:ChrimsonR vs. ON:ChrimsonR: 0.0444 ON:mCherry vs. ON:ChrimsonR: 0.0334 |
| Fig. 5k | Glp1r S33W | 7-9 / injection | in vivo dopamine imaging | AAV-dLight1.3b to NAc | Measure dopamine release AUC into NAc in reponse to HFD after Veh or Dan injection at retrieval (k) and max dopamine Z-score (l) | Paired t-test | 8 | V: 17.0+/-3.34 D: 10.6+/-2.24 | t(8) = -2.49 p = 0.0373 | |
| Fig. 5l | | | | | | | | V: 10.7+/-1.0 D: 8.14+/-0.78 | t(8) = -2.46 p = 0.0391 | |
| Fig. 5n | | | | | Measure dopamine release AUC into NAc in reponse to HFD after Veh or Orfo injection at retrieval (n) and max dopamine Z-score (o) | | 6 | V: 23.47 +/- 4.961 O: 4.38+/-0.97 | t(6) = -2.92 p = 0.0156 | |
| Fig. 5o | | | | | | | | V: 10.08 +/- 1.406 O: 6.39+/-0.3 | t(6) = -2.794 p = 0.0312 | |
| Fig. 5r | Glp1r-cre | 7-8 / injection | | AAV-DIO-hGLP1R to CeA and AAV-dLight1.3b to Nac | Measure dopamine release AUC into NAc in reponse to HFD after Veh or Dan injection at retrieval (k) and max dopamine (l) | | 7 | V: 14.9+/-3.15 D: 7.53+/-1.45 | t(7) = -2.47 p = 0.0425 | |
| Fig. 5s | | | | | | | | V: 10.3+/-1.47 D: 5.87+/-0.76 | t(7) = -2.61 p = 0.0351 | |
| Fig. 5u | | | | | Measure dopamine release AUC into NAc in reponse to HFD after Veh or Orfo injection at retrieval (n) and max dopamine (o) | | 6 | V: 23.47 +/- 4.961 O: 4.85+/-2.03 | t(6) = -2.46 p = 0.0493 | |
| Fig. 5v | | | | | | | | V: 6.67+/-1.04 O: 4.48+/-0.82 | t(6) = -3.4 p = 0.0145 | |

| | | | | | | | | | | |
|---|---|---|---|---|---|---|---|---|---|---|
| Ext. Data Fig. 1b | Glp1r S33W and WT | 11 WT / 10 S33W | Metabolic Profilin | NA | Measure diurnal rhythms and sex differences between genotypes of respiratory exchange ratio (RER) and energy expenditure (EE) Meaure baseline bodyweight and sex difference between genotypes | Two-way ANOVA w/ Bonferroni Correction | 42 | WT<br>Dark: 0.87+/-0.02 Light: 0.80+/-0.02<br>S33W<br>Dark: 0.85+/-0.02 Light: 0.79+/-0.03 | F(3, 42) = 3.84<br>p = 0.0162 | Dark: 0.488<br>Light: 0.828 |
| Ext. Data Fig. 1c | | | | | | Welch's t-test | 17.1 | WT: 0.84+/-0.02<br>S33W:0.82+/-0.02 | t(17) = -0.45<br>p = 0.656 | |
| Ext. Data Fig. 1d | | 5 WT F / 4 S33W F<br>7 WT M / 7 WT S33W | | | | Two-way ANOVA w/ Bonferroni Correction | 19 | WT<br>Fem: 0.84+/-0.03 Male: 0.83+/-0.02<br>S33W<br>Fem: 0.81+/-0.06 Male: 0.83+/-0.02 | F(3, 19) = 0.171<br>p = 0.9149 | Fem: 0.496<br>Male: 0.980 |
| Ext. Data Fig. 1f | | 11 WT / 10 S33W | | | | | 42 | WT<br>Dark: 0.49+/-0.02 Light: 0.36+/-0.01<br>S33W<br>Dark: 0.51+/-0.02 Light: 0.39+/-0.01 | F(3, 42) = 22.64<br>p = <0.0001 | Dark: 0.326<br>Light: 0.093 |
| Ext. Data Fig. 1g | | | | | | Welch's t-test | 20.9 | WT: 0.42+/-0.02<br>S33W:0.45+/-0.01 | t(21) = 1.41<br>p = 0.172 | |
| Ext. Data Fig. 1h | | 5 WT F / 4 S33W F<br>7 WT M / 7 WT S33W | | | | Two-way ANOVA w/ Bonferroni Correction | 19 | WT<br>Fem: 0.39+/-0.02 Male: 0.45+/-0.02<br>S33W<br>Fem: 0.44+/-0.03 Male: 0.46+/-0.02 | F(3,19) = 2.36<br>p = 0.1037 | Fem: 0.100<br>Male: 0.692 |
| Ext. Data Fig. 1i | | 18 WT / 19 S33W | | | | Welch's t-test | 28.1 | WT: 25.1+/-1.01<br>S33W: 24.9+/-0.61 | t(28) = -0.21<br>p = 0.834 | |
| Ext. Data Fig. 1j | | 8 WT F / 9 S33W F<br>10 WT M / 10 S33W M | | | | Two-way ANOVA w/ Bonferroni Correction | 33 | WT<br>Fem: 22.4+/-1.24 Male: 27.3+/-1.13<br>S33W<br>Fem: 23.2+/-0.61 Male: 26.4+/-0.73 | F(3, 33) = 6.35<br>p = 0.0016 | Fem: 0.583<br>Male: 0.492 |
| Ext. Data Fig. 2a | Glp1r S33W | 8 - 15 / injection | Food Intake | Danuglipron and Vehicle | Vehicle, 3mg/kg, 10 mg/kg, 30 mg/kg Danuglipron dose response 2-hours post injection measuring SD intake | One-way ANOVA with Tukey's HSD | 43 | V: 0.89 +/-0.1<br>D3: 0.87+/-0.08<br>D10: 0.99+/-0.07<br>D30: 0.43+/-0.07 | F(3, 43) = 8.963<br>p < 0.0001 | V-D30:0.0007<br>D3-D30: 0.0067<br>D10-D30: 0.00063 |
| Ext. Data Fig. 2b | | 6 - 8 / injection | | Danuglipron, Orforglipron, Liraglutide, Vehicle | SD intake following a 16 hour overnight fast + (b) Lira, (c) Dan, (d) Orfo or control injection prior to refeeding | Two-way ANOVA w/ Bonferroni Correction | 30 | 1V: 1.1+/-0.09<br>2V: 1.6+/-0.07<br>4V: 2.6+/-0.19<br>1L: 0.58+/-0.07<br>2L: 0.63+/-0.04<br>4L: 0.77+/-0.06 | F(5,30) = 63.24<br>p < 0.0001 | 1hr: 0.0014<br>2hr: < 0.0001<br>4hr: < 0.0001 |
| Ext. Data Fig. 2c | | | | | | | 30 | 1V: 0.78+/-0.11<br>2V: 1.3+/-0.2<br>4V: 2.2+/-0.26<br>1D: 0.15+/-0.11<br>2D: 0.35+/-0.1<br>4D: 1.3+/-0.16 | F(5,30) = 21.2 p < 0.0001 | 1hr: 0.011<br>2hr: 0.0002<br>4hr: 0.0006 |
| Ext. Data Fig. 2d | | | | | | | 42 | 1V: 1.5+/-0.13<br>2V: 1.96+/-0.17<br>4V: 2.9+/-0.18<br>1O: 0.9+/-0.09<br>2O: 1.1+/-0.13<br>4O: 1.7+/-0.17 | F(5, 42) = 22.74<br>p < 0.0001 | 1hr: 0.0055<br>2hr: 0.0002<br>4hr: <0.0001 |

| | | | | | | | | | | |
|---|---|---|---|---|---|---|---|---|---|---|
| Ext. Data Fig. 2e | Glp1r S33W and WT | 6 WT / 6 S33W | Food Intake | Liraglutide, Danuglipron, Orforglipron, Vehicle | 24 hour SD intake after GLP1RA or vehicle/saline | Two-way ANOVA w/ Bonferroni Correction | 20 | WT<br>V: 4.4+/-0.11<br>L: 2.1+/-0.32<br>S33W<br>V: 4.9+/-0.26<br>L: 2.7+/-0.38 | F(3, 20) = 21.21<br>p = <0.0001 | WT: <0.0001<br>S33W: <0.0001 |
| Ext. Data Fig. 2f | | | | | | | | WT<br>V:4.8+/-0.35<br>D: 5.2+/-0.28<br>S33W<br>V: 4.8+/-0.20<br>D: 4.8+/-0.22 | F(3, 20) = 0.557<br>p = 0.649 | WT: 0.286<br>S33W: 0.965 |
| Ext. Data Fig. 2g | | | | | | | | WT<br>V:4.98+/-0.29<br>O: 5.9+/-0.2<br>S33W<br>V: 5.4+/-0.23<br>O: 4.2+/-0.38 | F(3, 20) = 6.38<br>p = 0.0033 | WT: 0.032<br>S33W: 0.008 |
| Ext. Data Fig. 3b | Glp1r S33W | 5 - 8 / injection | CTA | Danuglipron, Orforglipron, Liraglutide, LiCl, Vehicle | 24 hour conditioned taste avoidance assay | One-way ANOVA with Tukey's HSD | 38 | Veh: 0.8+/-0.01<br>LiCl: 0.37+/-0.09<br>Lira: 0.39+/-0.06<br>Dan: 0.58+/-0.08 oDan: 0.67+/-0.1<br>Orfo: 0.55+/-0.11<br>oOrfo: 0.53+/-0.13 | F(6, 38) = 3.264<br>p = 0.01093 | V-LiCl:0.01969<br>V-Lira: 0.01254 |
| Ext. Data Fig. 3c | | 8 / injection | Anxiety Testing | Danuglipron, Orforglipron, Vehicle | Elevated plus maze on (c-e) danuglipron or (h-j) orforglipron and open field test on (f,g) danuglipron or (k,l) orforglipron | Welch's t-test | 14 | V: 11.5+/-3.6<br>D: 9.5+/-3.2 | t(14) = -0.4056<br>p = 0.6912 | |
| Ext. Data Fig. 3d | | | | | | | 14 | V: 4.6+/-1.7<br>D: 3.3+/-1.4 | t(14) = -0.6294<br>p = 0.5396 | |
| Ext. Data Fig. 3e | | | | | | | 14 | V: 798+/-112<br>D: 590+/-98 | t(14) = -1.3996<br>p = 0.1838 | |
| Ext. Data Fig. 3f | | | | | | | 14 | V: 16.2+/-3.5<br>D: 11.7+/-3.0 | t(14) = -0.9811<br>p = 0.3437 | |
| Ext. Data Fig. 3g | | | | | | | 13 | V: 1656+/-174<br>D: 1504+/-242 | t(13) = -0.5127<br>p = 0.617 | |
| Ext. Data Fig. 3h | | | | | | | 12 | V: 16.2+/-2.0<br>O: 18.6+/-3.1 | t(12) = 0.668<br>p = 0.5167 | |
| Ext. Data Fig. 3i | | | | | | | 11 | V: 4.3+/-0.5<br>O: 4.2+/-1.0 | t(11) = -0.1420<br>p = 0.8897 | |
| Ext. Data Fig. 3j | | | | | | | 14 | V: 907+/-90<br>O: 912+/-75 | t(14) = 0.0471<br>p = 0.9631 | |
| Ext. Data Fig. 3k | | | | | | | 13 | V: 16.4+/-3.4<br>O: 19.7+/-2.6 | t(13) = 0.7697<br>p = 0.4552 | |
| Ext. Data Fig. 3l | | | | | | | 14 | V: 2217+/-120<br>O: 2056+/-101 | t(14) = -1.026<br>p = 0.3229 | |

| | | | | | | | | | | |
|---|---|---|---|---|---|---|---|---|---|---|
| Ext. Data Fig. 7a | Glp1r S33W | veh/dan: 9, veh/orfo:10, veh/lira: 9, veh/licl: 6 | Home cage behavior | Danuglipron, Orforglipron, Liraglutide, LiCl, Vehicle | Frequency of water bottle lciks (sensor) in 2 hour home cage recording (ZT 12-14) | Paired t-test (on raw data) | veh/dan: 8, veh/orfo:9, veh/lira: 8, veh/licl: 5 | V(lira): 996.4444 +/- 221.89826<br>lira: 330.1111 +/- 128.99639<br>V: 1300.0000 +/- 216.01607<br>D: 350.1111 +/- 156.69392<br>V(licl): 671.1667 +/- 235.48537<br>licl: 388.3333 +/- 113.85507<br>V(orfo): 1190.1000 +/- 234.12463<br>O: 1040.4000 +/- 206.63044 | Lira-V: t(8) = -2.91<br>p =0.0196<br>D-V: t(8) = -4.85<br>p = 0.00127<br>Licl-V: t(5) = -1.83<br>p=0.127<br>O-V: t(9) = -1.46<br>p=0.177 | |
| Ext. Data Fig. 7b | | | | | Proportion of time spent performing behavior in 2 hour home cage recording (ZT 12-14) | GLMM with beta regression, logit link due to approximately symmetric distribution, and a random intercept for mouse ID (paired design) | | V(lira): 0.445+/- 0.0416<br>lira: 0.373 +/- 0.0540<br>V: 0.510 +/- 0.0542<br>D: 0.372 +/- 0.0817<br>V(licl): 0.611+/- 0.0811<br>licl: 0.309+/- 0.116<br>V(orfo): 0.536+/- 0.0680<br>O: 0.579+/- 0.0506 | Lira-V: β = -0.334<br>p=0.2282<br>D-V: β = -0.62<br>p =0.1060<br>Licl-V: β = -1.35<br>p=0.0148<br>O-V: β = 0.161<br>p=0.6036 | |
| Ext. Data Fig. 7d | | | | | Proportion of time spent performing behavior in 2 hour home cage recording (ZT 12-14) | GLMM with beta regression, cloglog link due to right-skewed distribution, and a random intercept for mouse ID (paired design) | | V(lira): 0.0230 +/-0.00543<br>lira: 0.0646 +/- 0.0136<br>V: 0.0516 +/- 0.0188<br>D: 0.0820 +/- 0.0287<br>V(licl): 0.0130+/- 0.00423<br>licl: 0.0446+/- 0.0121<br>V(orfo): 0.0452+/- 0.0165<br>O: 0.0408+/- 0.0112 | Lira-V: β = 0.933<br>p=0.0014<br>D-V: β = 0.425<br>p =0.1378<br>Licl-V: β = 1.14<br>p<0.001<br>O-V: β = -0.00812<br>p=0.9722 | |
| Ext. Data Fig. 7e | | | | | Frequency of full wheel cycles (sensor) in 2 hour home cage recording (ZT 12-14) | Paired t-test (on raw data) | | V(lira): 4145.444 +/- 829.3747<br>lira: 2631.111 +/- 506.4680<br>V: 4642.000 +/- 705.9874<br>D: 2676.778 +/- 839.8130<br>V(licl): 5529.833+/- 962.5975<br>licl: 2234.500+/- 988.4082<br>V(orfo): 4928.100+/- 1179.4251<br>O: 4871.400+/- 921.0457 | Lira-V: t(8) = -1.6603<br>p =0.1354<br>D-V: t(8) = -0.52368<br>p = 0.6147<br>Licl-V: t(5) = -3.1214<br>p=0.02621<br>O-V: t(7) = -0.10525<br>p=0.9191 | |
| Ext. Data Fig. 7f | | | | | Total distance travelled using SLEAP upper body/neck keypoint in 2 hour home cage recording (ZT 12-14) | Paired t-test | | V(lira): 4145.444 +/- 829.3747<br>lira: 2631.111 +/- 506.4680<br>V: 4642.000 +/- 705.9874<br>D: 2676.778 +/- 839.8130<br>V(licl): 5529.833+/- 962.5975<br>licl: 2234.500+/- 988.4082<br>V(orfo): 4928.100+/- 1179.4251<br>O: 4871.400+/- 921.0457 | Lira-V: t(8) = -2.6004<br>p =0.0316<br>D-V: t(8) = -1.9083<br>p = 0.09278<br>Licl-V: t(5) = -3.8093<br>p=0.01251<br>O-V: t(9) = -0.19629<br>p=0.8487 | |
| Ext. Data Fig. 7g | | dan: 9, orfo:10, lira: 9, licl: 6, fed: 10 | | Danuglipron, Orforglipron, Liraglutide, LiCl, Fed | Proportion of time spent performing behavior in 2 hour home cage recording (ZT 12-14) | GLMM with beta regression, cloglog link due to right-skewed distribution. Post hoc comparisons between the Fed group and all other groups were performed with Holm-adjusted p-values. | | lira: 0.0937 +/- 0.0167<br>D: 0.0919 +/- 0.0138<br>licl: 0.0515 +/- 0.00665<br>O: 0.120 +/- 0.0255<br>F: 0.0337 +/- 0.0104 | Lira-fed: β = 1.209<br>p<0.001<br>D-fed: β = 1.259<br>p<0.001<br>Licl-fed: β = 0.831<br>p=0.025656<br>O-fed: β = 1.458<br>p<0.001 | Lira-fed: β = 1.209<br>p<0.001<br>D-fed: β = 1.259<br>p<0.001<br>Licl-fed: β = 0.831<br>p=0.0257<br>O-fed: β = 1.458<br>p<0.001 |

| | | | | | | | | | | |
|---|---|---|---|---|---|---|---|---|---|---|
| Ext. Data Fig. 7h | Glp1r S33W | dan: 9, orfo:10, lira: 9, licl: 6, fed: 10 | Home cage behavior | Danuglipron, Orforglipron, Liraglutide, LiCl, Fed | Proportion of time spent performing behavior in 2 hour home cage recording (ZT 12-14) | GLMM with beta regression, cloglog link due to right-skewed distribution. Post hoc comparisons between the Fed group and all other groups were performed with Holm-adjusted p-values. | | lira: 0.00786 +/- 0.00173<br>D: 0.00985 +/- 0.00347<br>licl: 0.0130 +/- 0.00310<br>O: 0.0284 +/- 0.00241<br>F: 0.0106 +/- 0.00291 | Lira-fed: β = -0.320 p=0.3222<br>D-fed: β = -0.689 p=0.0459<br>Licl-fed: β = 0.273 p=0.3975<br>O-fed: β = 1.024 p<0.001 | Lira-fed: β = -0.320 p=0.6443<br>D-fed: β = -0.689 p=0.1377<br>Licl-fed: β = 0.273 p=0.6443<br>O-fed: β = 1.024 p<0.001 |
| Ext. Data Fig. 7l | | | | | Frequency of head entries using TTL beam break in 2 hour home cage recording (ZT 12-14) | One-way ANOVA. Post hoc comparisons between the Fed group and all other groups were performed with bonferroni-adjusted p-values. | dan: 8, orfo:9, lira: 8, licl: 5, fed: 9 | lira: 59.22222 +/- 13.099807<br>D: 58.88889 +/- 12.393372<br>licl: 39.83333 +/- 8.392126<br>O: 86.30000 +/- 15.182629<br>F: 22.70000 +/- 5.176979 | F(4, 39) = 4.384 p=0.00506 | Lira: p=0.1207<br>D: p=0.1265<br>Licl: p=1.0000<br>O: p=0.0010 |
| Ext. Data Fig. 7m | | | | | Frequency of water bottle lciks (sensor) in 2 hour home cage recording (ZT 12-14) | One-way ANOVA. Post hoc comparisons between the Fed group and all other groups were performed with bonferroni-adjusted p-values. | | lira: 330.1111 +/- 128.99639<br>D: 350.1111 +/- 156.69392<br>licl: 388.3333 +/- 113.85507<br>O: 1040.4000 +/- 206.63044<br>F: 287.9000 +/- 67.55943 | F(4, 39) = 5.005 p=0.00236 | Lira: p=1.0000<br>D: p=1.0000<br>Licl: p=1.0000<br>O: p=0.0019 |
| Ext. Data Fig. 8a | | | | | Sum of move–groom transition probabilities (both directions) | One-way ANOVA. Post hoc comparisons between the Fed group and all other groups were performed with bonferroni-adjusted p-values. | | lira: 0.562 +/- 0.0555<br>D: 0.587 +/- 0.107<br>licl: 0.410 +/- 0.128<br>O: 0.560 +/- 0.0455<br>F: 0.842 +/- 0.107 | F(4, 39) = 2.94 p=0.0322 | Lira: p=0.1077<br>D: p=0.1727<br>Licl: p=0.0124<br>O: p=0.0904 |
| Ext. Data Fig. 8b | | | | | Sum of move–drink transition probabilities (both directions) | One-way ANOVA. Post hoc comparisons between the Fed group and all other groups were performed with bonferroni-adjusted p-values. | | lira: 0.237 +/- 0.0416<br>D: 0.284 +/- 0.0683<br>licl: 0.249 +/- 0.0597<br>O: 0.438 +/- 0.0447<br>F: 0.531 +/- 0.113 | F(4, 39) = 3.19 p=0.0233 | Lira: p=0.0234<br>D: p=0.0757<br>Licl: p=0.0693<br>O: p=1.0000 |
| Ext. Data Fig. 8d | | | | | Sum of move–shelter transition probabilities (both directions) | One-way ANOVA. Post hoc comparisons between the Fed group and all other groups were performed with bonferroni-adjusted p-values. | | lira: 0.390 +/- 0.0664<br>D: 0.412 +/- 0.0595<br>licl: 0.272 +/- 0.0815<br>O: 0.543 +/- 0.0557<br>F: 0.893 +/- 0.133 | F(4, 39) = 7.42 p<0.001 | Lira: p<0.001<br>D: p=0.0011<br>Licl: p<0.001<br>O: p=0.0187 |
| Ext. Data Fig. 8h | | | | | average bout length (seconds) of food motivated behaviors | One-way ANOVA. Post hoc comparisons between the Fed group and all other groups were performed with bonferroni-adjusted p-values. | | lira: 17.2 +/- 2.35<br>D: 16.9 +/- 1.49<br>licl: 15.0 +/- 2.35<br>O: 16.6 +/- 2.33<br>F: 8.35 +/- 1.46 | F(4, 39) = 3.75 p=0.0113 | Lira: p=0.0104<br>D: p=0.0145<br>Licl: p=0.1568<br>O: p=0.0154 |

| | | | | | | | | | | |
|---|---|---|---|---|---|---|---|---|---|---|
| Ext. Data Fig. 8l | Glp1r S33W | dan: 9, orfo:10, lira: 9, licl: 6, fed: 10 | Home cage behavior | Danuglipron, Orforglipron, Liraglutide, LiCl, Fed | average bout length (seconds) of rest/groom in shelter behaviors | One-way ANOVA. Post hoc comparisons between the Fed group and all other groups were performed with bonferroni-adjusted p-values. | dan: 8, orfo:9, lira: 8, licl: 5, fed: 9 | lira: 39.9 +/- 9.31<br>D: 45.3 +/- 12.5<br>licl: 38.2 +/- 6.95<br>O: 19.5 +/- 3.55<br>F: 16.7 +/- 2.97 | F(4, 39) = 2.90 p=0.0340 | Lira: p=0.1419<br>D: p=0.0419<br>Licl: p=0.3219<br>O: p=1.0000 |
| Ext. Data Fig. 8m | | | | | maximum bout length (seconds) of drinking behaviors | One-way ANOVA. Post hoc comparisons between the Fed group and all other groups were performed with bonferroni-adjusted p-values. | | lira: 10.7 +/- 2.17<br>D: 11.6 +/- 2.48<br>licl: 15.0 +/- 3.23<br>O: 20.5 +/- 2.02<br>F: 12.0 +/- 1.92 | F(4, 39) = 3.38 p= 0.0181 | Lira: p=1.0000<br>D: p=1.0000<br>Licl: p=1.0000<br>O: p=0.0306 |
| Ext. Data Fig. 8n | | | | | maximum bout length (seconds) of food motivated behaviors | One-way ANOVA. Post hoc comparisons between the Fed group and all other groups were performed with bonferroni-adjusted p-values. | | lira: 107.0 +/- 21.6<br>D: 77.9 +/- 12.1<br>licl: 79.7 +/- 4.18<br>O: 99.9 +/- 15.4<br>F: 36.9 +/- 8.79 | F(4, 39) = 3.86 p=0.00976 | Lira: p=0.00469<br>D: p= 0.1844<br>Licl: p= 0.2523<br>O: p=0.0095 |
| Ext. Data Fig. 8q | | | | | maximum bout length (seconds) of move/explore in shelter behaviors | One-way ANOVA. Post hoc comparisons between the Fed group and all other groups were performed with bonferroni-adjusted p-values. | | lira: 5.00 +/- 0.747<br>D: 4.73 +/- 0.289<br>licl: 4.47 +/- 0.330<br>O: 5.38 +/- 0.247<br>F: 7.42 +/- 0.622 | F(4, 39) = 5.54 p=0.00124 | Lira: p=0.0051<br>D: p=0.0016<br>Licl: p=0.0022<br>O: p= 0.0184 |
| Ext. Data Fig. 8s | | | | | total number (frequency) of drink bouts | One-way ANOVA. Post hoc comparisons between the Fed group and all other groups were performed with bonferroni-adjusted p-values. | | lira: 8 +/- 1.88<br>D: 9.33 +/- 3.33<br>licl: 11 +/- 1.88<br>O: 23.8 +/- 2.92<br>F: 9.7 +/- 1.87 | F(4, 39) = 7.13 p<0.001 | Lira: p=1.0000<br>D: p=1.0000<br>Licl: p=1.0000<br>O: p<0.001 |
| Ext. Data Fig. 8t | | | | | total number (frequency) of food motivated behavior bouts | One-way ANOVA. Post hoc comparisons between the Fed group and all other groups were performed with bonferroni-adjusted p-values. | | lira: 34.4 +/- 5.37<br>D: 35.9 +/- 5.21<br>licl: 23.5 +/- 2.26<br>O: 46.6 +/- 6.74<br>F: 21 +/- 7.05 | F(4, 39) = 3.00 p=0.0298 | Lira: p=0.4546<br>D: p=0.3234<br>Licl: p=1.0000<br>O: p=0.0120 |
| Ext. Data Fig. 8w | | | | | total number (frequency) of move/explore in shelter behavior bouts | One-way ANOVA. Post hoc comparisons between the Fed group and all other groups were performed with bonferroni-adjusted p-values. | | lira: 65.6 +/- 14.7<br>D: 44.3 +/- 6.14<br>licl: 65.5 +/- 13.4<br>O: 54.8 +/- 7.95<br>F: 101.0 +/- 13.2 | F(4, 39) = 3.87 p= 0.00970 | Lira: p=0.1116<br>D: p=0.0031<br>Licl: p=0.1946<br>O: p=0.0162 |
| Ext. Data Fig. 8x | | | | | total number (frequency) of rest/groom in shelter behavior bouts | One-way ANOVA. Post hoc comparisons between the Fed group and all other groups were performed with bonferroni-adjusted p-values. | | lira: 94.3 +/- 17.3<br>D: 80.0 +/- 12.1<br>licl: 107.0 +/- 15.9<br>O: 71.8 +/- 9.31<br>F: 166.0 +/- 16.9 | F(4, 39) = 7.35 p<0.001 | Lira: p=0.0035<br>D: p<0.001<br>Licl: p=0.0486<br>O: p<0.001 |

| | | | | | | | | | | |
|---|---|---|---|---|---|---|---|---|---|---|
| Ext. Data Fig. 10b | Glp1r-Cre | 4 Glp1r-S33W / 8 mGlp1r / 9 hGLP1R | Food Intake | Danuglipron and Vehicle AAV-DIO-mGLP1R or -mGLP1R-S33W or AAV-DIO-hGLP1R | Normailzed food intake after 4 hours for SD (b) or HFD (c) with danuglipron or vehicle | Two-way ANOVA w/ Bonferroni Correction | 36 | hGlp1r:<br>V: 1.0+/-0.05<br>D: 0.98+/-0.13<br>Glp1r-S33W:<br>V: 1.0+/-0.15<br>D: 0.94+/-0.04<br>mGlp1r:<br>V: 1.0+/-0.09<br>D: 0.96+/-0.12 | F(5, 36) = 0.05<br>p = 0.998 | hGlp1r:<br>p = 0.865<br>Glp1r-S33W:<br>p = 0.752<br>mGlp1r:<br>p = 0.772 |
| Ext. Data Fig. 10c | | | | | | | 36 | hGlp1r:<br>V: 1.0+/-0.08<br>D: 0.78+/-0.08<br>Glp1r-S33W:<br>V: 1.0+/-0.16<br>D: 0.55+/-0.07<br>mGlp1r:<br>V: 1.0+/-0.04<br>D: 0.90+/-0.07 | F(5, 36) = 3.47<br>p = 0.01167 | hGlp1r:<br>p = 0.0386<br>Glp1r-S33W:<br>p = 0.0061<br>mGlp1r:<br>p = 0.3381 |
| Ext. Data Fig. 12c | Glp1r-Cre | 7 cells for mGlp1r<br>5 cells for hGLP1R | Electrophysiolo gy | Danuglipron & AAV-DIO-mGlp1r+eYFP or AAV-DIO-hGLP1R+eYFP | Measuring changes to resting membrane potential between mGlp1r vs hGLP1R-expressing cells during DAN infusion | Unpaired t-test | 10 | mGlp1r ΔRMP:<br>1.170 +/- 0.8740<br>hGLP1R ΔRMP:<br>4.962 +/- 0.6442 | p = 0.0091 | |
| Ext. Data Fig. 13f | Glp1r-Cre | 6 / injection | in vivo calcium imaging | AAV-DIO-GCaMP7s+hGLP1R | Number of calcium events after Vehicle or Lira injection | Paired t-test | 5 | hGLP1R - Veh:<br>15.83 +/- 3.027<br>hGLP1R - Lira:<br>28.67 +/- 3.263 | p = 0.0312 | |
| Ext. data Fig. 13g | | | | | AUC quantification of calcium recordings after Vehicle or Lira injection in hGLP1R-expressing mice | Paired t-test | 5 | hGLP1R - Veh: -0.0005 +/- 0.005<br>hGLP1R - Lira:<br>1.742 +/- 0.3908 | p = 0.0312 | |
| Ext. Data Fig. 16c | Glp1r S33W | 7 / injection | in vivo dopamine imaging | AAV-dLight1.3b to NAc | Measure dopamine release AUC into NAc in reponse to HFD after Veh or Lira injection at retrieval (c) and max dopamine Z-score (d) | Paired t-test | 6 | V: 23.47 +/- 4.961<br>L: 5.202 +/- 1.603 | p =0.0156 | |
| Ext. Data Fig. 16d | | | | | | | 6 | V: 10.08 +/- 1.406<br>L: 5.821 +/- 0.7804 | p = 0.0238 | |
| Ext. Data Fig. 16g | WT | 5 / injection | in vivo dopamine imaging | AAV-dLight1.3b to NAc | Measure dopamine release AUC into NAc in reponse to HFD after Veh or Dan injection at retrieval (g) and max dopamine Z-score (h) | Paired t-test | 4 | V: 11.92 +/- 3.961<br>D: 15.88 +/- 4.022 | p =0.652 | |
| Ext. Data Fig. 16h | | | | | | | 4 | V: 6.301 +/- 1.232<br>D: 5.801+/- 1.596 | p > 0.9999 | |
